# Engineered Feedback Employing Natural Hypoxia-Responsive Factors Enhances Synthetic Hypoxia Biosensors

**DOI:** 10.1101/2024.09.28.615614

**Authors:** Kathleen S. Dreyer, Patrick S. Donahue, Jonathan D. Boucher, Katherine M. Chambers, Marya Y. Ornelas, Hailey I. Edelstein, Benjamin D. Leibowitz, Katherine J. Zhu, Kate E. Dray, Joseph J. Muldoon, Joshua N. Leonard

**Affiliations:** Department of Chemical and Biological Engineering, Northwestern University, Evanston, Illinois 60208, United States; Medical Scientist Training Program, Northwestern University Feinberg School of Medicine, Chicago, Illinois 60611, United States; Interdisciplinary Biological Sciences Program, Northwestern University, Evanston, Illinois 60208, United States; Department of Biomedical Engineering, Northwestern University, Evanston, Illinois 60208, United States; Center for Synthetic Biology, Northwestern University, Evanston, Illinois 60208, United States; Chemistry of Life Processes Institute, Northwestern University, Evanston, Illinois 60208, United States; Member, Robert H. Lurie Comprehensive Cancer Center, Northwestern University, Evanston, Illinois 60208, United States

## Abstract

DNA-based hypoxia biosensors conditionally express a gene of interest when a cell is in a state of inadequate oxygen supply, which is a feature of several acute and chronic diseases. These biosensors can be deployed in engineered cells to study or treat disease. Although the central mediators of hypoxia responsiveness have been characterized, the dynamics of this response are generally less understood, and there is no general approach to modulate hypoxia biosensors to tune their performance to meet application-specific needs. To address the need for high-performing hypoxia biosensors, we investigated strategies to enhance biosensor performance by identifying minimal promoter choices and positive feedback circuits that both achieved low background and amplified hypoxia-induced gene expression. To generate insight into the mechanisms by which feedback drives differential performance, we developed an explanatory mathematical model. Our analysis suggests a previously unreported dual regulatory mechanism that was necessary to explain the full set of experimental observations and that provides new insights into regulatory dynamics in chronic hypoxia. This study exemplifies the potential of using synthetic gene circuits to perturb natural systems in a manner that uniquely enables the elucidation of novel facets of natural regulation.

## INTRODUCTION

Hypoxia, a condition in which a tissue lacks sufficient oxygen, is present in a variety of processes including normal embryogenesis, acute pathologies (e.g., heart attacks, stroke, and wound healing), and chronic diseases (e.g., diabetes, peripheral arterial disease, lung disease, and cancer) (1–6). Although healthy tissues have multiple mechanisms to counteract hypoxia, including altering cellular metabolism and increasing blood flow through arterial dilatation, diseased tissues such as tumors lack the ability to mitigate oxygen restriction (7). Likewise, after an acute or chronic interruption in blood flow resulting in ischemia, cells must either reestablish the flow or adapt to hypoxia (8,9). These pathologies have a large impact on human health, with the aforementioned diseases responsible for 58% of US deaths in 2019 and many more people living with these chronic diseases (10). Consequently, the ability to study and target hypoxia for therapeutic intervention has important implications for human health (11,12).

The cellular response to hypoxia is coordinated by two hypoxia-inducible factors (HIFs), HIF1α and HIF2α, each of which is stabilized and accumulates as oxygen levels decrease (**Figure 1A**) (13,14). These transcription factors (TFs) each heterodimerize with HIF1β, bind to hypoxia response elements (HREs), and induce the expression of genes that promote adaptation to and resolution of hypoxia (13–19). This signaling system forms the basis of DNA-based hypoxia biosensors (HBSs). By placing HREs upstream of a minimal promoter, a downstream gene of interest can be conditionally expressed when the cell experiences hypoxic conditions (20–22). HBSs have enabled *in vivo* imaging of the response to hypoxia in mice (21–23). Additionally, several cell therapy approaches have been proposed that employ HBSs. Among these are hypoxia-sensing viral vectors that drive expression of therapeutic genes for treatment of ischemic disease (24–26), cancer (24), anemia (27), and choroidal neovascularization (28,29), and hypoxia-sensing CAR-T cells for the treatment of solid tumors (30).

**Figure 1.**
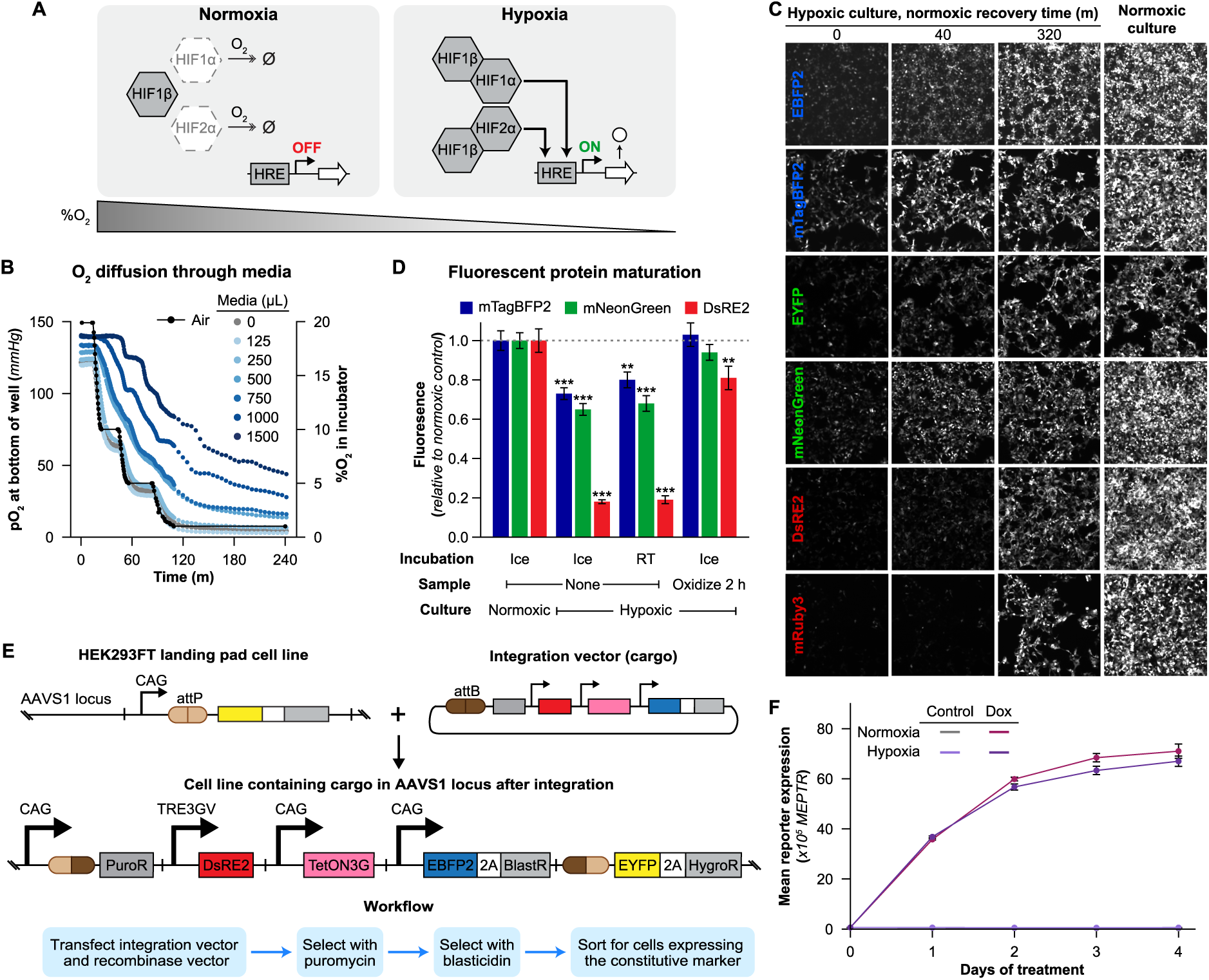
Validation of methods for studying hypoxia. (**A**) Schematic of the cellular response to hypoxia, where HIF1α and HIF2α are destabilized in the presence of oxygen, while in the absence of oxygen they bind to HIF1β and activate genes containing hypoxia response elements (HRE). (**B**) Profile of O_2_ pressure measured at the bottom of cell-culture wells under various conditions of interest, using an electronic sensor (n=1). (**C**) Micrographs illustrating oxidation of fluorophores in cells cultured in hypoxia compared to fluorophores in cells cultured in normoxia. (**D**) Comparison of methods for fluorophore oxidation. Following culture in normoxia or hypoxia, cells were harvested for immediate analysis or allowed to oxidize in normoxia for 2 hours prior to harvest for analysis. Fluorescence was then evaluated by flow cytometry. Significant differences relative to the normoxic condition are identified for each fluorescent protein (1-way ANOVA for each fluorescent protein, ** P < 0.01, *** P < 0.001). (**E**) Schematic depicting the Landing Pad (LP) system implemented in the AAVS1 locus of the HEK293FT cell line and Bxb1 recombinase-mediated integration of a vector (in this case, the vector cargo comprises a module conferring doxycycline-inducible DsRE2). (**F**) Validation that core methods yield comparable control circuit output under normoxia and hypoxia conditions. HEK293FT-LP cells with the integration shown in **E** were cultured as described with samples collected each day for flow cytometry. (for **D**, **F** n=3, error bars represent SEM). Outcomes from ANOVAs and Tukey’s HSD tests for **D** are in **Supplementary Note 5.**

HBS output can be modulated in several ways. Established strategies include adding elements upstream to increase the response (20–22) or modulating the choice of minimal promoter to influence the response magnitude (ON state) and gene expression in normoxic conditions (OFF state, or leak) (31). There also exist gaps in our understanding for applying HBSs. First, the dynamics of the response to hypoxia are generally not characterized, and no approach for modulating dynamics is available. Second, it is unclear how well HBSs that rely exclusively on the endogenous response will perform when deployed in cancerous cells in which dysregulation of the hypoxia response is common (6,32). To address these limitations, we sought to better understand potential strategies to tune and enhance the performance of hypoxia biosensors.

In this study, we developed synthetic genetic circuits constructed using natural hypoxia regulatory factors to interrogate, characterize, and modulate the cellular response to hypoxia. We identified new HBS designs that exhibit improved performance compared to prior ones, including faster biosensor output after exposure to hypoxia. To explain the behaviors observed across our ensemble of experimental HBS designs, we developed an explanatory dynamic model using ordinary differential equations. This combined approach, which integrated experimental perturbation of the hypoxia response using synthetic genetic circuits and development of a model to formally pose and test mechanistic hypotheses, led us to propose a novel potential mode of the natural hypoxia regulation system. These technologies and insights will benefit the use of hypoxia biosensors for applications in fundamental research, biotechnology, and medicine.

## MATERIAL AND METHODS

### Plasmids, cloning, and DNA sources

All plasmid maps are available in **Supplementary Data 1**. Constructs were initially characterized in the pPD005 plasmid backbone, which is a version of pcDNA3.1(+), modified as described previously (33). All HBS components were transferred into the mMoClo system (34) with previously described modifications (33). Coding sequences are generally flanked by NheI and NotI restriction sites. Promoter regions are generally flanked by a BglII or a MluI site on the 5’ end and an NheI site on the 3’ end.

A previously characterized HBS (20) was synthesized as overlapping oligonucleotides, as were the minimal promoters YB_TATA and CMV_min (31). The minimal promoter SV40_min, the EF1α and TRE3GV promoters, the genes encoding the stable HIF1α mutant, TetON3G, BlastR, EBFP2, and EYFP; and the CHS4 insulator were sourced from plasmids from Addgene and Clontech as described in **Supplementary Table 11** (31,35–39). The genes for mTagBFP2 (40), mNeonGreen (41), mRuby3 (42), miRFP670 (43,44), and miRFP720 (45) were synthesized as codon-optimized Gene Strings by Thermo Fisher. The gene for DsRed-Express2 (DsRE2) was obtained by site directed mutagenesis of pDsRed2-N1, and an internal BpiI restriction site in the coding region was ablated by making a sense mutation with site directed mutagenesis. Plasmids from the mMoClo system were a gift from Ron Weiss (**Supplementary Table 11**) (34). Plasmids for Cas9-mediated targeted integration were obtained from Addgene and modified in this study by moving AdE4orf6 from pPD782 into pPD720 to make pPD783 (**Supplementary Table 11**).

### Assembly of mMoClo Integration Vectors via Golden Gate Assembly

The mMoClo integration vectors were assembled through a BpiI-mediated Golden Gate reaction. Each 20 μL reaction comprised 2 µL 10x T4 ligase buffer, 2 µL 10x BSA (1 mg/mL stock), 0.8 µL BpiI-FD, 0.8 µL T4 DNA Ligase (400 U/µL stock), 20 fmol integration vector backbone (pPD630 or pPD1178), and 40 fmol of each transcription unit and linker plasmid to be inserted. The reaction was incubated at 37°C for 15 min, then subjected to 55 iterations of thermocycling (37°C for 5 min, 16°C for 3 min, repeat), followed by 37°C for 15 min, 50°C for 5 min, 80°C for 10 min to terminate the reactions; then the mixture was cooled to room temperature (approximately 20–25°C, throughout) and then optionally held at 4°C, and placed on ice prior to immediate transformation into bacteria. Combinations of plasmids and the resultant vectors are described in **Supplementary Table 2**.

### Plasmid preparation

TOP10 E. coli were grown overnight in 100 mL of LB with an appropriate selective antibiotic. The following morning, cells were pelleted at 3000 x g for 10 min and then resuspended in 4 mL of a solution of 25 mM Tris pH 8.0, 10 mM EDTA, and 15% sucrose. Cells were lysed for 15 min by addition of 8 mL of a solution of 0.2 M NaOH and 1% SDS, followed by neutralization with 5 mL of 3 M sodium acetate (pH 5.2). Precipitate was pelleted by centrifugation at 9000 x g for 20 min. Supernatant was decanted and treated with RNAse A for 1 h at 37°C. 5 mL of phenol chloroform was added, and the solution was mixed and then centrifuged at 7500 x g for 20 min. The aqueous layer was removed and subjected to another round of phenol chloroform extraction with 7 mL of phenol chloroform. The aqueous layer was subjected to an isopropanol precipitation (41% final volume isopropanol, 10 min at room temperature, 9000 x g for 20 min), and the pellet was briefly dried and resuspended in 420 µL of water. The DNA mixture was incubated on ice for at least 12 h in a solution of 6.5% PEG 20,000 and 0.4 M NaCl (1 mL final volume). DNA was precipitated with centrifugation at maximum speed for 20 min. The pellet was washed once with ethanol, dried for several h at 37°C, and resuspended for several h in TE buffer (10 mM Tris, 1 mM EDTA, pH 8.0). DNA purity and concentration were measured using a Nanodrop 2000 (Thermo Fisher).

### Oxygen sensing system

Oxygen pressure at the bottom of the well, for the experiment in **Figure 1B** was assessed using the PreSens SDR SensorDish system (Applikon Biotechnology) with a 24-well plate.

### Cell culture

#### HEK293FT

The HEK293FT cell line was purchased from Thermo Fisher/Life Technologies. Cells were cultured in DMEM with 4.5 g/L glucose (1 g/L base; 3.5 g/L additional), 3.7 g/L sodium bicarbonate, 10% FBS, 6 mM L-glutamine (2 mM base; 4 mM additional), penicillin (100 U/μL), and streptomycin (100 μg/mL) in a 37°C incubator with 5% CO_2_. Cells were subcultured between a 1:5 to 1:10 ratio every 2–3 d using Trypsin-EDTA. The HEK293FT cell line tested negative for mycoplasma using the MycoAlert Mycoplasma Detection Kit.

#### B16F10

The B16F10 cell line was purchased from ATCC. B16F10 cells were cultured in DMEM with 4.5 g/L glucose (1 g/L, base; 3.5 g/L additional), 3.7 g/L sodium bicarbonate, 10% FBS, 6 mM L-glutamine (2 mM base; 4 mM additional), penicillin (100 U/μL), and streptomycin (100 μg/mL) in a 37°C incubator with 5% CO_2_. Cells were subcultured between a 1:10 to 1:20 ratio every 2– 3 d using Trypsin-EDTA.

### Transfection-based experiments

#### HEK293FT with calcium phosphate

Transient transfection of HEK293FT cells was conducted using the calcium phosphate methodology. Cells were plated at a minimum density of 1.5×10^5^ cells per well in a 24-well plate in 0.5 mL DMEM, supplemented as described above. For surface staining experiments, cells were plated at a minimum density of 3.0×10^5^ cells per well in a 12-well plate in 1 mL DMEM. After at least 6 h, by which time the cells had adhered to the plate, the cells were transfected. For transfection, plasmids were mixed in H_2_O, and 2 M CaCl_2_ was added to a final concentration of 0.3 M CaCl_2_. This mixture was added dropwise to an equal-volume solution of 2× HEPES-buffered saline (280 mM NaCl, 0.05 M HEPES, 1.5 mM Na_2_HPO_4_) and gently pipetted up and down approximately four times. After 2.5–4 min, the solution was mixed vigorously by pipetting approximately ten times. Per well, 100 µL of this mixture was added dropwise to cells plated in 24-well plates or 200 µL was added to cells plated in 12-well plates.. Then, the plates were gently swirled prior to being returned to the incubator. The next morning, the medium was aspirated and replaced with fresh, complete medium. In some assays, fresh medium contained cobalt(ii) chloride (CoCl_2_, to induce the hypoxia response), as described in the pertinent figures. Cells were typically harvested 36–48 h after transfection and at least 24 h after media change. This methodology was used for experiments that were analyzed by flow cytometry, unless the cells were to be cultured in hypoxia, in which case the protocol specified below was used.

#### HEK293FT with lipofectamine

From exponentially growing HEK293FT or HEK293FT-LP cells, 1.0 x 10^5^ cells were plated per well (1.0 mL medium) in 12-well format, and the cells were cultured for 24 h to provide time to attach and spread. When cells reached 50–75% confluence, plasmids were transfected by lipofection using Lipofectamine LTX with PLUS Reagent. Plasmids were mixed with 1.0 μL of PLUS reagent in a 50 μL total volume reaction, with the remainder of the volume being OptiMEM. In a separate tube, 3.8 μL of LTX reagent was mixed with 46.2 μL of OptiMEM. The DNA/PLUS Reagent mix was added to the LTX mix, pipetted up and down four times, and incubated at room temperature for 5 min. 100 μL of this transfection mix was added dropwise to each well of cells, and plates were mixed by gentle swirling. For the microscopy experiment in **Figure 1C**, and **Supplementary Figure 2**, cells were incubated in normoxia overnight and then cultured for 1–2 d in normoxia or hypoxia prior to microscopy. For the oxidation experiment in **Figure 1D** and **Supplementary Figure 3**, a 24-well plate was used rather than a 12-well plate; accordingly, half of the number of cells, volumes of media and reagents, and masses of DNA were used as listed above.

#### B16F10 with lipofectamine

From exponentially growing B16F10 cells, 0.4 x 10^5^ cells were plated per well (24-well plate) with 0.5 mL medium and cultured for 12–24 h to provide time for cells to attach and spread. Up to 400 ng of DNA was diluted to 25 μL total volume reaction with OptiMEM. In a separate tube, 2.5 μL of Lipofectamine LTX reagent was mixed with 22.5 μL of OptiMEM. The DNA mix was added to the LTX mix, pipetted up and down four times, and then incubated at room temperature for 5 min. 50 μL of this transfection mix was added dropwise to each well of cells, and plates were then mixed by gentle swirling and returned to the incubator.

### Microscopy assays

Microscopy was conducted on a Keyence BZ-X800E fluorescence microscope. Cells were maintained at 37°C and supplied with air containing 5% CO_2_, 21% O_2_, and 74% N_2_ using a stage top incubator (TOKAI HIT, INU-KIW-F1). Filter sets were purchased from Chroma in Keyence BZX cubes and are described in **Supplementary Table 3**. Experiments for microscopy were conducted in Phenol Red-free DMEM with 3.7 g/L sodium bicarbonate, 4.5 g/L glucose (1 g/L base; 3.5 g/L additional), 1 mL/L of a pyridoxine-HCl (4 mg/mL) and sodium phosphate (16 mg/mL) solution, Fetal Bovine Serum, L-glutamine, and Penicillin-Streptomycin (as described in **Cell culture**). For transfected samples, cells were transfected with Lipofectamine LTX with PLUS Reagent.

### B16F10-LP development

#### Cas9 and sgRNA expressing vector cloning

pU6-(BbsI)_CBh-Cas9-T2A-BFP-P2A-Ad4E4orf6 (Addgene #64220; referred to as pPD782) and pU6-sgRosa26-1_CBh-Cas9-T2A-BFP-P2A-Ad4E1B (Addgene #64219; referred to as pPD720) (46) were obtained from Addgene. The region encoding the BFP-P2A-Ad4E1B region (flanked by NheI/EcoRI) in pPD720 was replaced by a fragment encoding BFP-P2A-Ad4E4orf6 (flanked by NheI/EcoRI) to generate pPD783 pU6-sgRosa26-1_Cas9-T2A-BFP-P2A-Ad4E4orf6.

#### Landing pad vector cloning

The CAG promoter and homology arms for the Rosa26 locus were obtained from pR26 CAG/GFP Asc (Addgene #74285) (47). The LP for the B16F10 cells was built by assembling the following components into the Destination Vector as per mMoClo standards (34): TU1 contained the Rosa26 left homology arm; TU2 was assembled from the pPart series (CHS4x2 in pInsulator, CAG in pPro, a placeholder 5’UTR in p5’UTR, EYFP-P2A-HygroR in pGene, an inert 3’ UTR in p3’UTR, and rbGlob PA terminator CHS4x2 in pPolyA) and then an attP site was inserted in place of the placeholder 5’UTR; TU3 contained a placeholder homology arm, pLink3. After a Golden Gate assembly, the placeholder homology arm was replaced with a Rosa26 right homology arm. The final vector was named pPD864 and contained the LP flanked by Rosa26 homology arms. The CAG promoter was then replaced with the EF1a promoter with restriction enzyme digest, with the resulting plasmid termed pPD864.

#### Vector integration and cell selection

From exponentially growing B16F10 cells, 4 x 10^5^ cells were plated per well (6-well plate) with 2 mL medium and cultured for 24 h to allow them to attach and spread. Cells were transfected with 160 ng each of pPD720, pPD783, and pPD864, 1520 ng of pPD005, in 100 μL total volume (balance OptiMEM), mixed with 100 μL of OptiMEM solution containing 10 μL of Lipofectamine LTX (total volume of 200 μL transfection mixture per well). Beginning at 3 d after transfection, cells were selected with 1 mg/mL Hygromycin for 15 d. Individual EYFP^+^ cells were sorted into a 96-well plate (see **Flow cytometry-based cell sorting**). Microscopy was used to verify that wells only contained one cell per well (wells with more than 1 cell were discarded from use). Cell lines were expanded for 2–4 weeks in continuous antibiotic selection with Hygromycin. Approximately 60 monoclonal lines were generated.

#### Line validation

Validation was performed by genomic PCR through the Rosa26 right homology arm. Genomic DNA from each line was extracted using the GeneJET Genomic DNA Purification Kit. Lines were also assessed for their ability to maintain EYFP fluorescence for 6 weeks without antibiotic selection and for the performance of an integrated circuit.

### Landing pad integration and cell selection

#### HEK293FT-LP

From exponentially growing HEK293FT-LP cells, 0.5 x 10^5^ cells were plated per well (24-well plate) in 0.5 mL medium and cultured for 24 h to provide time for cells to attach and spread. When cells reached 50–75% confluence, Bxb1 recombinase vector was co-transfected with the integration vector by lipofection using Lipofectamine LTX with PLUS Reagent. 300 ng of Bxb1 expression vector was mixed with 300 ng of integration vector and 0.5 μL of PLUS reagent in a 25 μL total volume reaction, with the remainder of the volume being OptiMEM. In a separate tube, 1.9 μL of LTX reagent was mixed with 23.1 μL of OptiMEM. The DNA/PLUS Reagent mix was added to the LTX mix, pipetted up and down four times, and then incubated at room temperature for 5 min. 50 μL of this transfection mix was added dropwise to each well of cells, which was then mixed by gentle swirling and returned to the incubator. Cells were cultured until the well was confluent (typically 3 d) without any medium changes.

Cells were harvested by trypsinization, transferred to a single well of a 6-well plate in 2 mL of medium, and cultured until they reached 50–70% confluence. The medium was aspirated and replaced with 2 mL of fresh medium containing an appropriate selection antibiotic: 1 μg/mL puromycin and/or 6 μg/mL blasticidin. Selection was performed in puromycin alone for 7 d, with the medium replaced daily with fresh medium containing antibiotics until cell death was no longer evident. Cells were expanded for 7 d without antibiotics then cultured in both puromycin and blasticidin to maintain selective pressure until assay, flow sorting, or freezing. Cells were sorted as described for each experiment and in **Flow cytometry-based cell sorting**.

#### B16F10-LP

From exponentially growing B16F10-LP cells, 0.8 x 10^5^ cells were plated per well (12-well plate) with 1 mL medium and cultured for 24 h to allow cells to attach and spread. Bxb1 recombinase vector was co-transfected with the integration vector by lipofection using Lipofectamine LTX. 200 ng of Bxb1 expression vector was mixed with 200 ng of integration vector in a 50 μL total volume reaction, with the remainder of the volume being OptiMEM. In a separate tube, 5 μL of LTX reagent was mixed with 45 μL of OptiMEM. The DNA mix was added to the LTX mix, pipetted up and down four times, and then incubated at room temperature for 5 min. 100 μL of this transfection mix was added dropwise to each well of cells, which was then mixed by gentle swirling and returned to the incubator. Cells were cultured until the well was confluent (typically 3 d) without any medium changes.

Cells were harvested by trypsinization, transferred to a single well of a 6-well plate in 2 mL of medium with appropriate selection antibiotic, and cultured until reaching 50–70% confluence, with frequent trypsinization to remove dead cells. Antibiotic concentrations were 1 μg/mL puromycin and 15 μg/mL blasticidin. Medium was replaced daily with fresh medium containing antibiotics until cell death was no longer evident and until assay, flow sorting, or freezing. Cells were sorted as described for each experiment and in **Flow cytometry-based cell sorting**.

### Flow cytometry-based cell sorting

Cells were harvested by trypsinization, resuspended at approximately 10^7^ cells per mL in pre-sort medium (DMEM with 10% FBS, 25 mM HEPES, and 100 ug/mL gentamycin), and kept on ice until sorting was performed. Cells were sorted using one of several BD FACS Aria Special Order Research Products with the optical configuration listed in **Supplementary Table 5**. Cells were collected for each line in post-sort medium (DMEM with 20% FBS, 25 mM HEPES, and 100 μg/mL gentamycin). Cells were kept on ice until centrifugation (150 x g for 5 min) and were then resuspended in DMEM supplemented as in **Cell culture**. Cells were plated and expanded until used in experiments. Gentamycin was included in the culture medium for one week after sorting. For monoclonal cell sorting, cells were sorted directly into 96-well plates and maintained in post-sort medium until adherent, at which point the medium was changed as described (*Vector integration and cell selection*). After selection, some lines were frozen in 5% DMSO in completed DMEM (as per **Cell culture**) and stored in liquid nitrogen until later use. Thawed cells were cultured for approximately 2 weeks prior to experimentation at low densities (generally not exceeding 50% confluence).

### Experiments with LP cell lines

Stable cell lines were plated in 0.75 mL of DMEM in triplicate in 24-well format at a density expected to generate 5% confluent wells and placed in a normoxic or hypoxic incubator. For some experiments, lines were plated in medium containing CoCl_2_ at 150 μM (unless otherwise specified) or 1 μg/mL doxycycline. Cells were harvested for flow cytometry analysis at the time point indicated in each figure.

### Flow cytometry assays

#### Cell harvesting

Cells were harvested for flow cytometry using FACS buffer (PBS pH 7.4, 2–5 mM EDTA, 0.1% BSA) or using 0.05% Trypsin-EDTA (with or without Phenol Red) for 5 min followed by quenching with medium (with or without Phenol Red). The resulting cell solution was added to at least 2 volumes of FACS buffer. Cells were centrifuged (150×g for 5 min), supernatant was decanted, fresh FACS buffer was added, and cells were resuspended by briefly vortexing.

#### Data collection and analysis

Flow cytometry was run on a BD LSR Fortessa Special Order Research Product. Lasers and filter sets used for data acquisition are listed in **Supplementary Table 4**. Samples were analyzed using FlowJo v10 software (FlowJo, LLC). Fluorescence data were compensated for spectral bleed-through. The HEK293FT and B16F10 cell populations were identified by SSC-A vs. FSC-A gating, and singlets were identified by FSC-A vs. FSC-H gating. In transfection-based experiments, to distinguish transfected from non-transfected cells, a control sample of cells was generated by transfecting cells with a mass of pcDNA (empty vector) equivalent to the mass of DNA used in other samples in the experiment. For the single-cell subpopulation of this pcDNA-only sample, a gate was made to identify cells that were positive for the constitutive fluorescent protein used as a transfection control in other samples, such that the gate included no more than 1% of the non-fluorescent cells. In landing pad-based experiments, a sample of the corresponding parental, non-landing pad line was used to distinguish cells expressing a given fluorescent protein.

#### Conversion of arbitrary units to standardized fluorescence units

To determine the conversion from the arbitrary unit (au) of fluorescence recorded by the flow cytometer to a standardized fluorescence unit (Molecular Equivalents of a Fluorophore (MEF): Fluorescein (MEFL), PE-Texas Red (MEPTR), or Pacific Blue (MEPB)), Rainbow Calibration Particles or UltraRainbow Calibration Particles were run as part of each flow cytometry experiment. These reagents contain six (RCP) or nine (URCP) subpopulations of beads, each of a specific size and with a known intensity of various fluorophores, that are provided with calibration data for each lot. The total bead population was identified by SSC-A vs. FSC-A gating, and subpopulations were identified through two fluorescent channels (**Supplementary Figure 1**). A calibration curve was generated for the experimentally determined au vs. manufacturer supplied MEF, and a linear regression was performed with the constraint that 0 au equals 0 MEF. The slope from the regression was used as the conversion factor, and error was propagated.

### Reagents

Reagents for Golden Gate reactions are listed in **Supplementary Table 6**. Reagents for cell culture are listed in **Supplementary Table 7**. Reagents for flow cytometer calibration are listed in **Supplementary Table 8**. Kits used in this study are listed in **Supplementary Table 9**. Instrumentation used in this study is listed in **Supplementary Table 10**.

### Biological resources

Plasmid maps for all plasmids generated in this study are supplied as annotated GenBank files in **Supplementary Data 1**. Lists of plasmids obtained from external sources for, plasmids used in this study and previously described by our lab, and plasmids generated in this study are in **Supplementary Table 11**, **Supplementary Table 12**, and **Supplementary Table 13**, respectively. Lists of cell lines obtained for and generated in this study are in **Supplementary Table 14** and **Supplementary Table 15**, respectively. Key plasmids generated in this study will be deposited and made available through AddGene (www.addgene.org).

### Statistical analysis

Unless otherwise stated, three independent biological replicates were evaluated for each condition. The data displayed reflect the mean across these biological replicates of the MFI of generally approximately 3,000 single transfected cells for transfection-based experiments, 3,000-10,000 single, landing pad marker positive cells for experiments with landing pad cells, or for experiments at very low cell densities all available cells. Error bars represent the standard error of the mean (S.E.M.).

ANOVA tests, Tukey’s HSD tests, and t-tests were performed in Python using the statsmodels (48) and Pingouin (49) packages. Tukey’s HSD tests were performed with α = 0.05. Pairwise comparisons were made using a two-tailed Welch’s t-test, which is a version of Student’s t-test in which variance between samples is treated as not necessarily equal. To decrease the false discovery rate, the Benjamini-Hochberg procedure was applied to each set of tests per figure panel, after which the null hypothesis was rejected for P-values < 0.05. Repeated measures ANOVA tests were used for time series data, in which the same samples are used across measurement days.

### Iterative Model Development and Analysis

Model formulation and parameter estimation were implemented using an iterative process based on a previously reported model development workflow (50). First, we defined a subset of the experimental data to use as training data, from which we identified a set of modeling objectives (**Table 1**). These objectives highlight key features of the training data that we aim to describe with the model. Then, we formulated a base-case model using ordinary differential equations (ODEs) to describe the time-dependent evolution of the concentration of each state in the system, with associated assumptions (**Supplementary Table 16**). The base case model is the simplest reasonable description of the data and acted as a starting point for systematically adding complexity when necessary to describe the data. Next, we selected a parameter estimation method (PEM) and ran parameter estimation simulations with synthetic data to ensure that the PEM could recover a parameter set that yields a reasonable fit to the synthetic data, as described in Dray et al (50). We then used the PEM to estimate model parameters based on the training data. The calibrated (i.e., optimal) parameter set was evaluated based on the agreement of the corresponding model simulations with the training data, in terms of the modeling objectives and the quantitative agreement (as measured by the *X*^2^ and R^2^). In the case that the objectives were not satisfied, we proposed biologically motivated mechanistic updates to improve the agreement, which were mathematically formulated and appended to the existing model. This process was iterated until arriving at a final model that satisfied all objectives and yielded adequate quantitative agreement with the training data. More detailed descriptions of this process are provided in **Supplementary Note 1**.

**Table 1.** Modeling objectives and candidate model analysis.

|  | Model A | Model B | Model B2 | Model C | Model D | Model D2 |
| --- | --- | --- | --- | --- | --- | --- |
| Additional mechanism(s) or free parameter →<br>Modeling objective ↓ | Base case | Model A + HAF degradation in short-term hypoxia | Model B + $k_{\text{txn}2}$ (HAF $k_{\text{txn}}$) | Model A + HAF SUMOylation in hypoxia | Model A + HAF degradation and SUMOylation in hypoxia | Model D + $k_{\text{txn}2}$ (HAF $k_{\text{txn}}$) |
| Maximum reporter expression of each feedback HBS > that of simple HBS | Yes | Yes | Yes | Yes | Yes | Yes |
| 24-hour reporter expression of HIF-1α feedback HBS > that of HIF-2α feedback HBS | No | No | No | No | Yes | Yes |
| Reporter expression of HIF-1α feedback HBS reaches max at ~48 hours, then decreases | No | No | No | No | No | Yes |
| Quantitative agreement ($\chi^2$) | 1.10 | 1.11 | 1.11 | 1.11 | 1.13 | 0.50 |
| Quantitative Agreement ($R^2$) | 0.93 | 0.93 | 0.93 | 0.93 | 0.92 | 0.97 |

<sup>1</sup>The additional mechanism(s) or free parameter row indicates the starting model + additional mechanism(s) or free parameter.

### Data Availability

#### Sequence Data Resources

Sequence data for all plasmids is provided in annotated GenBank files (**Supplementary Data 1**).

#### Novel Programs, Software, Algorithms

Code is provided as a public repository on GitHub [https://github.com/leonardlab/HBS_GAMES].

## RESULTS

### Validating methods for studying hypoxia

The study of hypoxia presents several challenges that can introduce artifacts if not well managed (51,52), and therefore we initiated this investigation by developing and validating enabling methodology. One consideration is that when transitioning a cell culture from normoxia to hypoxia, it takes time for oxygen to diffuse out of the medium. It is critical that cells experience the intended oxygen pressure for an accurate interpretation of the hypoxic response in those conditions (51,52). We explored the rate at which oxygen diffuses out of medium using a specialized plate with an electronic oxygen sensor at the bottom of the well as the O_2_ concentration in the incubator was decreased stepwise (**Figure 1B**). We found that in a 24-well plate, culture volumes of 250 µL or less rapidly equilibrated, culture volumes between 500–750 µL exhibited a time lag, and larger volumes required hours to equilibrate. Therefore, in future assays, we used volumes that did not exceed the equivalent depth of 750 µL in a 24-well plate.

Another consideration when studying hypoxia using fluorescent proteins is that fluorescent protein maturation requires an oxidation step for most fluorophores (the biliverdin-based iRFPs are an exception) (53,54). We evaluated the degree to which this maturation requirement impacted several fluorophores in this system (**Figure 1C**, **Supplementary Figure 2**). Cells transfected with plasmids for constitutive fluorescent protein expression were cultured in hypoxic conditions and then serially imaged by microscopy in normoxic conditions and compared to transfected cells that were cultured continuously in normoxic conditions. While miRFP670 and miRFP720 exhibited similar brightness after hypoxic and normoxic culture, other fluorescent proteins required various lengths of time to mature: mTagBFP2 and mNeonGreen matured rapidly within approximately 80 minutes; EYFP, EBFP2, and DsRed-Express2 required ∼2-3 h; and mRuby3 required several hours. These observations provide practical guides for experimental design.

We next compared sample preparation methods to evaluate and minimize potential bias due to differential effects upon fluorophore maturation. For the fastest maturing fluorophore of each color, we evaluated how several common methods of sample preparation could affect oxidation and thus maturation. Cells co-transfected with plasmids for constitutive fluorescent protein expression were cultured in normoxia or hypoxia and then subjected to various treatments prior to analysis (**Figure 1D**, **Supplementary Figure 3**). Compared to cells cultured under normoxia, cells cultured under hypoxia and then immediately taken for flow cytometric analysis after harvest (with intermediate storage either on ice or at room temperature) exhibited lower fluorescence, and this effect was most pronounced for DsRed-Express2. Cells incubated for 2 h in FACS buffer after harvest also displayed lower fluorescence compared to normoxic culture controls, and maturation was more complete for samples incubated at higher temperatures after harvest. In contrast, cells cultured in hypoxia and then cultured in a normoxic incubator for 2 h prior to harvest displayed complete oxidation of mTagBFP2, nearly complete oxidation of mNeonGreen, and 80% oxidation of DsRed-Express2. Cells that were fixed with paraformaldehyde after harvest and then oxidized at various temperatures displayed complete oxidation of mTagBPF2, near complete oxidation of mNeonGreen, and only minimal oxidation of DsRed-Express2. Given the timescales over which the oxidation of these fluorescent proteins occurs (**Figure 1C**, **Supplementary Figure 2**) and the minimal effect of temperature on these samples compared to that of unfixed samples, it is possible that some oxidation occurred during sample processing prior to complete fixation, and that fixation then inhibited the oxidation from progressing. Considering all of these factors, in subsequent experiments we used a protocol including a 2 h period of normoxic culture after the hypoxic culture was completed to allow for maturation of the fluorescent proteins.

To validate our protocols for studying induced gene expression under hypoxia, we applied these methods to a simple test case in which control of gene expression is well understood. We used a simple gene circuit in which doxycycline-inducible expression of DsRed-Express2 is driven from a single-copy, genomically integrated locus of the HEK293FT cell line. To do so, we employed a genomic landing pad (LP) system that was developed for rapidly prototyping genetic constructs in a genomic context (34,55,56). An LP is a recombinase target site genetically inserted into a genomic safe harbor locus, enabling insertion of large DNA constructs in a site-specific fashion using a transposase (**Figure 1E**). LPs have advantages over other methodologies, such as lentiviral transduction, including a higher limit on cargo size. Additionally, as the cells with cargo integrated into the landing pad locus are genetically identical, LP methodology makes the resulting population more homogenous. In this evaluation, cells with an integrated circuit that expresses DsRed-Express2 only in the presence of doxycycline (dox) (**Figure 1E**) were cultured in hypoxia or normoxia for four days with or without doxycycline, and a sample was harvested each day for analysis (**Figure 1F**). DsRed-Express2 levels from the hypoxic and normoxic conditions were largely similar over the course of 4 days, readily distinguishing dox-induced from uninduced conditions, which is the expected outcome and supports our protocol. We speculate that the slight differences observed between induced cases are attributable to minimally incomplete oxidation and cellular stress from hypoxia. Altogether, this experiment validates our oxidation and harvest protocols for studying regulation of gene expression in hypoxia.

### Characterization of a base case hypoxia biosensor

Toward building HBS variants exhibiting various performance characteristics, we first evaluated a base case biosensor. To facilitate characterization and eventual comparison across designs, we constructed a simple HBS construct and inserted it into the genome of the HEK293FT-LP cell line (**Figure 2A**). Our HBS employed the YB_TATA minimal promoter, which has been shown to confer low background expression (31), downstream of a set of HREs in an arrangement that has been shown to yield high hypoxia-induced gene expression and fold induction (20). In this construct, EBFP2 and blasticidin resistance genes were constitutively expressed, a puromycin resistance gene was only expressed if the construct was integrated downstream of a genomic promoter (as an indicator of successful genomic integration), and expression of DsRed-Express2 was controlled by the HBS. After integration, cells were selected with antibiotics for two weeks and then expanded. As a first test of activity of the HBS, we employed cobalt(II) chloride (CoCl_2_) treatment, which induces the hypoxia response system (**Figure 2B,C**) (57). CoCl_2_ induced a 19-fold increase in reporter expression, validating HBS function. However, this response was heterogenous, and many cells did not express the reporter. Analytically dividing the cells into ten equal-proportion decile bins (post hoc) for EBFP2 expression revealed that cells with higher EBFP2 exhibited higher reporter expression— both with and without CoCl_2_—such that fold induction was similar across these bins (**Figure 2C**). We hypothesized that since these cells were putatively genetically identical, the differences in reporter expression could be explained by either non-heritable differences (such as metabolic activity, cell cycle state, etc.) or epigenetic, and potentially heritable, differences. Since distinguishing these contributions could be important for understanding opportunities for improving biosensor performance, we decided to investigate further.

**Figure 2.**
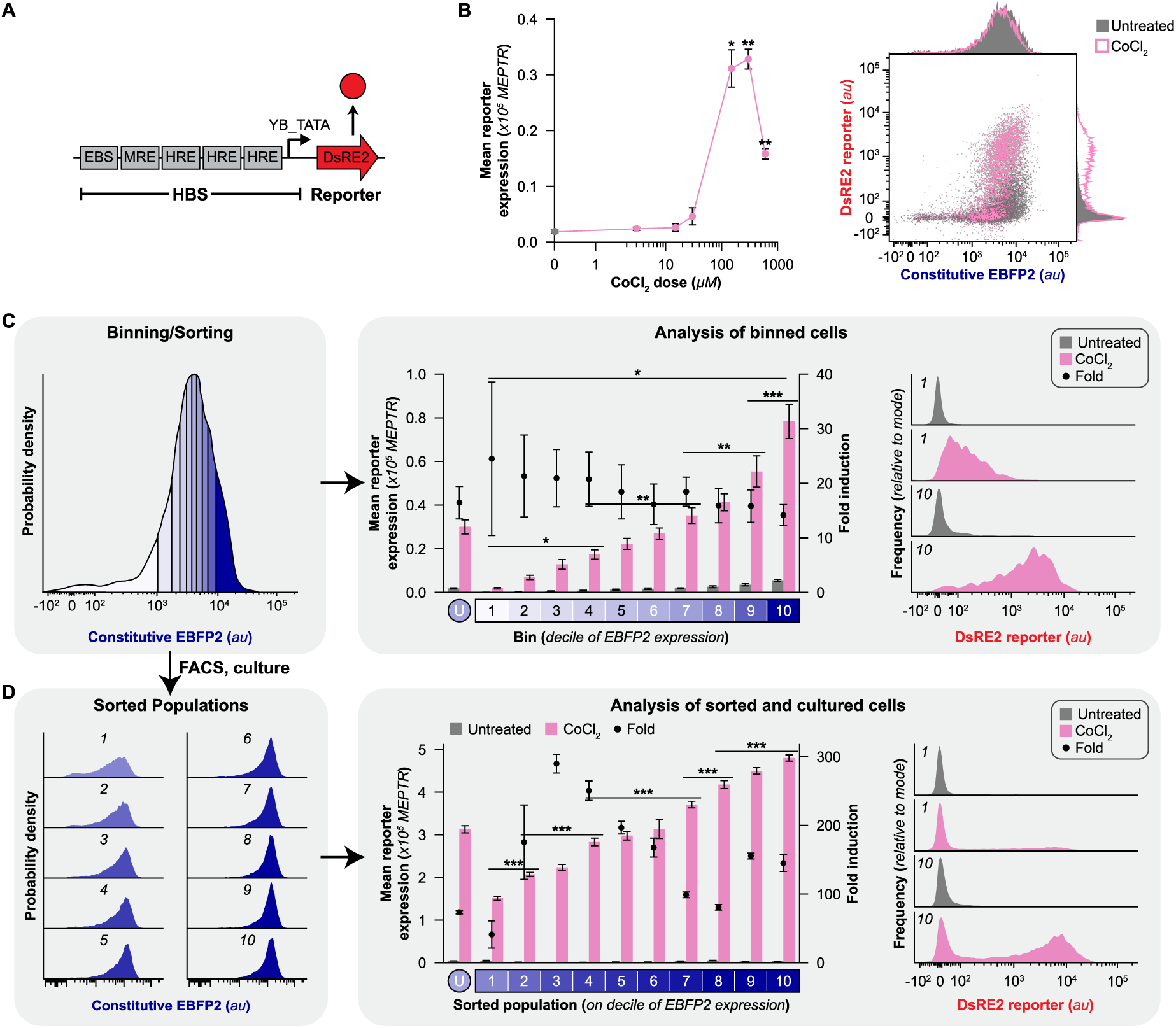
Heritable performance differences in a genomically integrated HBS. (**A**) Schematic of a HBS that produces DsRE2. (**B**) Cobalt-induced validation of novel HBS activation. HEK293FT-LP cells with the HBS shown in **A** integrated into the LP were cultured with CoCl_2_, a hypoxia mimetic, and analyzed by flow cytometry. Significant differences are indicated relative to the untreated condition for each CoCl_2_ dose (two-tailed Welch’s *t*-test, * P < 0.05, ** P < 0.01). Right: a representative flow cytometry plot of the 150 µM CoCl_2_ condition. (**C-D**) Investigating the heritability of variable HBS performance. (**C**) Post-hoc subpopulation analysis of flow cytometry data from cells cultured with 150 µM CoCl_2_ in **B**. For each of 3 samples, cells were subdivided by level of constitutive EBFP2 expression into 10 bins with equal numbers of cells and metrics calculated for each subpopulation (U indicates the unsorted population). Horizontal lines are placed to indicate the first bin (moving right) that differs significantly from the reference bin (at left of the bar) (2-way ANOVA for mean reporter expression and two-tailed Welch’s *t*-test for fold induction, * P < 0.05, ** P < 0.01, *** P < 0.001). (**D**) Cells were then sorted by FACS into 10 populations based on the EBFP2 level, expanded for approximately 2-3 weeks and then cultured with or without CoCl_2_ prior to analytic flow cytometry (U indicates the unsorted population). Horizontal lines are placed to indicate the first subpopulation (moving right) that differs significantly from the reference subpopulation (at left of the bar) (2-way ANOVA, *** P < 0.001). Throughout figure, n=3, error bars represent SEM; histograms in **C**, **D** are representative samples. Outcomes from t-tests for **B** are in **Supplementary Note 8** and outcomes from ANOVAs and Tukey’s HSD tests for **C-D** are in **Supplementary Note 6**.

We evaluated whether selecting cells for high constitutive proxy marker (EBFP2) levels could yield a stably homogenous, high-magnitude HBS response. The parental cell line was sorted into ten (polyclonal) populations based on decile of EBFP2 expression, cells were cultured at a low density for two weeks, and HBS activation was again evaluated using CoCl_2_ treatment. In support of a heritable model of HBS performance differences, the magnitude of CoCl_2_-induced reporter expression increased with the expression level of the EBFP2 measured at the time of sorting (**Figure 2D**). These differences were not driven by an increase in the level (mode) of reporter expression in the reporter-expressing cells within any one population, but rather by an increase in the fraction of cells that exhibited reporter activation within each population (**Supplementary Figure 4**). A similar trend was observed when quantifying the constitutive EBFP2 proxy within each sorted subpopulation: cells with higher EBFP2 expression at the time of sorting were more likely to continue to express EBFP2. Therefore, enriching for cells that highly express the cargo (e.g., proxy marker) can improve HBS performance and homogeneity, and since this phenotype was at least somewhat stable, we hypothesize that this improvement was driven by selecting for cells that evade silencing during the initial period after integration when the risk of silencing could be most acute (58).

### Evaluating the role of minimal promoters

As the choice of minimal promoter is a determinant of HBS performance by influencing both background and induced gene expression (31), we evaluated several minimal promoters (**Figure 3A**). Cells were transfected with plasmids containing HBS variants with various minimal promoters and treated with CoCl_2_. In HEK293FT cells: SV40_min led to high background, which we interpret is due to the presence of the SV40 large T antigen in this line (SV40_min and the SV40 origin of replication have sequence homology); YB_TATA conferred very low background expression leading to a fold induction of 108; and CMV_min increased both the induced and background output (compared to YB_TATA), yielding a fold induction of 54 (**Figure 3B**). To investigate how these performance characteristics depend on choice of cell type, we used the B16F10 murine melanoma cell line, which, unlike the HEK293FT line, is derived from a tumor and might have adaptations to survive in hypoxia. In the B16F10 line, YB_TATA yielded much lower background expression than did SV40_min or CMV_min, leading to the largest fold induction of the set albeit with lower induced expression as well. These trends agree with those previously described for these minimal promoters (31), and in both cases, YB_TATA-based HBS designs conferred larger fold inductions than did SV40_min-based HBS designs.

**Figure 3.**
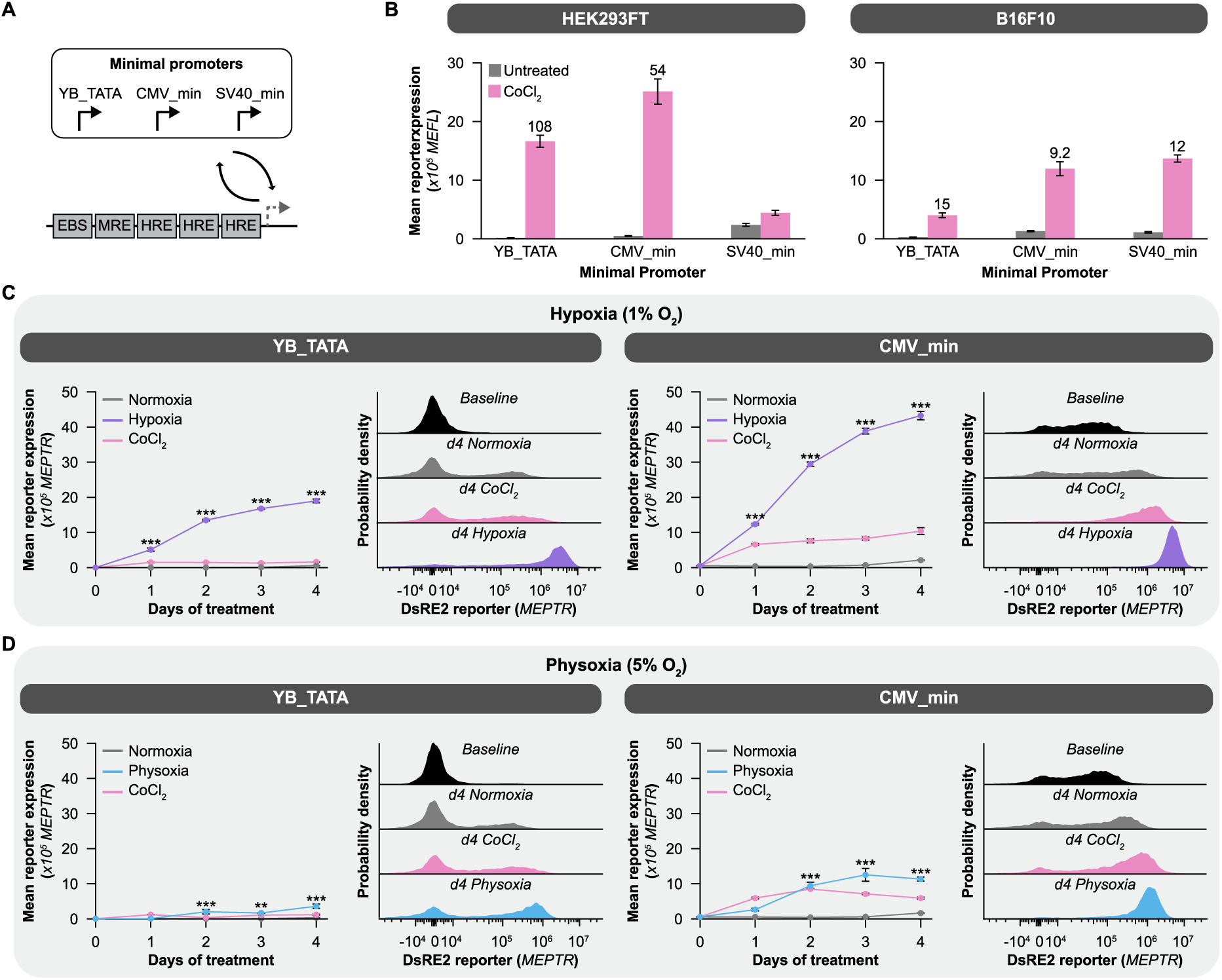
Optimizing HBS performance by minimal promoter choice. (**A**) Schematic of HBSs with several minimal promoters. (**B**) Comparison of HBS variants under cobalt-induction. HBS constructs in **A** were delivered by transient transfection, and CoCl_2_-induced activation was analyzed by flow cytometry. Fold induction is indicated above bars where differences relative to the untreated condition are statistically significant (2-way ANOVA, both P < 0.001). (**C-D**) Evaluation of HBS variant activation in hypoxia (**C**) or physoxia (**D**). HEK293FT-LP with HBSs were cultured as indicated, with samples collected each day for analysis by flow cytometry. Significant HBS activation under each oxygen level is indicated relative the normoxic control on each given day (2-way repeated measures ANOVA for each promoter, ** P < 0.01, *** P < 0.001). Throughout figure, n=3, error bars represent SEM, and histograms are representative samples. Outcomes from ANOVAs and Tukey’s HSD tests for **B** are in **Supplementary Note 6** and outcomes from ANOVAs and Tukey’s HSD tests for **C and D** are in **Supplementary Note 7**.

We next investigated performance of these HBS variants when genomically integrated into HEK293FT-LP cells. In these experiments, the SV40_min construct could not be evaluated, as cells did not survive selection, potentially due to the presence of the SV40 large T antigen. After antibiotic selection and FACS, cells were cultured for 4 d in hypoxia, in normoxia, or in normoxia with CoCl_2_, and samples were harvested each day for analysis by flow cytometry. For both YB_TATA and CMV_min HBS designs, exposure to hypoxia induced higher output that did treatment with CoCl_2_, driven by increases in both the mode of reporter expression and the fraction of cells expressing the reporter (**Figure 3C**, **Supplementary Figure 5**). HBS output increased substantially over the first 3 d of hypoxia treatment, with less marginal increase after that point. Although the CMV_min HBS conferred a more homogenous response to hypoxia than did the YB_TATA HBS, CMV_min was also leakier; in the normoxic cultures, cells began expressing small amounts of reporter by day 4 (**Figure 3C**, inset histograms). We hypothesize that this increase at later timepoints might be due to cell overgrowth and resultant local hypoxia, as previously the cells had been cultured for several weeks at low density without spurious HBS activation.

Finally, we investigated how HBS designs perform under physoxia—5% O_2_, a concentration that approximates the partial pressure of O_2_ in many human tissues. For both CMV_min and YB_TATA HBS constructs, some reporter induction was observed after a few days of culture (**Figure 3D, Supplementary Figures 6, 7**), although much less than was observed after culture at 1% O_2_ (**Figure 3C**). Microscopy analysis of these cultures indicated that reporter output was detectable primarily at the center of clumps of cells (differing from the spatially widespread reporter activation after CoCl_2_ treatment) (**Supplementary Figure 7**), supporting the interpretation that local hypoxia due to overgrowth drives HBS activation, and this may occur at a lower cell density when cells are cultured at 5% O_2_ than was required to trigger this phenomenon at 21% O_2_. Overall, these analyses support the interpretation that our HBS constructs respond to hypoxia.

### Exploring strategies for increasing hypoxia-induced HBS output

To identify opportunities to improve HBS performance, we performed an experimental sensitivity analysis. Given that the YB_TATA HBS exhibited the desirable property of very low background expression in normoxia (compared to alternative designs), we opted to focus on exploring whether the magnitude of hypoxia-induced output could be increased. We first employed a previously validated human HIF1α mutant (HIF1α*) that is stable and induces gene expression even in the presence of oxygen (35). In HIF1α*, the three proline residues that are the targets of oxidation are mutated to alanine residues, and therefore HIF1α* does not degrade in the presence of oxygen (**Figure 1A**). We placed this stable HIF1α* gene under the control of a doxycycline-responsive promoter and integrated a construct including this transcription unit, the YB-TATA-based HBS, a constitutive EBFP2 with antibiotic selection marker, and the tetON system into HEK293FT-LP cells (**Figure 4A**). Following selection, flow sorting, and recovery, we evaluated performance of these new HBS lines (**Figure 4B, C, Supplementary Figure 8**). Doxycycline-induced expression of the stable HIF1α* led to higher levels of induced reporter gene expression than did culture with CoCl_2_ or in hypoxia, and the cell population was more homogenous. Therefore, in HEK293FT cells, since supplementation with exogenous HIF1α* increases HBS-induced gene expression, we conclude that the levels of endogenous HIF1α limit HBS output. This insight identified opportunities for constructing novel HBS architectures.

**Figure 4.**
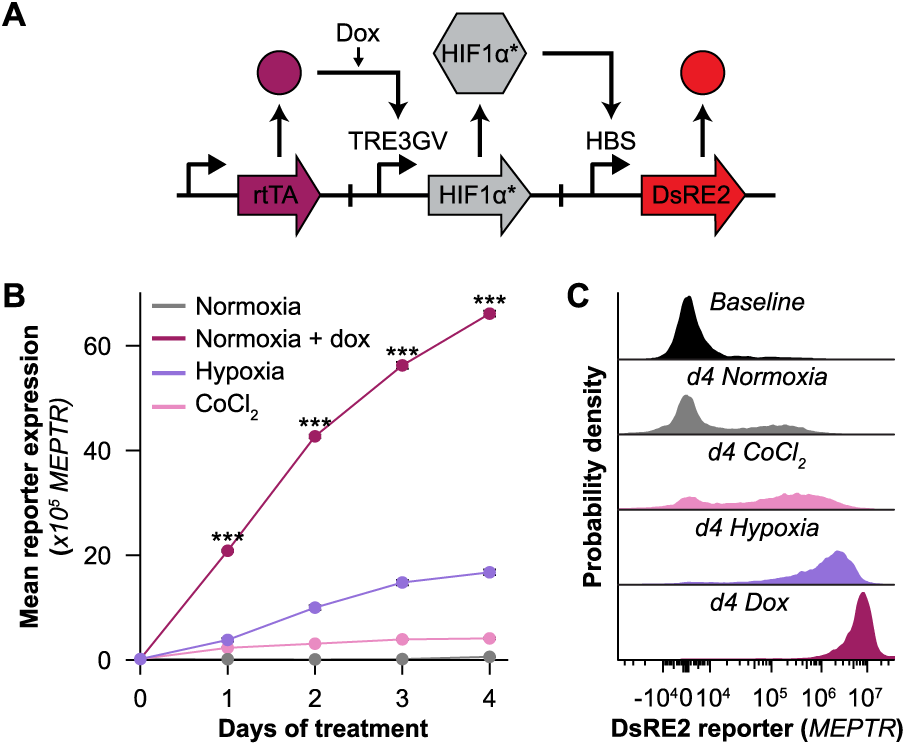
Ectopic overexpression of HIF1α can increase HBS output. **(A)** Schematic depicting a circuit that, upon activation by doxycycline, produces HIF1α*, a HIF1α mutant which is resistant to oxidation and is therefore constitutively stable and active. **(B)** Evaluating the potential to increase HBS output by complementing endogenous HIF1α pools. HEK293FT-LP with the circuit shown in **A** were cultured as indicated, with representative histogram insets depicting day 4 HBS output. Significant differences relative to the normoxia condition on a given day are indicated (2-way repeated measures ANOVA, *** P < 0.001). N=3, error bars represent SEM. Outcomes from ANOVAs and Tukey’s HSD tests for **B** are in **Supplementary Note 7**.

Building upon the aforementioned mechanistic insights into drivers of HBS performance, we explored HBS designs comprising genetic circuits based upon feedback. As HIF1α regulates hypoxia-responsive genes in conjunction with HIF2α and HIF1β, we designed genetic circuits with the goal of increasing the supply of the wild type versions of these proteins and HIF1α. Since overexpressing these factors constitutively could be problematic due to activation of endogenous targets of the hypoxia response system (i.e., if over-expression exceeds the capacity of natural degradation mechanisms), we instead placed the expression of each regulator under the control of a HBS promoter, thereby implementing positive feedback (**Figure 5A**). As a comparator, we built several circuits to investigate whether simply increasing the number of copies of the HBS at each locus could increase hypoxia-induced reporter expression.

**Figure 5.**
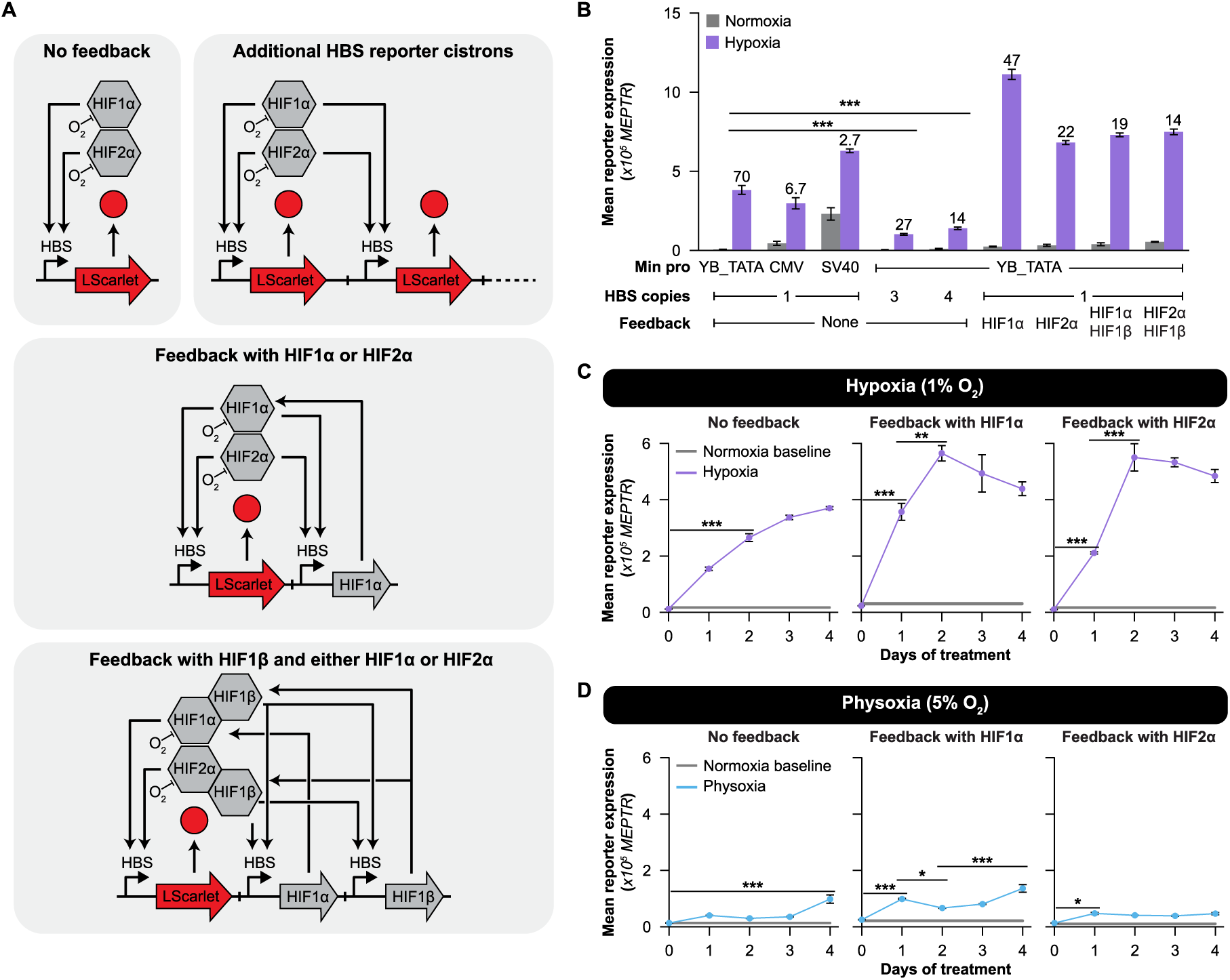
Genetic circuits can enhance HBS performance. (**A**) Schematic summarizing the set of HBS design variants and circuits explored, grouped by strategy and topology (LScarlet indicates LumiScarlet). (**B**) Functional evaluation of HBS designs from **A** implemented in B16F10-LP cells. Lines were cultured for 3 days as indicated and then analyzed by flow cytometry (n=3). Fold induction is indicated where differences relative to the normoxia condition are significant (2-way ANOVA, all P < 0.001). (**C-D**) Evaluation of HBS circuit performance dynamics in hypoxia (**C**) and physoxia (**D**). HBS circuits implemented in B16F10-LP cells were cultured as indicated and analyzed by flow cytometry daily. Baseline was calculated by averaging samples from this line cultured in normoxia at a low density on 3 different days (n=3 samples, n=3 days, width represents SEM). Horizontal lines are placed to indicate the first subpopulation (moving right) that differs significantly from the reference subpopulation (at left of the bar) (2-way repeated measures ANOVA for all 3 topologies simultaneously, * P < 0.05, ** P < 0.01, *** P < 0.001). Throughout the figure, error bars represent SEM. Outcomes from ANOVAs and Tukey’s HSD tests for **B** are in **Supplementary Note 6**, outcomes from ANOVAs and Tukey’s HSD tests for **C-D** are in **Supplementary Note 7**, and outcomes from t-tests for **C-D** are in **Supplementary Note 8**.

Given its relevance to cancer biology research, we opted to explore these designs in the B16F10 murine melanoma system, thus using circuits employing murine hypoxia regulatory factors. We generated a B16F10-LP cell line by integrating a landing pad into the Rosa26 safe harbor locus using Cas9-mediated integration, with inhibition of the NHEJ pathway to increase efficiency (**B16F10-LP development**, **Supplementary Figure 9**). Fluorescent cells (those with an integrated landing pad) were flow-sorted and expanded as monoclonal populations. These lines were validated by flow cytometry, genomic PCR, and integration of a test circuit (**Supplementary Figure 10**).

We next investigated the performance of our novel HBS circuits integrated in the B16F10-LP line. Generally, the choice of minimal promoter displayed the same overall trends in genomically integrated constructs as in circuits that were transiently transfected (**Figure 3B**)— YB_TATA-based HBS designs exhibited the largest fold induction, driven by low background expression (**Figure 5B**). In general, increasing the copy number of base case (open loop) HBS designs did not increase reporter output; this change instead decreased reporter output. A plausible explanation is that increasing HBS copy number dilutes the limited supply of HIF1α across multiple promoters, potentially leading to a loss of cooperative transcriptional activation, as observed in other systems with multiple TF binding sites in a single promoter (33). Notably, gene circuits including positive feedback motifs employing HIF1α or HIF2α exhibited increased hypoxia-induced output (**Figure 5B, Supplementary Figure 11**). The magnitude of HBS output was most increased for HIF1α feedback. The addition of HIF1β positive feedback to either of these strategies conferred no further increases in hypoxia-induced signaling, suggesting that HBS signaling is not limited by HIF1β in these cells. These observations support the use of feedback-based gene circuits to enhance HBS performance.

We next investigated how gene circuits conferring positive feedback influence the dynamics of the HBS response. Given that in prior experiments, overcrowding of the cells in normoxia led to artifactual HBS induction, in this experiment, we quantified background (i.e., normoxic) signaling as a 3-day average baseline. Notably, while the base case (no feedback) HBS required 4 days in hypoxia to reach maximal output, HBS designs with positive feedback exhibited a rapid increase to maximum output by day 2 (**Figure 5C, Supplementary Figure 12-13**). Interestingly, signal then proceeded to decrease for both feedback circuits after day 2, with a subset of cells turning off. Importantly, the addition of positive feedback only mildly increased HBS activation under normoxia and physoxia compared to the base case HBS topology (**Figure 5D, Supplementary Figure 12**). Altogether, employing positive feedback mediated by HIF1α or HIF2α can enhance the performance of HBS constructs while avoiding spurious activation under conditions outside true hypoxia.

### A mechanistic mathematical model provides novel insights into the hypoxia response

To gain more insight into why our biosensor designs perform differently and how they might interact with the native hypoxia response, we developed an explanatory mathematical model. Explanatory models are useful tools for improving understanding beyond what is accessible to intuition alone, as these models comprise a way to quantitatively test whether a set of proposed mechanisms can explain a set of experimental observations. Ordinary differential equations (ODEs) are often well-suited for describing systems and observations such as those of interest here, as ODEs describe the time-dependent evolution of component abundances as governed by hypothesized mechanisms (reactions). Model development is described in detail in **Supplementary Note 1**, and here we focus on key conclusions.

Toward the goal of building the understanding required to relate biosensor design to performance, we formulated a model to describe both native hypoxia regulation and our hypoxia biosensors, integrating and building upon existing biological understanding and prior modeling investigations (**Figure 6**). The oxygen-dependent degradation of the HIFs occurs via a von Hippel–Lindau protein (pVHL) dependent pathway and previously has been modeled for HIF1α (13,15,17,59,60). However, while several other potential modes of HIF regulation have been suggested based on experimental observations, there has been no validation of these hypotheses or mechanistic models of these types of regulation (13–15,18,19,61). Our modeling effort thus utilized previously reported modes of HIF regulation and explored multiple potential modifications of this base model as needed to explain our observations. To guide model development, we first set goals for the model, including recapitulation of the unexpected decrease in reporter expression observed after 2 d for HBS designs employing feedback. These goals were defined as three qualitative objectives based on experimental observations (**Table 1**), and we iteratively formulated and tested candidate models to evaluate the extent to which each one was consistent with experimental observations. Since each model is a mathematical implementation of assumptions on underlying mechanisms, comparing model structures with experimental data enables evaluation of which explanation is most consistent with the observations.

**Figure 6.**
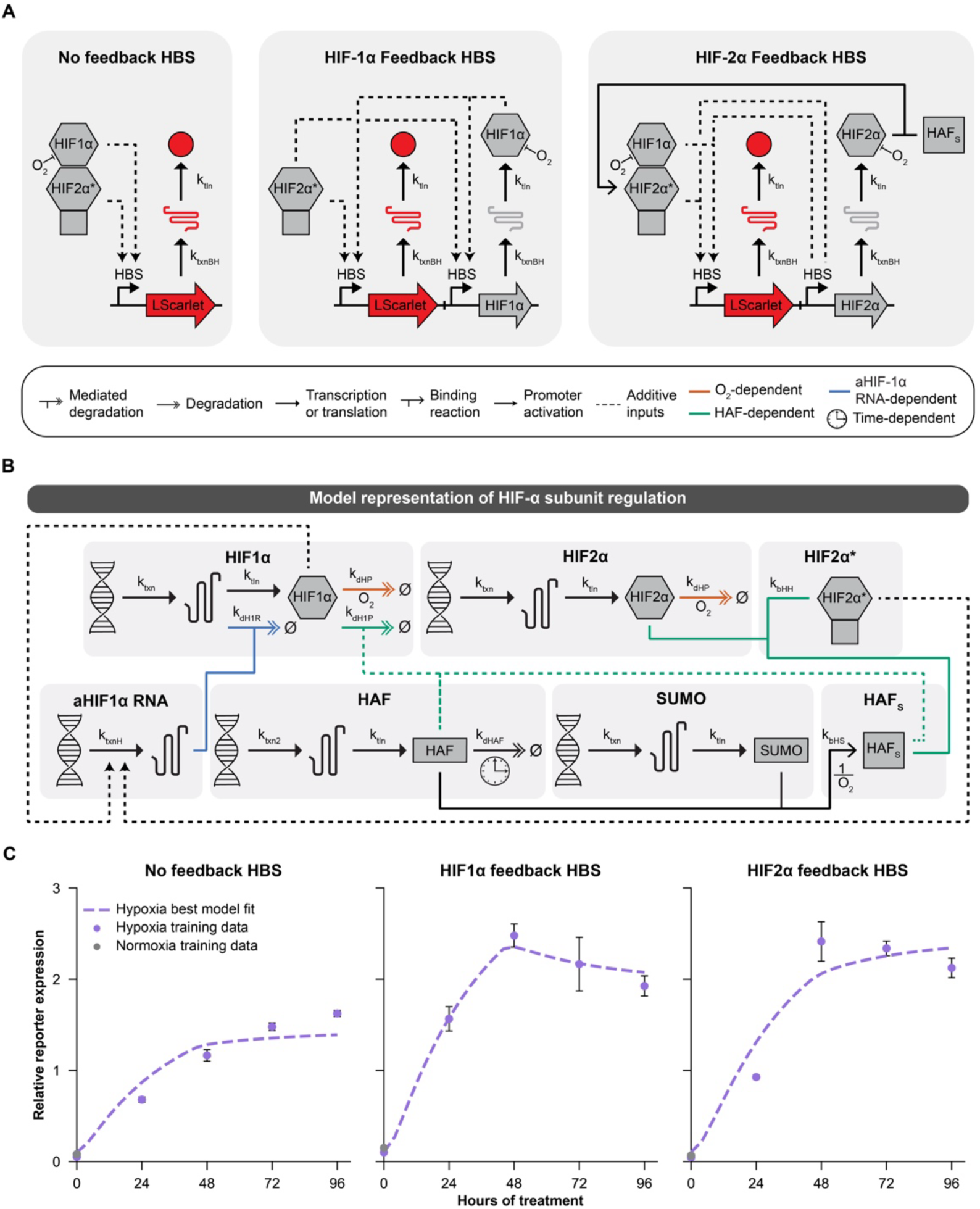
A mechanistic model explains differences in HBS performance across designs. (**A**) HBS topologies used in the model. In the no feedback HBS, HIF1α and active HIF2α-HAF complex (HIF2α*) drive expression of LumiScarlet. In the feedback topologies, the HIFs also drive expression of ectopic HIF1α or HIF2α (LScarlet indicates LumiScarlet). (**B**) Regulation mechanisms of HIF-α subunits incorporated into the model. Both HIFs are degraded at the protein level in an O_2_-dependent manner. HIF1α protein is also degraded by HAF, regardless of SUMOylation. HIF1α mRNA is degraded by antisense HIF1α RNA (aHIF1α RNA). HIF2α protein binds SUMOylated HAF (HAF_S_), forming active HIF2α-HAF complex (HIF2α*), which transactivates target genes. (**C**) Collective model fit to training data describing all three HBSs. Relative reporter expression indicates the experimental or simulated reporter expression normalized to the average reporter expression in the no feedback HBS time series. Model parameters with calibrated parameter values are in **Supplementary Table 18**, and ODEs for the simple HBS are in **Supplementary Table 17**. ODEs for Models A, B, and C (see **Table 1**) for the simple HBS are in **Supplementary Table 19, Supplementary Table 20,** and **Supplementary Table 21** respectively. **Supplementary Note 3** details the required updates to the ODEs comprising the simple HBS models to arrive at the set of equations describing each of the feedback-containing circuits.

Our strategy started with a base case model that is well-supported by prior work, and we added new elements only when required to explain our (new) experimental observations. The base case HBS model incorporates mechanisms for HIF TF-dependent reporter output (LumiScarlet) gene expression via transcriptional regulation, O_2_-dependent HIFα subunit protein degradation, and O_2_-independent HIFα subunit mRNA and protein regulation based on previously reported experimental observations and known mechanisms that have been previously modeled (**Figure 6A, B**, **Supplementary Table 19**). Our description of the oxygen-dependent degradation of the HIFs was premised on reported models describing regulation of HIF1α (13,15,17,59,60). To fill in the remaining parts of this model, in the absence of known or previously modeled mechanisms, we proposed mechanisms based on experimental observations and previously reported (or hypothesized) modes of regulation, and we briefly introduce each proposal here. At the mRNA level, antisense HIF1α RNA (aHIF1α RNA) accumulates in hypoxia, while HIF1α mRNA half-life decreases in prolonged hypoxia (13–15). It has been suggested that aHIF1α RNA could bind to the HIF1α mRNA 3’ untranslated region and expose AU-rich elements, thereby increasing the degradation rate of HIF1α mRNA (14). Additionally, the aHIF-1α promoter contains HREs, so transcription driven by this promoter is plausibly upregulated by the HIFs (14). At the protein level, hypoxia-associated factor (HAF), an E3-ubiquitin ligase, binds both HIFα subunits. HAF binds and ubiquitinates HIF1α through a described mechanism. However, HAF binds HIF2α at a different site, and in its absence a decrease in HIF2α was observed, suggesting that HAF–HIF2α association promotes HIF2α-mediated transactivation of target genes (18,61). After representing each of these mechanisms as equations, we found that this base case model formulation could not satisfy the second and third modeling objectives (**Supplementary Figure 15**, **Table 1**) for any choice of parameters. This limitation motivated the need to include additional mechanisms to explain, in particular, the differences in longer-time behaviors of HBS designs including HIF1α vs. HIF2α feedback.

We hypothesized that additional modes of HAF regulation not included in the base case model could be necessary to describe the differential behavior in the HIF1α vs. HIF2α feedback HBSs. An initial decrease in HAF protein levels in 1% O_2_ has been experimentally observed in human glioblastoma (U87) and pancreatic cancer (PANC-1) cells, with recovery of HAF protein at around 48 h in both cell types (18), supporting the incorporation of a HAF degradation mechanism. However, HAF mRNA levels were not affected by prolonged exposure to 1% O_2_ (18), so the exact mechanism of HAF degradation and recovery is not known. In the absence of a plausible HAF regulation mechanism, we used a heuristic piecewise function to describe the time dependence of the degradation parameter for HAF, k_dHAF_, in 1% O_2_:

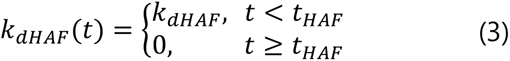

Here, t_HAF_ (a free parameter) is the time at which HAF degradation stops. Additionally, post-translational modification of HAF by a small ubiquitin-related modifier (SUMO) protein (SUMOylation) has been experimentally observed in 786-O clear-cell renal cell carcinoma cells (19). Levels of SUMOylated HAF increased in 1% O_2_ relative to 3% or 20% O_2_ (19), suggesting that HAF SUMOylation could be induced by hypoxia. In the absence of SUMO, a decrease in HIF2α was observed, and it has been posited that HAF SUMOylation is required prior to its binding to HIF2α to form a transcriptional complex that can drive maximal induction of HIF2α target genes (19). We represented HAF SUMOylation in the model as an additional binding reaction between HAF and SUMO to produce SUMOylated HAF (HAF_S_). Both HAF and HAF_S_ can bind HIF1α in the model, but only HAF_S_ can bind HIF2α. After calibration (parameter estimation), we found that a revised model including both mechanisms (model D2) met all 3 objectives (**Table 1** and **Figure 6C**). In contrast, modifying the base case model to include either one of these mechanisms alone (models B, B2, C), did not satisfy all 3 objectives. Altogether, these observations suggest that both HAF regulation mechanisms are required to explain the differential regulation of the HIFs in hypoxia, which is a conclusion that has not previously been reported. Thus, our modeling effort provides new insights into engineered HBS function as well as native hypoxia regulation, making use of behaviors that might be uniquely observable using genetic circuits to perturb the native hypoxia response.

## DISCUSSION

We evaluated several methodologies for studying hypoxia and characterized factors that can potentially influence these investigations (**Figure 1**). Increasing the volume of medium in a well increases the time required for the medium to acclimate to hypoxic conditions, and managing this requirement must be balanced with having sufficient medium in the well for cells to survive, or methods must be employed to allow for medium replacement with fresh de-oxygenated medium for longer-term culture under hypoxic conditions. If one is using fluorescent proteins as reporters in hypoxia, the samples must be treated so that the fluorescent proteins have sufficient time to mature, or fluorophores that do not require oxidation must be used. Cells must be plated at a sufficiently low density and evenly distributed over the plate to minimize formation of clumps of cells, which become hypoxic at their center, causing reporter protein expression in response to true local hypoxia despite normoxic culture conditions. This experiment also highlights the unique insights accessible through single-cell measurements such as by flow cytometry or microscopy, which might be obscured if biosensor outputs were measured in bulk by luciferase, SEAP, or other plate reader-based assays.

Our investigation into the core HBS yielded knowledge that informs the study of hypoxia and design of synthetic biosensors. Minimal promoter choice substantially influences biosensor performance and across multiple studies the YB_TATA confers both low background expression and reasonable, inducible gene expression that can be increased through iterative design and yield high fold inductions (31,33,62). We identified that the magnitude of induced gene expression driven from the HBS was limited by the endogenous pool of HIF1α, at least in the cell lines tested (**Figure 4**). This finding motivated us to build genetic circuits that implemented positive feedback using HIF1α and HIF2α to effectively amplify biosensor output. Excitingly, these feedback circuits amplified hypoxia-induced biosensor output with minimal increases in background in normoxia and physoxia (**Figure 5**). This property was plausibly conferred by two related principles: (i) the limiting HIF proteins are expressed only after the circuit is activated, avoiding constitutive overexpression that could overwhelm cellular degradation and lead to leaky biosensor output in normoxia, and (ii) induction of feedback was possible only in hypoxia (i.e., the magnitude of the feedback gain is greatly increased in hypoxia), as any leaky ectopic HIF is degraded in normoxia or physoxia, providing a fail-safe. These principles may be useful in the design of other synthetic biosensor circuits, conferring similar benefits by combining positive feedback with fail-safes.

This study produced tools that can be used to study hypoxia. The HBS designs characterized here exhibit improved performance compared to prior ones, with high induction of output gene expression at early time points after exposure to hypoxia. These designs could be used in the study of many of the physiologic and pathologic processes involving hypoxia, such as development, cancer, and ischemic heart disease (1–6). The biosensors developed here can be stably deployed in cell lines, which indicates potential for implementation in other cell types, monitoring over long time scales, and potentially extension to transgenic animal models. Additionally, as a melanoma cell line in which various genetic cargo can be stably integrated into the genome, the B16-F10 LP line represents a resource that could be used to study melanoma biology beyond hypoxia. We anticipate that this toolkit could facilitate cross-comparison across model systems and applications in which hypoxia is an important contributor.

An innovation in this study is the use of genetic circuits to perturb the natural hypoxia system and elicit new behaviors that improve our understanding. Towards this goal, we developed a mathematical model that explains the distinct behaviors of three related HIF-regulated HBS designs (**Figure 6**). Combined with empirical observations, this analysis generated mechanistic understanding of both HBS performance and HIF regulation in hypoxia. Our findings support HAF’s reported role as a “HIF switch”, shifting primary hypoxic regulation from HIF1α-driven gene expression at short time scales to HIF2α-driven gene expression at longer time scales (13,15,18). Our findings also motivate the novel hypothesis that differential HIF regulation in hypoxia seems to require both HAF degradation in short-term hypoxia and HAF SUMOylation prior to HIF2α binding. These testable postulates represent an avenue of potential future investigation.

Elucidating the mechanisms driving HBS regulation also conferred insights into potential biosensor refinements that could enhance performance beyond that of the HBS designs evaluated here. For example, a dual feedback HBS employing both HIF1α and HIF2α feedback could achieve higher sustained output over time. At short time scales, this boost could be driven by ectopic HIF1α (as observed in the HIF1α feedback HBS), and when HIF1α is degraded by HAF in long-term hypoxia, output gene expression could be maintained by ectopic HIF2α. A distinct potential strategy is to use HAF feedback or dual HAF/HIF2α feedback to enhance HIF2α-dependent HBS regulation in short-term hypoxia (increased HAF would enable additional production of the HAF-HIF2α complex), the latter of which could help maintain long-term output gene expression despite decrease in HIF1α in sustained hypoxia. Since HIF2α expression is generally a marker of poor cancer prognosis (5,16,18), a biosensor that rewires HIF2α-dependent signaling to induce a therapeutic output gene could be promising for applications that deploy HBSs in cancer cells, such as oncolytic viral therapy. Each of these ideas can be implemented and evaluated to build on the foundation of our HBS designs.

Extending the biosensors and circuits developed here to new applications, including translational usages, motivates a series of avenues for further work. First, it is likely that HIF1α, HIF2α, and HAF expression levels and dynamics will vary by cell type and state, and these may impact biosensor performance in ways that motivate application-directed tuning. Use of these biosensors for translational work such as with gene and cells therapies will necessitate distinct directions of inquiry. An advantage of the circuits reported here is that the transgenes encode native proteins, which limits concerns of immunogenicity. While these ectopic factors can target and modulate expression of endogenous genes, they would generally be expressed/stabilized alongside corresponding endogenous regulators. Nonetheless, ectopic expression of hypoxia regulators might impact cell state in ways that impact performance of a therapeutic. HIF1α is critical for survival of T cells (63), cytokine production and target cell killing by natural killer cells following activation (64), and regulation of T cell effector responses in the tumor microenvironment (65), but it is not clear how the mechanisms underlying these HIF1α regulatory modes could be affected by a modified HIF1α or HIF2α expression profile. Overall, the HBSs and insights into their function developed here can advance their use in fundamental research and therapeutic applications.

## DATA AVAILABILITY

Data are incorporated into the article and its online supplementary material (**Supplementary Data 3**).

## SUPPLEMENTARY DATA

Supplementary Data are available online. **Supplementary Data 1** contains all plasmid maps. **Supplementary Data 2** contains software. **Supplementary Data 3** contains all experimental data.

## AUTHOR CONTRIBUTIONS

K.S.D.: conceptualization, data curation, formal analysis, investigation, methodology, resources, software, visualization, writing – original draft, writing – review & editing. P.S.D.: conceptualization, data curation, formal analysis, investigation, methodology, project administration, resources, supervision, visualization, writing – original draft, writing – review & editing. J.D.B.: data curation, formal analysis, investigation, resources, visualization, writing – original draft, writing – review & editing. K.M.C. – investigation, resources, writing – review & editing. M.Y.O.: investigation, resources, writing – review & editing. H.I.E.: investigation, resources, writing – review & editing. B.D.L.: investigation, resources, writing – review & editing. K.J.Z.: investigation, resources, writing – review & editing. K.E.D.: methodology, writing – review & editing. J.J.M.: methodology, supervision, writing – review & editing. J.N.L.: conceptualization, funding acquisition, project administration, supervision, writing – review & editing. Manuscript was reviewed and edited by all authors prior to submission.

## ACKNOWLEDGEMENTS

The SV40_min promoter construct was a gift from Yvonne Chen (31). pcDNA3 mHIF-1α MYC (P402A/P577A/N813A) (Addgene #44028) was a gift from Celeste Simon (35). Lenti dCAS-VP64_Blast was a gift from Feng Zhang (Addgene #61425) (36). PhiC31-Neo-ins-5xTetO-pEF-H2B-Citrin-ins was a gift from Michael Elowitz (Addgene #78099) (37). pDsRed2-N1 was a gift from David Schaffer (University of California, Berkeley). pEBFP2-Nuc was a gift from Robert Campbell (Addgene #14893) (38). pU6-(BbsI)_CBh-Cas9-T2A-BFP-P2A-Ad4E4orf6 (Addgene #64220; referred to as pPD782), pU6-sgRosa26-1_CBh-Cas9-T2A-BFP-P2A-Ad4E1B (Addgene #64219; referred to as pPD720), and pR26 CAG/GFP Asc (Addgene #74285) were gifts from Ralf Kuehn (46,47). The pPart series vectors, pTU series vectors, pLink series vectors, destination vectors, and HEK293FT-LP cell line were gifts from Ron Weiss (34). Supplementary Figure 9B was created in BioRender by P.S.D. (2024) (BioRender.com/m62t381m62t381).

