## Supplementary Information for "Engineered Feedback Employing Natural Hypoxia-Responsive Factors Enhances Synthetic Hypoxia Biosensors"

**Contents:**
Supplementary Figures 1-16
Supplementary Tables 1-21
Supplementary Notes 1-7
References cited in supplementary information

### SUPPLEMENTARY FIGURES

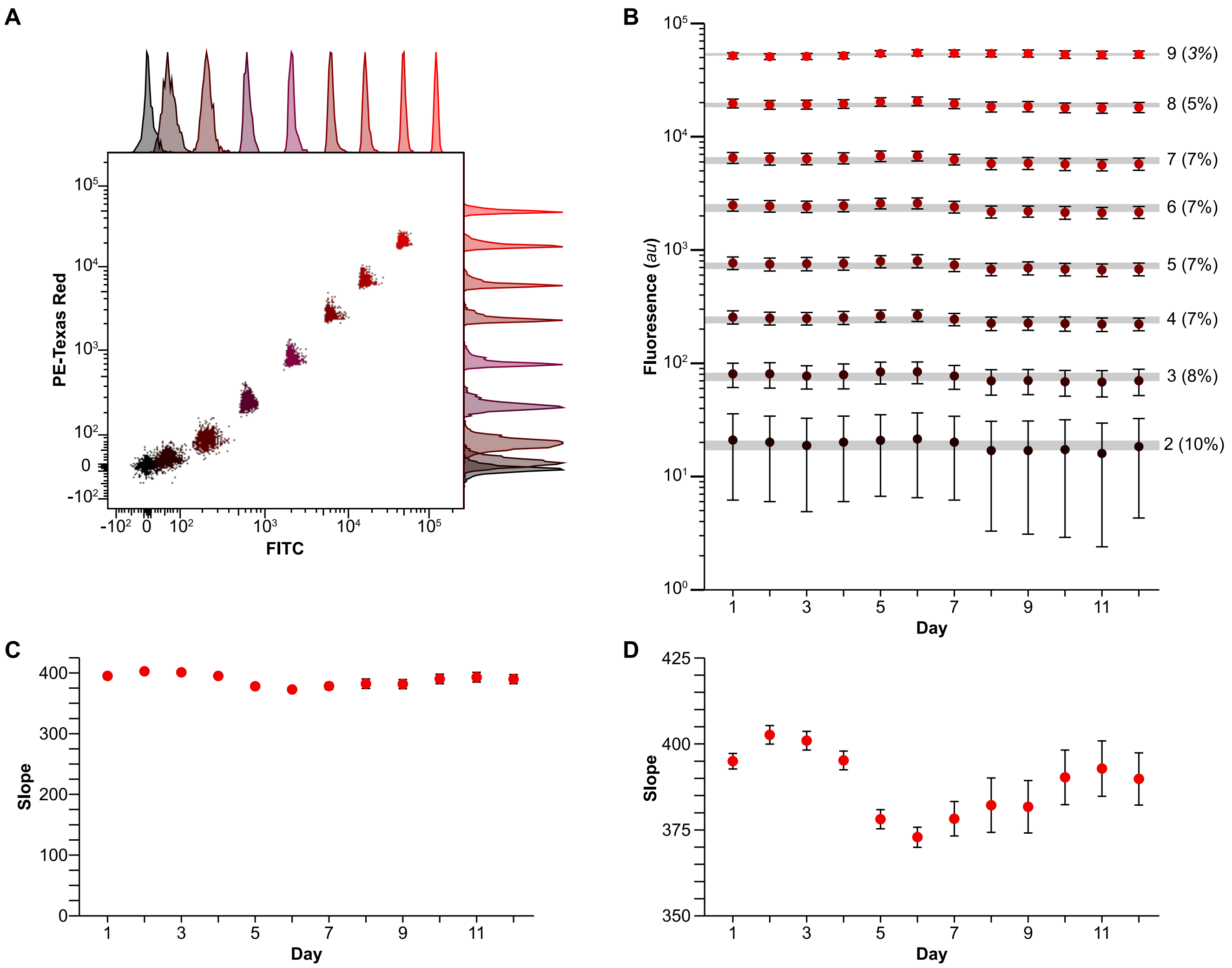

**Supplementary Figure 1. Conversion of arbitrary to standardized units and demonstration of possible impact on time course assays.** (**A**) Flow plot showing the 9 bead populations in the FITC and PE-Texas Red Channels. (**B**) The fluorescence (in *au*) of the 9 bead populations over 12 days. Error bars represent the standard deviation of the beads in each population. (**C**) Slope of the linear regression between AU and PETR for each of the 12 consecutive days (**D**) The plot shown is C with axis set to non-0 number to better illustrate day-to-day variation in the calibration. Error bars in **C** and **D** calculated by linear regression

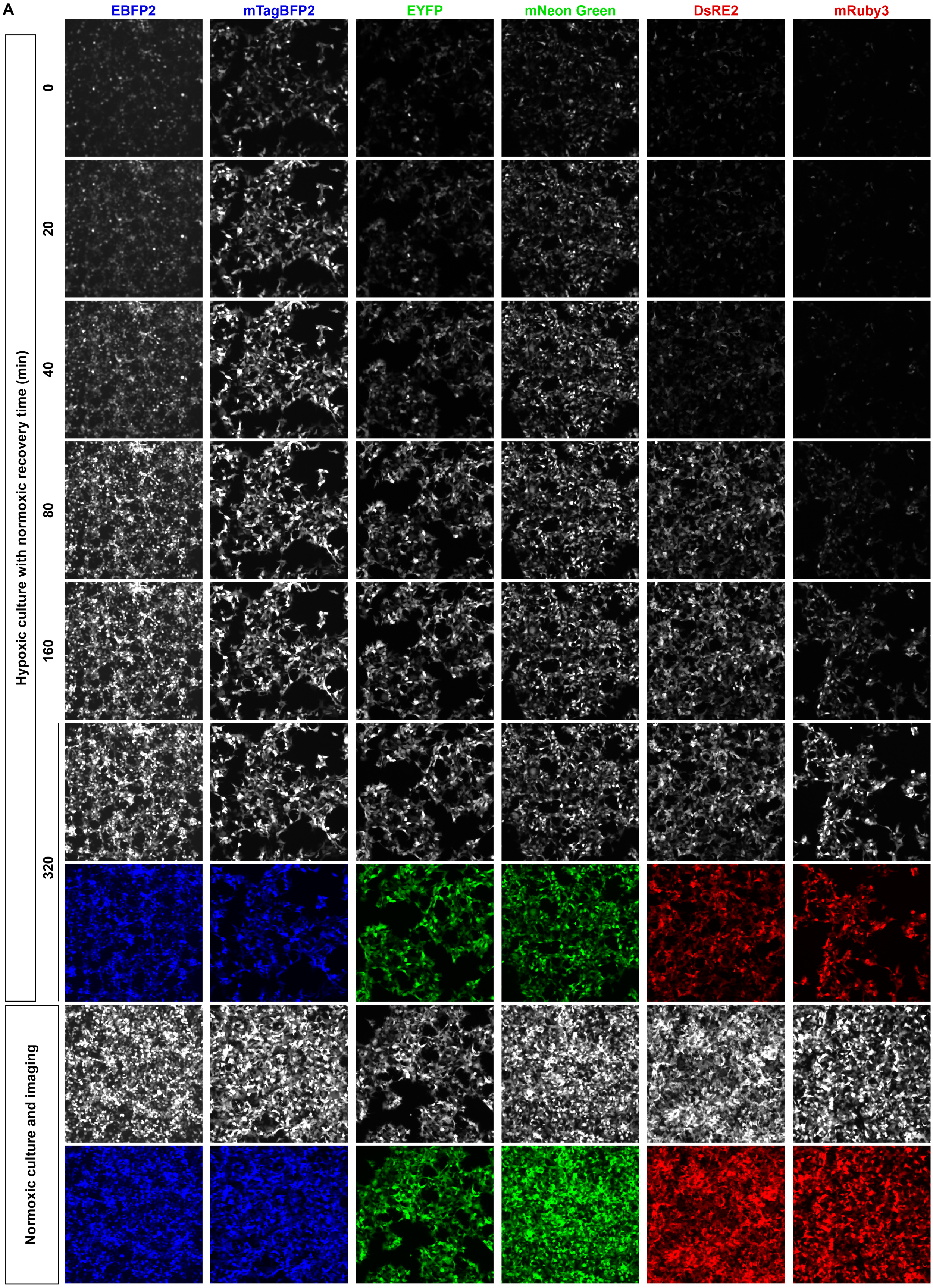
A
EBFP2
mTagBFP2
EYFP
mNeon Green
DsRE2
mRuby3
Hypoxic culture with normoxic recovery time (min)
0
20
40
80
160
320
Normoxic culture and imaging

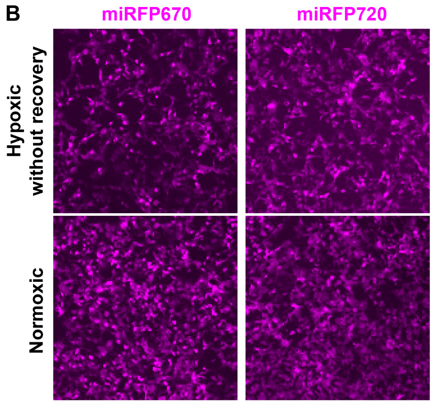

**Supplementary Figure 2. Oxidation rates of fluorescent proteins vary based on the fluorophore**. (**A**) Cells transfected with fluorescent protein expressing plasmids were cultured in hypoxic conditions and then serially imaged in a microscope in normoxic conditions, at 37˚C and 5% CO2; these cells were compared to cells that were cultured continuously in normoxic conditions after transfection. (**B**) Cells transfected with fluorescent protein expressing plasmids were cultured in hypoxic conditions and then immediately imaged and compared to cells that were cultured continuously in normoxic conditions after transfection. This plot represents the full data set from this experiment, of which a representative sample is shown in **Figure 1C**.

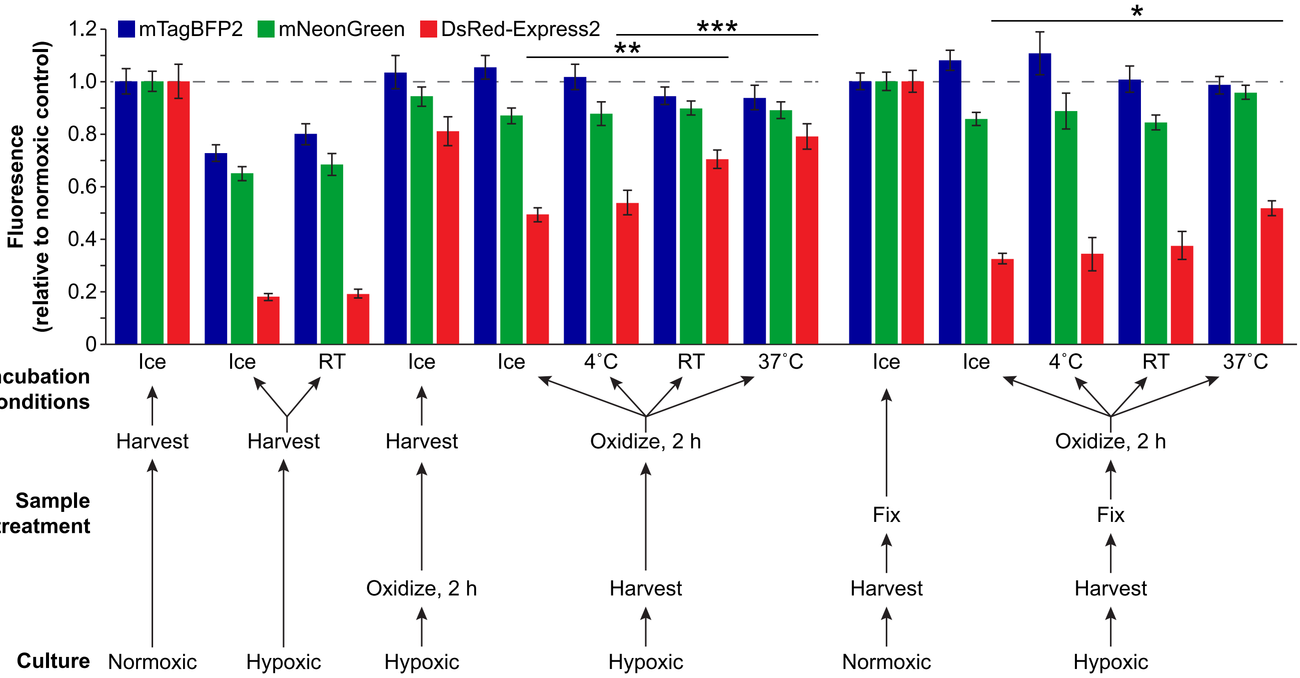

**Supplementary Figure 3. Comparison of methodologies for oxidizing fluorescent proteins to maturation after hypoxic culture.** After hypoxic or normoxic culture, cells transfected with fluorescent protein-expressing plasmids were analyzed after being oxidized under various conditions as indicated or not oxidized (by proceeding directly to flow cytometry after harvest and sample processing). The fluorescence values in each channel were normalized to the fluorescence of cells that had been continuously cultured under normoxia after transfection. Non-fixed and fixed samples were normalized separately. For samples oxidized at different temperatures, significant differences relative to the hypoxic condition incubated on ice are identified for each fluorescent protein (1-way ANOVA for each fluorescent protein, * $P < 0.05$, ** $P < 0.01$, *** $P < 0.001$). Outcomes from ANOVAs and Tukey's HSD tests are in **Supplementary Note 5**.

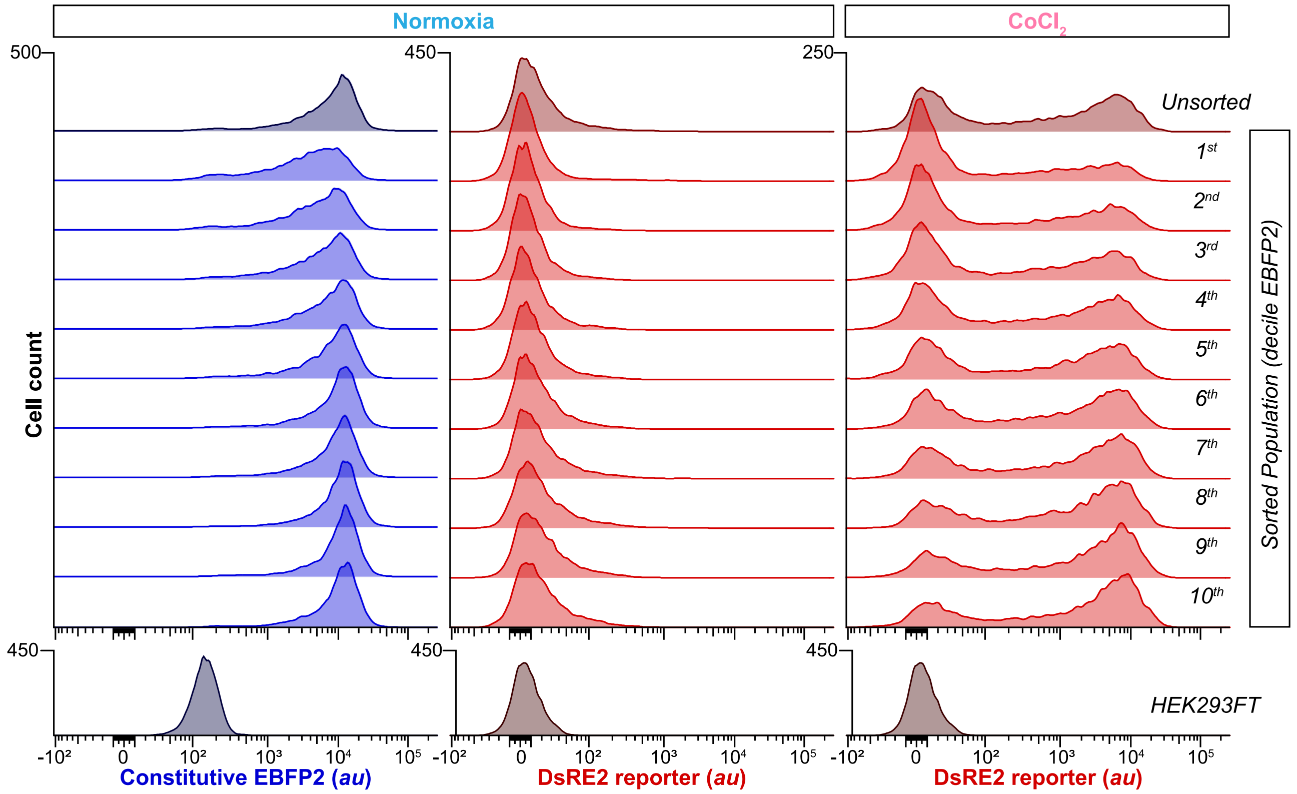

**Supplementary Figure 4. Representative flow cytometry plots corresponding to the experiment in Figure 2E.** The 10 populations of HEK-293FT with HBS integrated into the LP, which were sorted on the level of EBFP2 expression), were cultured with and without oxygen for 24 h. Lines that were derived from cells having higher EBFP2 expression demonstrated higher DsRE2 reporter expression. For cells cultured in hypoxia, the increase in reporter expression was driven by a higher percentage of cells expressing DsRE2 rather than a shift in the mode DsRE2 of these cells.

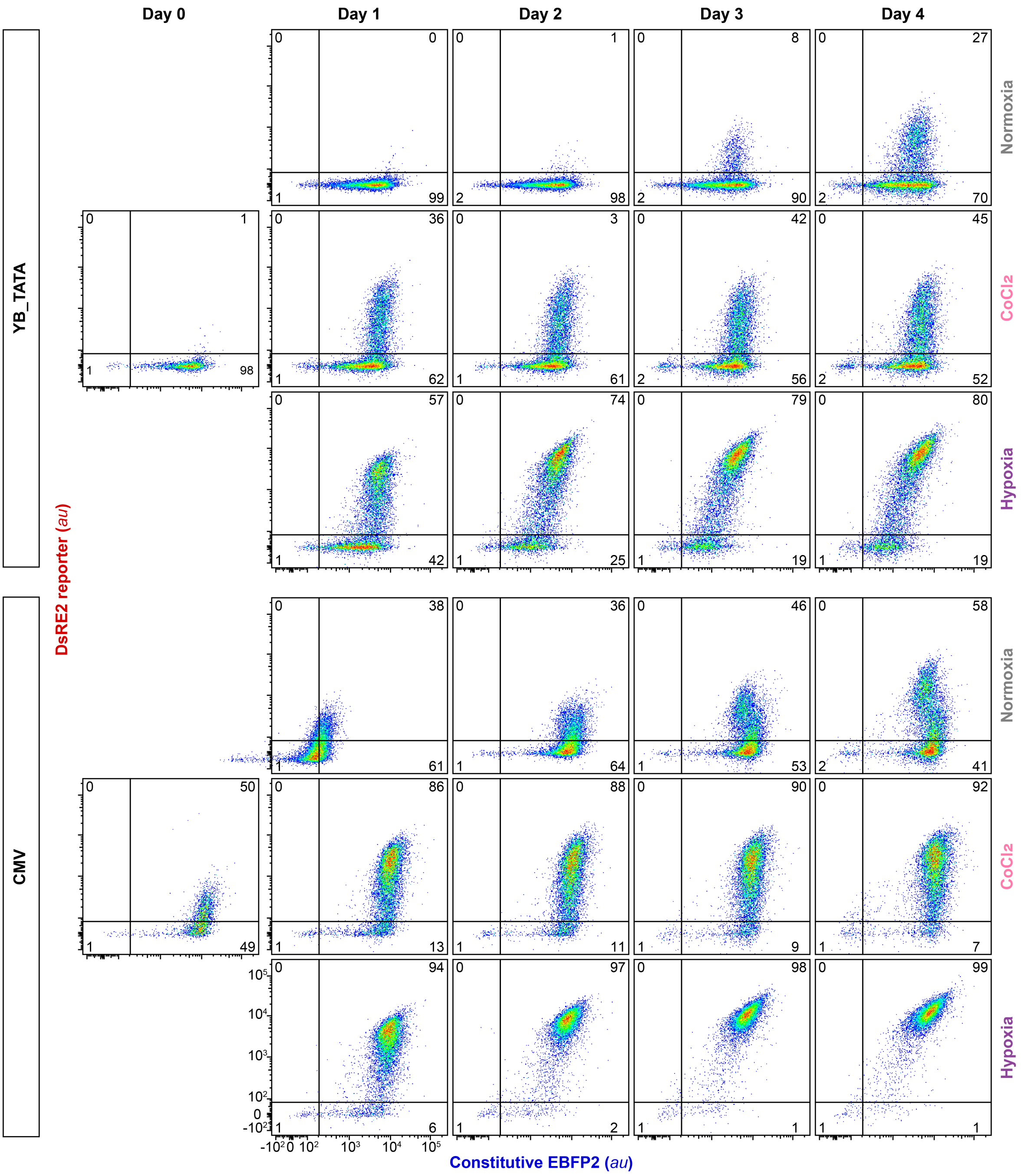

**Supplementary Figure 5. Representative flow cytometry plots corresponding to the experiment in Figure 3C.**

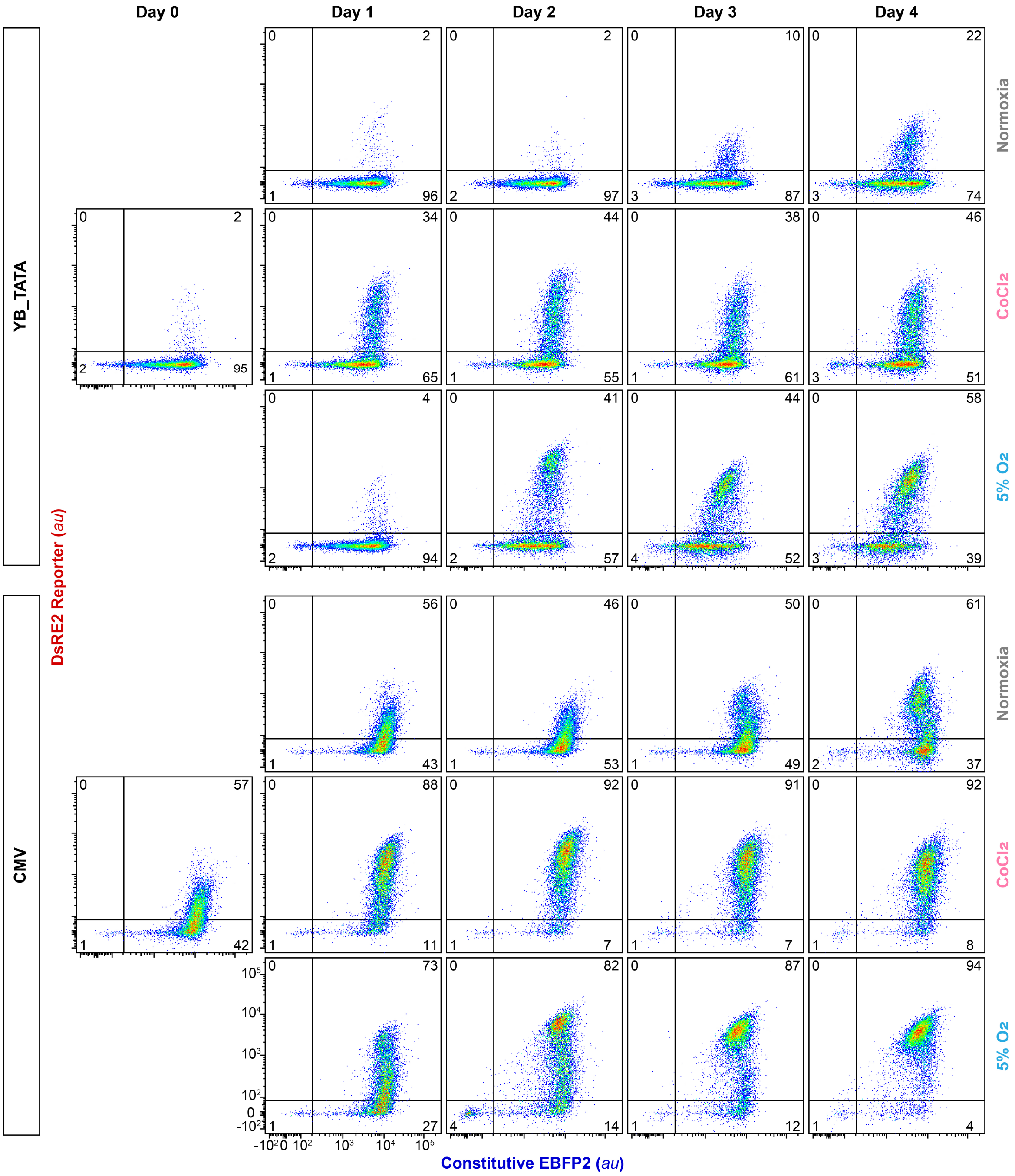

**Supplementary Figure 6. Representative flow cytometry plots corresponding to the experiment in Figure 3D.** Over the course of 5 d, some activation was seen for all cell lines tested. When comparing the flow cytometry plots from this experiment with the plots from the corresponding experiment during which cells were cultured under hypoxic conditions (**Supplementary Figure 5**), it appears that this activation started as a small subset of cells which gradually increased in number each day. When examined by microscopy (**Supplementary Figure 7**), this population of activated cells is noted at the center of large clumps of cells. This can be contrasted with the diffuse activation that is induced by $CoCl_2$. This activation is therefore likely a response to true hypoxic or near-hypoxic conditions, as the cells overgrow and consume oxygen faster than it can diffuse through the mass of cells. In other words, this is likely an experimental artifact of cell overgrowth rather than a direct activation of biosensors under physoxia.

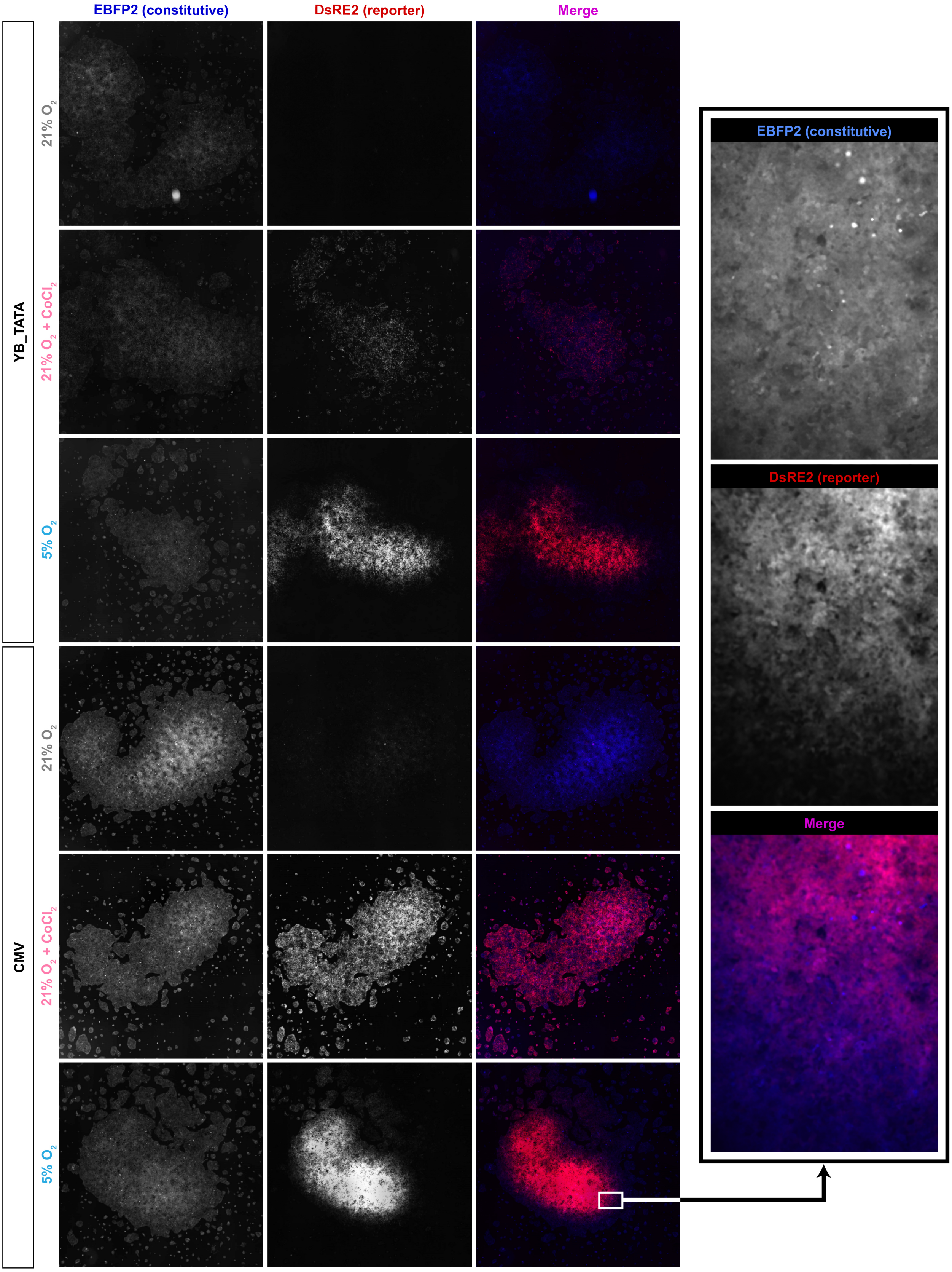

**Supplementary Figure 7. Flow C.** Images are representative samples from **Supplementary Figure 6**. Selected and sorted HEK293FT-LP containing BS with YB_TATA and CMV_min were cultured under the above conditions. Fluorescent micrographs of representative samples demonstrate stronger activation of the HBS at the center of clumps of cells in physoxic culture, contrasting with diffuse activation with $CoCl_2$. Inset is at higher magnification and rotated 90 degrees.

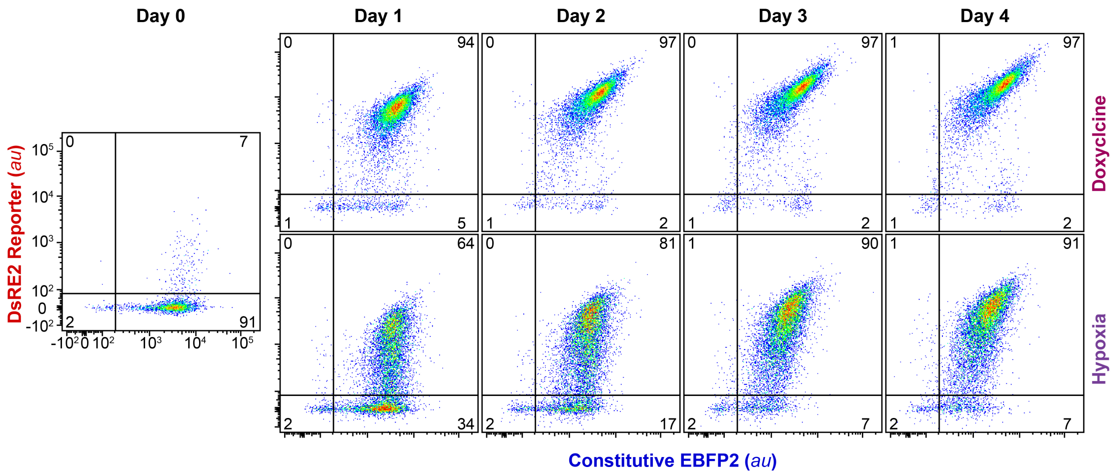

**Supplementary Figure 8. Representative flow cytometry plots corresponding to the experiment in Figure 4**

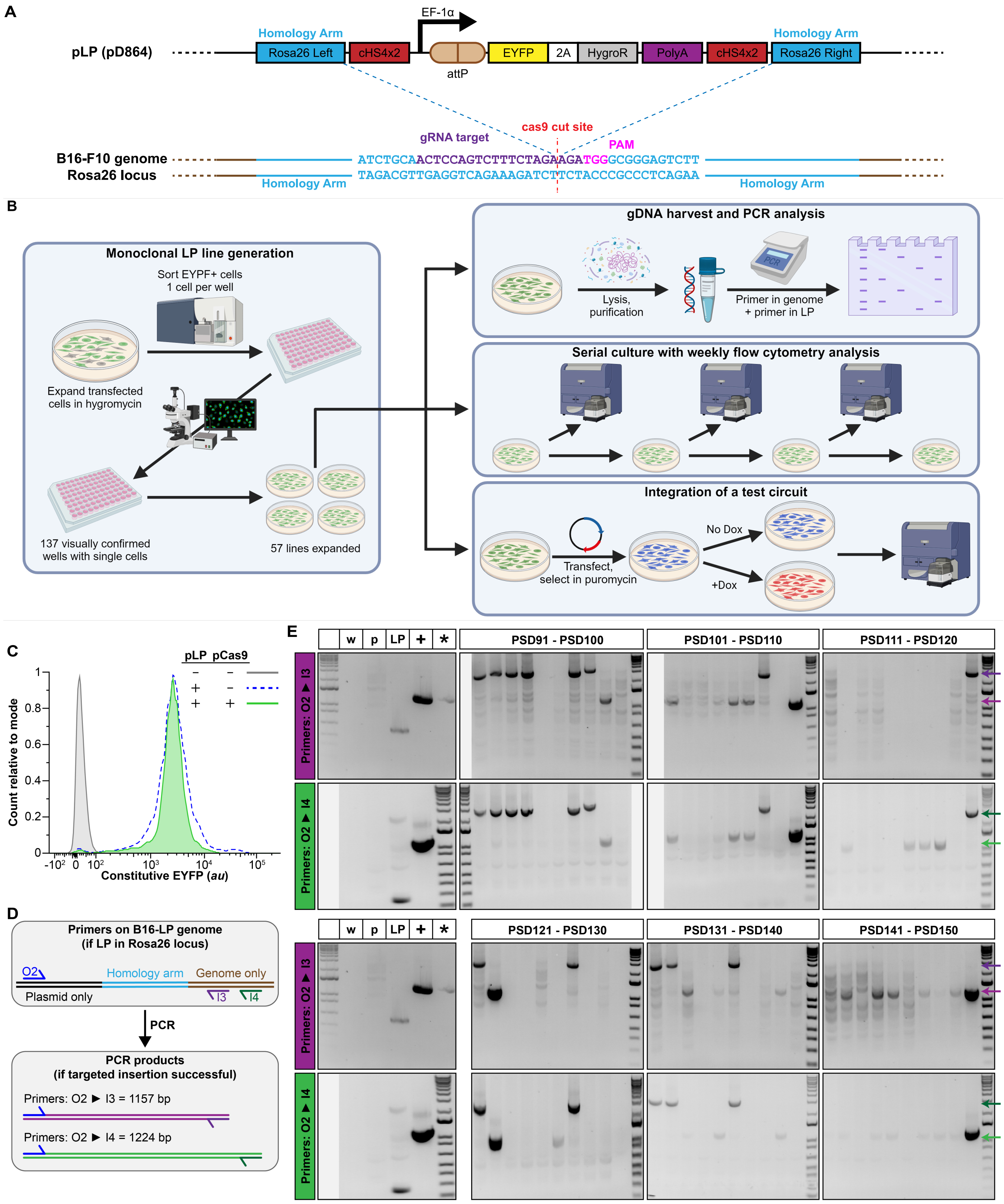
A
pLP (pD864)
Homology Arm
Rosa26 Left
cHS4x2
EF-1α
attP
EYFP
2A
HygroR
PolyA
cHS4x2
Homology Arm
Rosa26 Right
gRNA target
cas9 cut site
PAM
B16-F10 genome
Rosa26 locus
ATCTGCAACTCCAGTCTTTCTAGAAGATGGGCGGGAGTCTT
TAGACGTTGAGGTCAGAAAGATCTTCTACCCGCCCTCAGAA
Homology Arm
Homology Arm
B
Monoclonal LP line generation
Sort EYPF+ cells
1 cell per well
Expand transfected
cells in hygromycin
137 visually confirmed
wells with single cells
57 lines expanded
gDNA harvest and PCR analysis
Lysis,
purification
Primer in genome
+ primer in LP
Serial culture with weekly flow cytometry analysis
Integration of a test circuit
Transfect,
select in puromycin
No Dox
+Dox
C
pLP pCas9
Count relative to mode
Constitutive EYFP (au)
D
Primers on B16-LP genome
(if LP in Rosa26 locus)
O2
Homology arm
Genome only
Plasmid only
I3
I4
PCR
PCR products
(if targeted insertion successful)
Primers: O2 ► I3 = 1157 bp
Primers: O2 ► I4 = 1224 bp
E
w
p
LP
+
*
PSD91 - PSD100
PSD101 - PSD110
PSD111 - PSD120
Primers: O2 ► I3
Primers: O2 ► I4
PSD121 - PSD130
PSD131 - PSD140
PSD141 - PSD150

**Supplementary Figure 9. Generation of a B16F10-LP line**.

(**A**) Schematic depicting Cas9 mediated insertion of a landing pad into the Rosa26 locus of the B16F10 cell line. pLP is used as a template for HDR. (**B**) Workflow for generation and validation of the B16F10-LP line. (**C**) Initially two polyclonal cell lines were generated by transfection of pLP with or without the Cas9 expressing plasmid into the B16F10 line. Flow cytometry analysis after selection of these lines by culture in media containing hygromycin demonstrates the polyclonal line generated with Cas9 expression has a narrower distribution of EYFP expression. This likely indicates successful targeted integration in this polyclonal line, versus the polyclonal cell line generated by transfection of pLP without Cas9, in which integration of pLP was likely random. (**D**) Schematic depicting primer design for detection of monoclonal lines with successful integration of the LP into the Rosa26 locus. A forward primer is located on a DNA sequence only existing on the plasmid, while a reverse primer is located on a DNA sequence only existing in the genome. Two sets of primers, sharing a common forward (outward) primer and with different reverse (inward) primers, are shown. The primers span the homology arm and should only produce a PCR product of the appropriate size if the polymerase can proceed through the homology arm. (**E**) Results of the PCR reactions illustrated in D on genomic DNA from 59 cell lines (line PSD101 died during expansion). Reactions with primer set O2/I3 and O2/I4 were conducted independently. The image comprising the control wells for each set of primers are duplicated on each line for ease of visualization. Controls include (from left to right): w = water used instead of genomic DNA (should be negative); p = genomic DNA from the parental B16F10 line (should be negative), LP = purified pLP plasmid (should be negative), + = purified plasmid containing the target sequence (should be positive), * = the polyclonal cell line generated by transfection of pLP and pCas9 from which the monoclonal lines were derived (would be positive if integration occurred in some portion of cells; for the O2/I4 reaction, this control is replaced by a ladder). The anticipated band size is shown by light purple and light green arrows. Sanger sequencing of the appropriately sized band (~1200 bp) from the O2/I3 reaction revealed the anticipated sequence. An extra, high molecular weight band was noted for some cell lines (indicated by dark purple and dark green arrows). Sequencing of this band revealed the *KanR* gene that was located on the pLP backbone, suggesting that these lines were derived from integration of the plasmid into the genome, rather than Cas9 mediated dsDNA break and subsequent HDR from the pLP template. Solid grey background is used for lanes without data. Cell lines chosen for further analysis in **Supplementary Figure 10** are demarcated with their line numbers at the bottom of the gel.

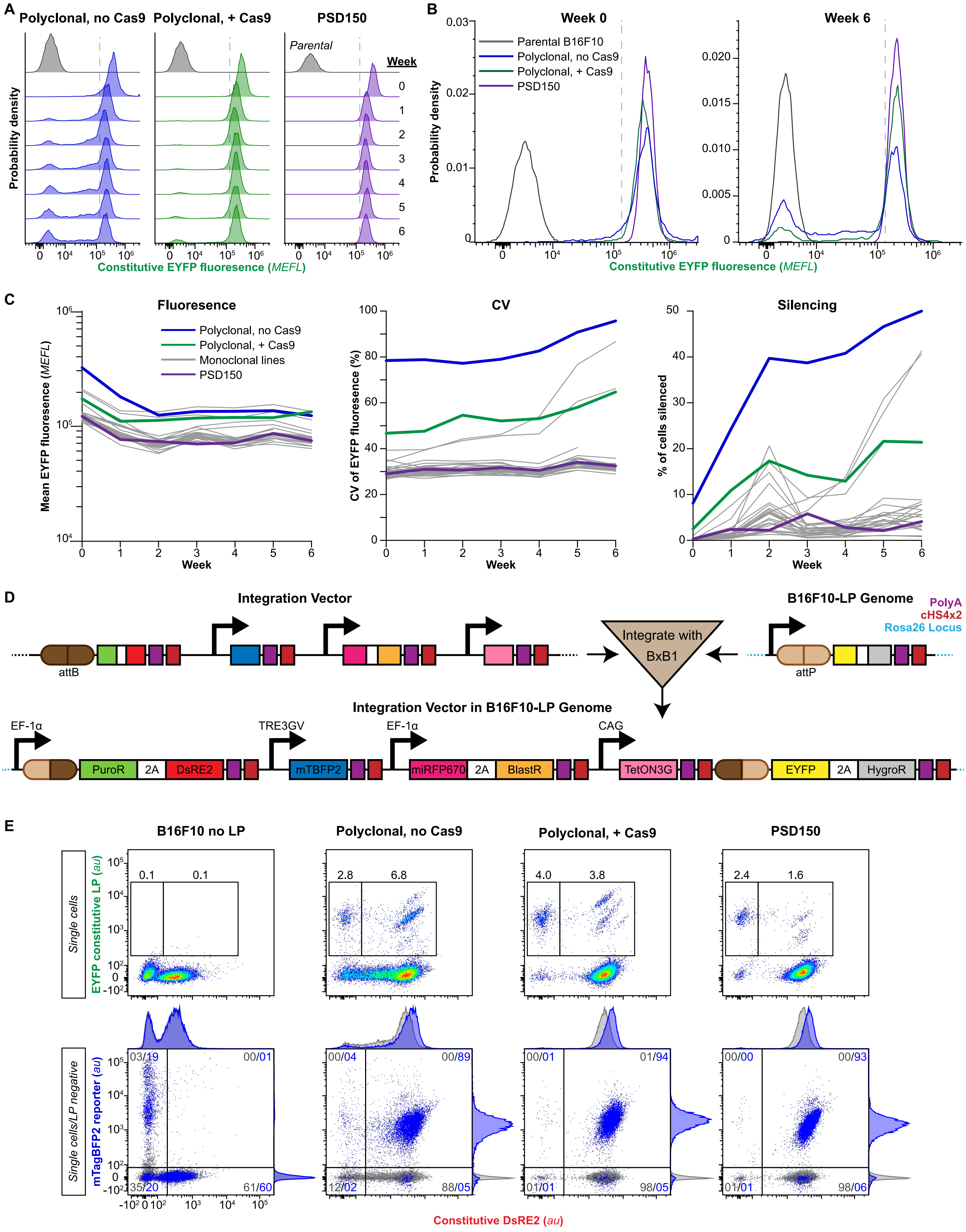

A
Polyclonal, no Cas9
Polyclonal, + Cas9
PSD150
Parental
Week
Probability density
Constitutive EYFP fluorescence (MEFL)
B
Week 0
Week 6
Parental B16F10
Polyclonal, no Cas9
Polyclonal, + Cas9
PSD150
Probability density
Constitutive EYFP fluorescence (MEFL)
C
Fluoresence
CV
Silencing
Polyclonal, no Cas9
Polyclonal, + Cas9
Monoclonal lines
PSD150
Mean EYFP fluorescence (MEFL)
CV of EYFP fluorescence (%)
% of cells silenced
Week
D
Integration Vector
B16F10-LP Genome
PolyA
cHS4x2
Rosa26 Locus
attB
attP
Integrate with BxB1
Integration Vector in B16F10-LP Genome
EF-1α
TRE3GV
EF-1α
CAG
PuroR
2A
DsRE2
mTBFP2
miRFP670
2A
BlastR
TetON3G
EYFP
2A
HygroR
E
B16F10 no LP
Polyclonal, no Cas9
Polyclonal, + Cas9
PSD150
Single cells
EYFP constitutive LP (au)
Single cells/LP negative
mTagBFP2 reporter (au)
Constitutive DsRE2 (au)

**Supplementary Figure 10. Validation of a B16F10-LP line**.

**(A-C)** Comparison of constitutive EYFP fluorescence over a 6-week culture period, with weekly flow cytometry analysis. Over the 6-week period, the polyclonal population generated by random integration (blue lines) largely underwent silencing, indicating integration into non-safe harbor loci. In contrast the polyclonal population generated by targeted integration with Cas9 contained fewer silenced cell, indicating successful integration into a safe harbor locus, presumably the targeted Rosa26 locus. PSD150 is the B16F10-LP line ultimately chosen as the LP line for all further experiments (purple lines). This line demonstrated a very low percentage of silencing and maintained a tight CV throughout the experiment. The other candidate cell lines (as demonstrated by asterisks in **Supplementary Figure 9**) are shown in grey. Silencing is calculated as the percentage of cells with lower EYFP fluorescence than the set value (grey dashed line). This value was calculated as the EYFP fluorescence (in MEFLs) 2 SD below the mean for the targeted integration polyclonal population (green lines). CV = coefficient of variance, which is the ratio of the standard deviation to the mean, where lower values signify populations with lower variation. **(D)** Schematic depicting integration of a test circuit into the LP in the B16F10-LP cell line utilizing the BxB1 recombinase, which splices the attP site of the Integration Vector (IV) into the attB site of the LP. Given the absence of a constitutive promoter in the IV, cells only become resistant to puromycin and fluorescent red when the attP site is integrated in front of a constitutive promoter, presumably the one upstream of the attB site in the LP. The displacement of the EYFP gene from this promoter renders the cells no longer fluorescent in the EYFP (FITC) channel. Cells also constitutively express an infrared fluorescent protein and a gene for blasticidin resistance, constitutively regardless of whether the IV is integrated or remains exogenous. Finally, the addition of doxycycline induces expression of mTagBFP2 from the tet-responsive TRE3GV promoter. **(E)** Comparison of B16F10-LP lines with the IV shown in **(C)** integrated, after selection with puromycin and blasticidin. This experiment includes a line in which the IV was transfected into parental B16F10 cells with no LP. Interestingly, this line produced 2 populations of cells, of which the dim constitutive red population demonstrated doxycycline-inducible mTagBFP2 expression, while the slightly brighter constitutive population only contained rare cells. In contrast to this, cells containing a landing pad showed markedly brighter constitutive DsRE2 expression. Among these lines, those generated by targeted integration of the LP with Cas9 into the Rosa26 locus showed more homogenous levels of the constitutive DsRE2. Additionally, it appears that the B16F10 LP line readily integrates plasmids into its genome—as demonstrated by the population of cells retaining EYFP fluorescence after attempted integration of the IV into the LP. This reinforces the need for the dual antibiotic selection and subsequent sorting by flow cytometry, for the cells that no longer have an intact LP (EYFP negative) and those that have the highest levels of constitutive protein expression (here, DsRE2).

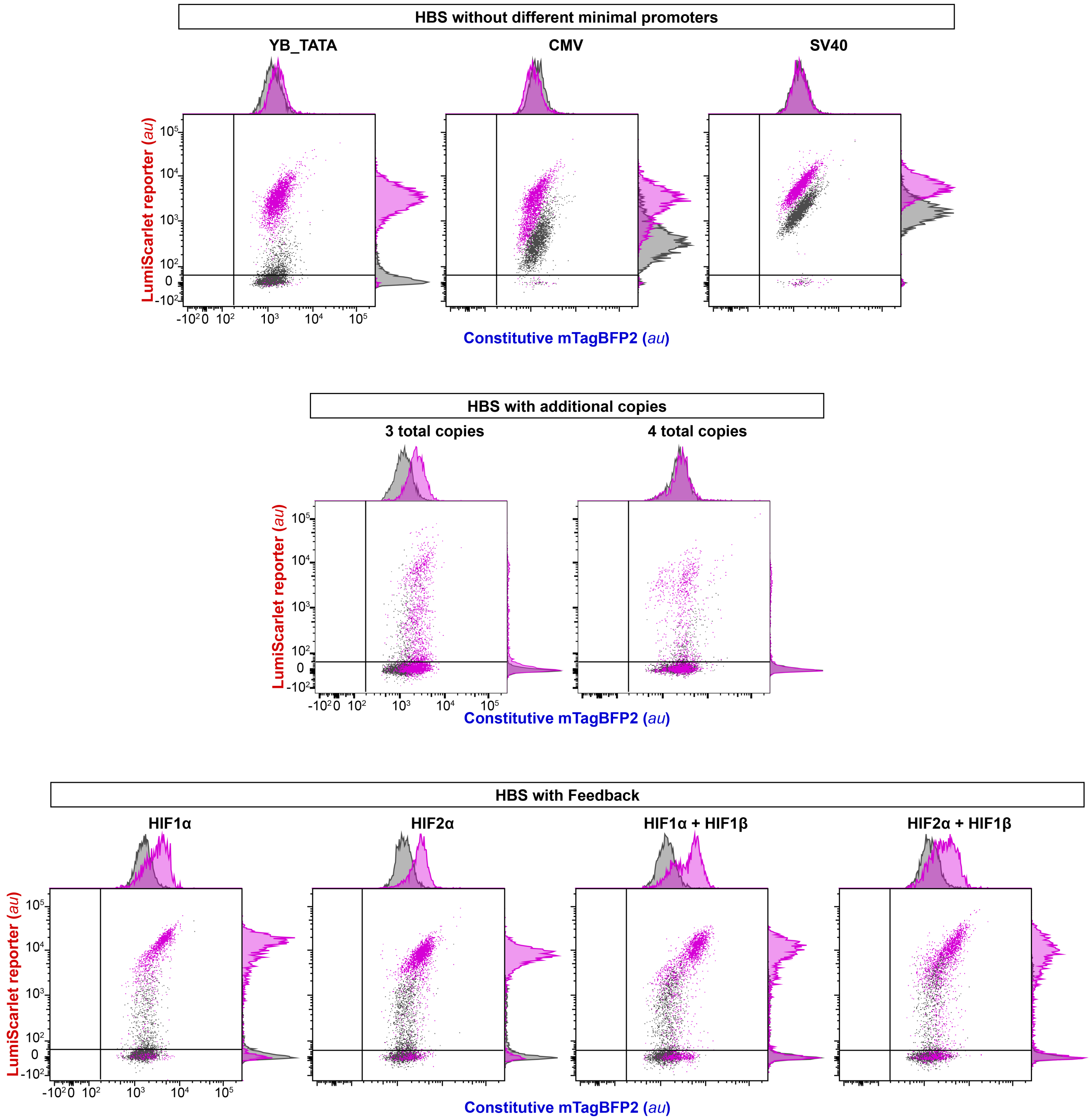

**Supplementary Figure 11. Representative flow cytometry plots corresponding to the experiment in Figure 5B**

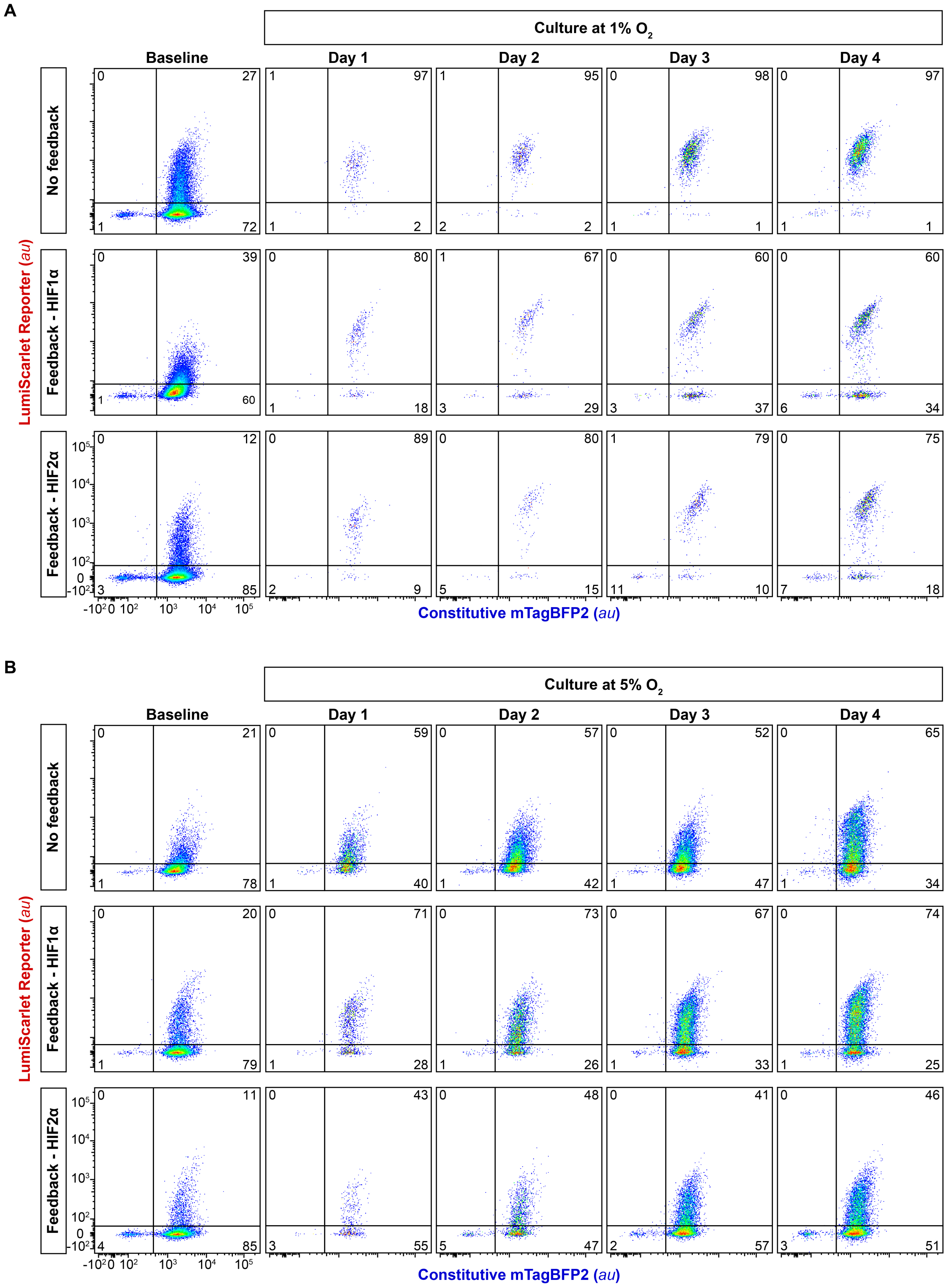

**Supplementary Figure 12. Representative flow cytometry plots corresponding to the experiments in Figure 5C and 5D.**

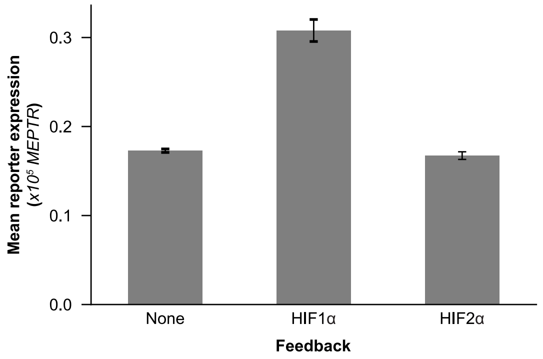

**Supplementary Figure 13. Comparison of the effects of feedback circuits on background signals.** Data correspond to the baselines shown in **Figure 5D**.

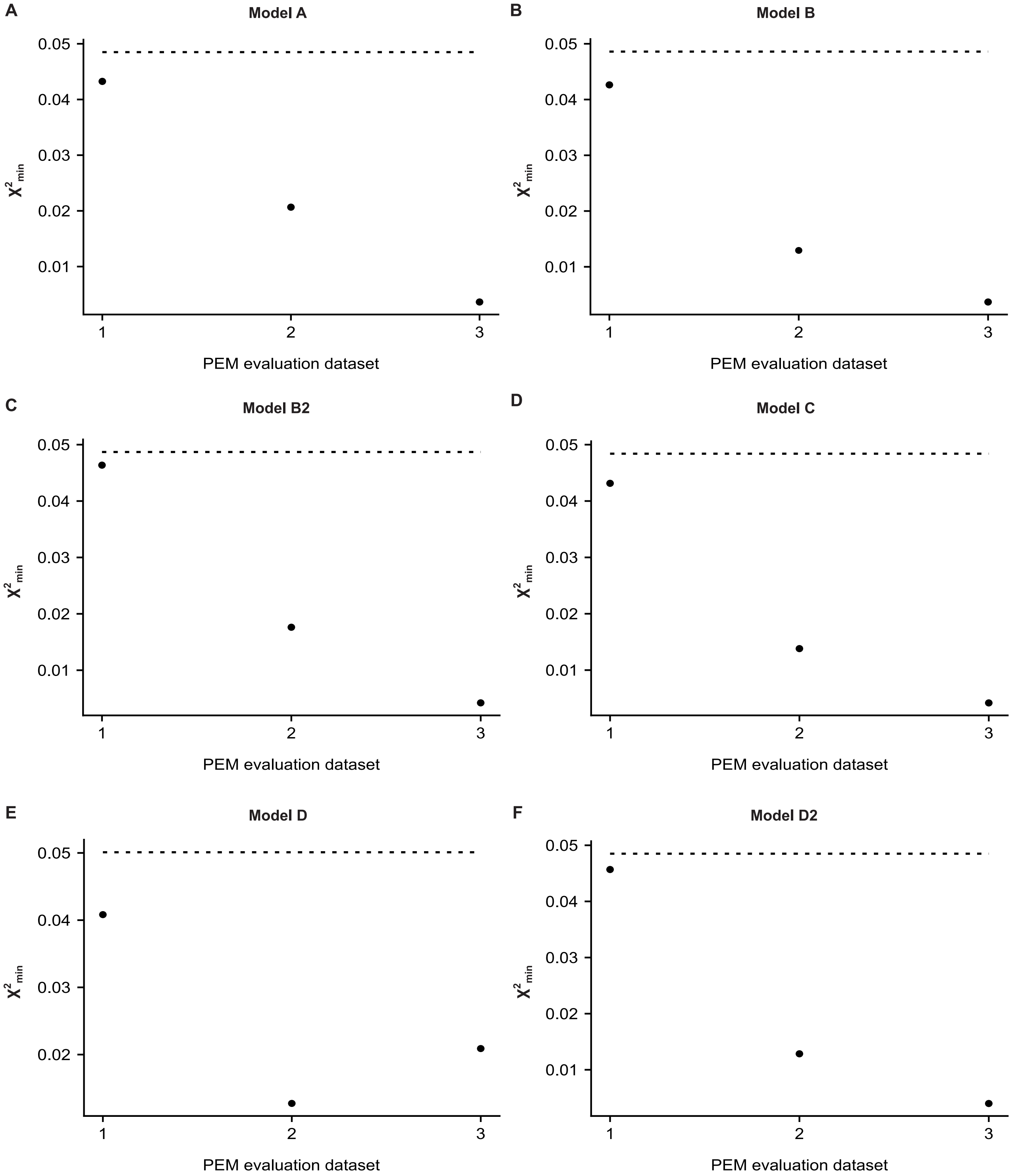

**Supplementary Figure 14. Parameter estimation method (PEM) evaluation results for each model version.** For each model, 3 PEM evaluation datasets with noise representative of the training data error were generated following the method described by Dray et al (1) with $n_{search}$= 1000 and $n_{init}$= 100, as was used in the PEM (**MATERIALS AND METHODS**). The PEM was then used to estimate parameters. Plots show the $\chi^2$ between simulated values and the PEM evaluation dataset for parameter sets yielding the minimum $\chi^2$ value. The dashed line indicates the $\chi^2$PEM evaluation criterion, which was defined as the maximum across the 3 datasets of the $\chi^2$between the PEM evaluation dataset with vs. without noise added.

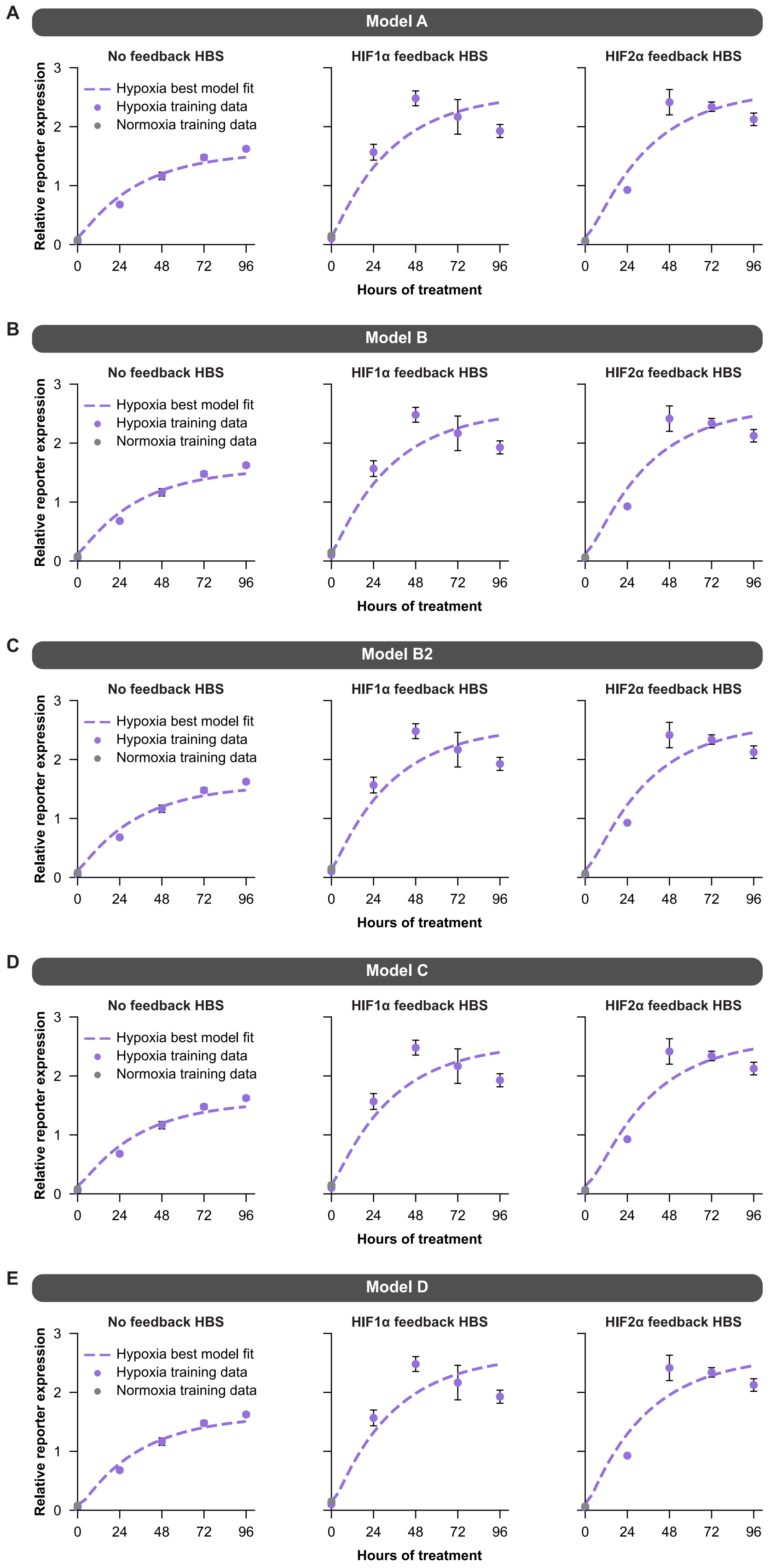

**Supplementary Figure 15. Analysis of sub-optimal candidate models.** Models A-D shown with best fit to training data for the 3 HBSs. Table 1 indicates which modeling objectives were met for each model.

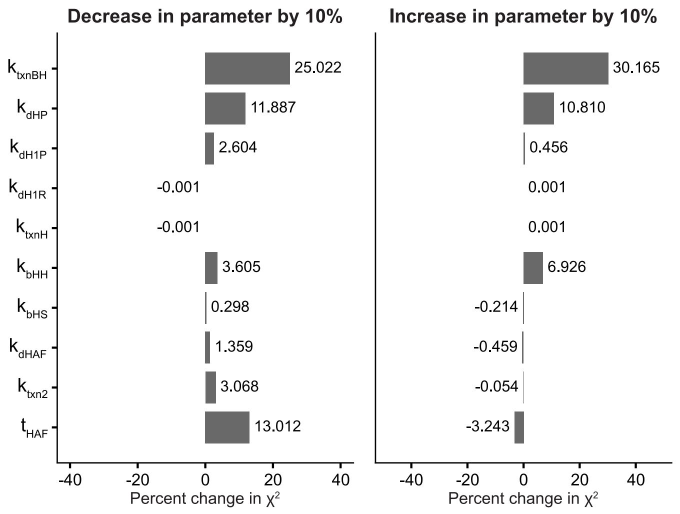

**Supplementary Figure 16. Parameter sensitivity analysis using $\chi^2$ as a metric.** Percent change in $\chi^2$ when decreasing each parameter individually by 10% (left) or increasing each parameter individually by 10% (right), relative to the $\chi^2$ for the calibrated parameter set.

### SUPPLEMENTARY TABLES

#### Supplementary Table 1. Abbreviations

| Abbreviation | Full name |
|---|---|
| ANOVA | Analysis of variance |
| AU | Arbitrary units |
| BSA | Bovine serum albumin |
| CMV | Cytomegalovirus |
| DMEM | Dulbecco's Modified Eagle Medium |
| DNA | Deoxyribonucleic acid |
| Dox | Doxycycline |
| EBFP2 | Enhanced blue fluorescent protein 2 |
| EDTA | Ethylenediaminetetraacetic acid |
| EYFP | Enhanced yellow fluorescent protein |
| FACS | Fluorescence activated cell sorting |
| FBS | Fetal bovine serum |
| FSC-A | Area in the Forward Scatter channel |
| FSC-H | Height in the Forward Scatter channel |
| HBS | Hypoxia biosensor |
| HEPES | 4-(2-hydroxyethyl)-1-piperazineethanesulfonic acid |
| HIF | Hypoxia inducible factor |
| HRE | Hypoxia response element |
| HSD | Tukey's honest significance test |
| IV | Integration vectors |
| LP | Landing pad |
| MEFLs | Molecules of Equivalent Fluorescein |
| MEPB | Molecules of Equivalent Pacific Blue |
| MEPTRs | Molecules of Equivalent PE-Texas Red |
| MFI | Mean fluorescence intensity (in arbitrary units) |
| miRFP670 | monomeric infrared fluorescent protein (with 670 nm emission) |
| miRFP720 | monomeric infrared fluorescent protein (with 720 nm emission) |
| NEB | New England Biolabs |
| PBS | Phosphate-buffered saline |
| RCP | Rainbow calibration particles |
| RNA | Ribonucleic acid |
| SDS | Sodium dodecyl sulfate |
| SEM | Standard error of the mean |
| SNR | Signal-to-noise ratio |
| SSC-A | Area in the Side Scatter channel |
| SV40 | Simian vacuolating virus 40 |
| TBS | Tris-buffered saline |
| TBST | Tris-buffered saline with Tween |
| TE | Tris EDTA buffer |
| TME | Tumor microenvironment |
| TUPV | Transcription unit positioning vector |
| URCP | Ultra-rainbow calibration particles |

**Supplementary Table 2. mMoClo Integration Vector construction[1]**

| Plasmid[2] | Backbone | TUPV1 | TUPV2 | TUPV3 | TUPV4 | TUPV5 | TUPV6 |
|---|---|---|---|---|---|---|---|
| **pPD787** | pPD630 | pPD774 | pPD852 | pPD612 | | | |
| **pPD788** | pPD630 | pPD775 | pPD852 | pPD612 | | | |
| **pPD790** | pPD630 | pPD774 | pPD769 | pPD770 | pPD854 | pPD614 | |
| **pPD793** | pPD630 | pPD766 | pPD847 | pPD853 | pPD613 | | |
| **pPD1401** | pPD630 | pPD1291 | pPD1151 | | | | |
| **pPD1402** | pPD630 | pPD1046 | pPD1360 | pPD1361 | pPD1153 | | |
| **pPD1403** | pPD630 | pPD1354 | pPD1360 | pPD1361 | pPD1153 | | |
| **pPD1404** | pPD630 | pPD1049 | pPD1360 | pPD1361 | pPD1153 | | |
| **pPD1411** | pPD1178 | pPD1352 | pPD1292 | pPD1152 | | | |
| **pPD1412** | pPD1178 | pPD1351 | pPD1292 | pPD1152 | | | |
| **pPD1413** | pPD1178 | pPD1350 | pPD1292 | pPD1152 | | | |
| **pPD1417** | pPD1178 | pPD1350 | pPD1357 | pPD1285 | pPD1294 | pPD1154 | |
| **pPD1418** | pPD1178 | pPD1350 | pPD1357 | pPD1358 | pPD1359 | pPD1295 | pPD1155 |
| **pPD1431** | pPD1178 | pPD1350 | pPD1271 | pPD1293 | pPD1153 | | |
| **pPD1432** | pPD1178 | pPD1350 | pPD1272 | pPD1293 | pPD1153 | | |
| **pPD1433** | pPD1178 | pPD1350 | pPD1271 | pPD1284 | pPD1294 | pPD1154 | |
| **pPD1434** | pPD1178 | pPD1350 | pPD1272 | pPD1284 | pPD1294 | pPD1154 | |
| **pMO18** | pPD1060 | pPD1043 | pPD1031 | pPD770 | pPD613 | | |

[1]See Supplementary Tables 11-13 for further details of the vectors used for assembly.
[2]See Supplementary Table 13 for further details of the constructed Integration Vectors.

**Supplementary Table 3. Filters for microscopy**

| Name | Product No. | Fluorescent Proteins | Filter 1 | Filter 2 | Filter3 |
|---|---|---|---|---|---|
| ET-EBFP2/Coumarin/ Attenuated DAPI | 49021 | EBFP2, mTagBFP2 | ET405/20x | ET460/50m | T425lpxr-UF1 |
| ET - EYFP | 49003 | EYFP, mNeonGreen | ET500/20x | ET535/30m | T515lp-UF1 |
| ET - DSRed (TRITC/Cy3) | 49005 | DsRed-Express2, LumiScarlet, mRuby3 | ET545/30x | ET620/60m | T570lp-UF1 |
| ET Cy5.5 | 49022 | miRFP670, miRFP720 | ET650/45x | ET720/60m | T685lpxr-UF1 |

**Supplementary Table 4. Filters for analytical flow cytometry**

| Instrument | Fluorescent proteins | Parameter/ Channel name | Excitation laser | Filter set |
|---|---|---|---|---|
| BD LSRFortessa | EBFP2, mTagBFP2 | Pacific Blue | Violet, 405 nm | 450/50 |
| | EYFP, mNeonGreen | FITC | Blue, 488 nm | 505LP, 530/30 |
| | DsRed-Express2, LumiScarlet | PE-Texas Red | Light Green, 552 nm | 600LP, 610/20 |

**Supplementary Table 5. Filters for flow cytometry sorting**

| Instrument | Fluorescent proteins | Parameter/ Channel name | Excitation laser | Filter set |
|---|---|---|---|---|
| BD FACS Aria | EBFP2, mTagBFP2 | Pacific Blue | Violet 407 nm | 450/50 |
| | EYFP | FITC | Blue 488 nm | 505LP, 525/30 |

**Supplementary Table 6. Select reagents for Golden Gate reactions**

| Reagent | Manufacturer | Product ID |
|---|---|---|
| BpiI-FD | ThermoFisher, MA, USA | FD1014 |
| BSA | Amresco, OH, USA | 0332 |
| T4 DNA Ligase | New England BioLabs, MA, USA | M202 |
| T4 ligase buffer | New England BioLabs, MA, USA | B0202 |

**Supplementary Table 7. Select reagents for cell culture**

| Reagent | Manufacturer | Product ID |
|---|---|---|
| HEPES | Sigma, MA, USA | H3375 |
| Blasticidin | Alfa Aesar/ThermoFisher, MA, USA | J61883 |
| $CoCl_2$ | Sigma-Aldrich, MO, USA | C-2644 |
| DMEM | ThermoFisher/Gibco, MA, USA | 31600-091 |
| Doxycycline | Sigma-Aldrich, MO, USA | D9891 |
| FBS | ThermoFisher/Gibco, MA, USA | 16140-071 |
| Gentamycin | ThermoFisher/Gibco, MA, USA | 15750060 |
| Hygromycin | EMD Millipore, MA, USA | 400052 |
| L-glutamine | ThermoFisher/Gibco, MA, USA | 25030-081 |
| Lipofectamine LTX with PLUS Reagent | ThermoFisher/Invitrogen, MA, USA | 15338100 |
| OptiMEM | ThermoFisher/Gibco, MA, USA | 31985062 |
| Penicillin-Streptomycin | ThermoFisher/Gibco, MA, USA | 15140122 |
| Phenol Red-free DMEM Powder | Sigma, MA, USA | D2902 |
| Puromycin | InvivoGen, CA, USA | ant-pr |
| Pyridoxine-HCl | Sigma, MA, USA | P6280 |
| Trypsin-EDTA | ThermoFisher/Gibco, MA, USA | 25300054 |
| Trypsin-EDTA (pheno-red free) | ThermoFisher/Gibco, MA, USA | 15400054 |

**Supplementary Table 8. Reagents for flow cytometer calibration**

| Reagent | Manufacturer | Product ID |
|---|---|---|
| RCP | Spherotech, IL, USA | RCP-30-5 |
| URCP | Spherotech, IL, USA | URCP-100-2H |

**Supplementary Table 9. Kits used in this study**

| Reagent | Manufacturer | Product ID |
|---|---|---|
| GeneJET Genomic DNA Purification Kit | ThermoFisher, MA, USA | K7021 |
| MycoAlert Mycoplasma Detection Kit | Lonza, Switzerland | LT07-318 |

**Supplementary Table 10. Instrumentation**

| Instrument | Manufacturer |
|---|---|
| All-in-One Fluorescence Microscope BZ-X800E | Keyence, IL, USA |
| BD LSRFortessa Special Order Research Product | BD Biosciences, NJ, USA |
| BD FACS Aria | BD Biosciences, NJ, USA |
| Heracell 150i Tri-Gas Incubator, 150L, custom ordered to go down to 1% $O_2$ | Thermo Scientific, MA, USA Fisher Scientific Cat No 13-998-037 |
| PreSens SDR SensorDish system | PreSens Precision Sensing, Germany |

**Supplementary Table 11. Plasmids obtained from external sources**

| Plasmid Name | Source | Reference |
|---|---|---|
| Bxb1 (codon optimized), (pPD610) | Ron Weiss, MIT | NAR (2) |
| Destination Vector (pPD630) | Ron Weiss, MIT | NAR (2) |
| lenti dCAS-VP64_Blast | Addgene | RRID: Addgene_61425 |
| p3'UTR | Ron Weiss, MIT | RRID: Addgene_139248 |
| p5'UTR | Ron Weiss, MIT | NAR (2) |
| pcDNA3 mHIF-1α MYC (P402A/P577A/N813A) | Addgene | RRID: Addgene_44028 |
| pDestination Vector | Ron Weiss, MIT | NAR (2) |
| pEBFP2-Nuc | Addgene | RRID: Addgene_14893 |
| pGene | Ron Weiss, MIT | NAR (2) |
| PhiC31-Neo-ins-5xTetO-pEF-H2B-Citrin-ins | Addgene | RRID: Addgene_78099 |
| pInsulator | Ron Weiss, MIT | NAR (2) |
| pLink2 (pPD612) | Ron Weiss, MIT | NAR (2) |
| pLink3 (pPD613) | Ron Weiss, MIT | NAR (2) |
| pLink4 (pPD614) | Ron Weiss, MIT | NAR (2) |
| pLVX-Tet3G | Clontech | Takara: 631187 |
| pLVX-TRE3G | Clontech | Takara: 631187 |
| pPolyA | Ron Weiss, MIT | NAR (2) |
| pPro | Ron Weiss, MIT | NAR (2) |
| pR26 CAG/GFP Asc | Addgene | RRID: Addgene_74285 |
| pTU1 | Ron Weiss, MIT | NAR (2) |
| pTU2 | Ron Weiss, MIT | NAR (2) |
| pU6-(BbsI)_CBh-Cas9-T2A-BFP-P2A-Ad4E4orf6 (pPD782) | Addgene | RRID: Addgene_64220 |
| pU6-sgRosa26-1_CBh-Cas9-T2A-BFP-P2A-Ad4E1B (pPD720) | Addgene | RRID: Addgene_64219 |

**Supplementary Table 12. Plasmids used in this study, previously described by our lab**

| Plasmid# | Plasmid name | Backbone | Leonard Lab Plasmid ID | Reference |
|---|---|---|---|---|
| **pPD436** | pBI-MCS-EYFP | pBI | L2881 | RRID: Addgene_58855 |
| **pHIE096** | CMB Bxb1 | pPD005 | L1571 | RRID: Addgene_139252 |
| **pPD005** | pcDNA Golden Gate | pcDNA3.1/HygroR(+) | L1267 | RRID: Addgene_138749 |
| **pPD1046** | TRE3GV mNeonGreen | TUPV1 | L1587 | RRID: Addgene_139268 |
| **pPD1049** | TRE3GV miRFP720 | TUPV1 | L1595 | RRID: Addgene_139276 |
| **pPD561** | CAG MCS | TUPV1 | L1494 | RRID: Addgene_139221 |
| **pPD562** | CAG MCS | TUPV2 | L1495 | RRID: Addgene_139222 |
| **pPD563** | CAG MCS | TUPV3 | L1496 | RRID: Addgene_139223 |
| **pPD564** | CAG MCS | TUPV4 | L1497 | RRID: Addgene_139224 |
| **pPD565** | CAG MCS | TUPV5 | L1498 | RRID: Addgene_139225 |
| **pPD566** | CAG MCS | TUPV6 | L1499 | RRID: Addgene_139226 |
| **pPD1151** | pLink1 | TUPV2 | L1560 | RRID: Addgene_139239 |
| **pPD1152** | pLink2 | TUPV3 | L1561 | RRID: Addgene_139240 |
| **pPD1153** | pLink3 | TUPV4 | L1562 | RRID: Addgene_139241 |
| **pPD1154** | pLink4 | TUPV5 | L1563 | RRID: Addgene_139242 |
| **pPD1155** | pLink5 | TUPV6 | L1564 | RRID: Addgene_139243 |

**Supplementary Table 13. Plasmids generated in this study**

| Plasmid# | Plasmid name and description | Backbone | Leonard Lab Plasmid ID | Reference[1] |
|---|---|---|---|---|
| | **B16F10-LP** | | | |
| **pPD783** | pU6-sgRosa26-1_CBh-Cas9-T2A-BFP-P2A-Ad4E4orf6 | pPD720 | L3114 | |
| **pPD864** | pLP Rosa26 EF1a attP EYFP-P2A-Hygro | | L3148 | |
| | **Backbones** | | | |
| **pPD1178** | Integration Vector (PuroR-P2A-miRFP720) | | L1599 | RRID: Addgene_139966 |
| **pPD1060** | pPD1060 Integration Vector (PuroR-P2A-DsRedExpress2) | | L1569 | |
| | **Transfected HBS (Figure 3b)** | | | |
| **pPD211** | HBS(CMV_min) EYFP | pPD005 | L2933 | |
| **pPD214** | HBS(YB_TATA) EYFP | pPD005 | L2936 | |
| **pPD217** | HBS(SV40_min) EYFP | pPD005 | L2939 | |
| | **Integration Vectors (Figures 1-4)** | | | |
| **pPD787** | *Basic HBS, CMV_min:* HBS(CMV_min) DsRE2; EBFP2/BlastR | pPD630 | L3118 | |
| **pPD788** | *Basic HBS, YB_TATA:* HBS(YB_TATA) DsRE2; EBFP2/BlastR | pPD630 | L3119 | |
| **pPD790** | *HBS, CMV_min, & dox-inducible HIF1α*:* HBS(CMV_min) DsRE2; Dox-inducible HIF1α*; EBFP2/BlastR | pPD630 | L3121 | |
| **pPD793** | *Control: dox-inducible DsRE2:* Dox-inducible DsRE2; EBP2/BlastR | pPD630 | L3124 | |
| | **Components for Integration Vectors (Figures 1-4)** | | | |
| **pPD766** | TRE3GV DsRE2 | TUPV1 | L3098 | |
| **pPD769** | TRE3GV HIF1α* | TUPV2 | L3100 | |
| **pPD770** | CAG TetOn3G | TUPV3 | L3101 | |

| | | | |
|---|---|---|---|
| **pPD847** | CAG TetOn3G | TUPV2 | L3132 |
| **pPD774** | HBS(CMV_min) DsRE2 | TUPV1 | L3105 |
| **pPD775** | HBS(YB_TATA) DsRE2 | TUPV1 | L3106 |
| **pPD852** | CAG EBFP2-2A-BlastR | TUPV2 | L3137 |
| **pPD853** | CAG EBFP2-2A-BlastR | TUPV3 | L3138 |
| **pPD854** | CAG EBFP2-2A-BlastR | TUPV4 | L3139 |
| **Integration Vectors (Figures 5-6)** | | | |
| **pMO18** | *Integration test vector:* Dox-Inducible mTagBFP2; miRFP670/BlastR; BB: PuroR | pD1060 | |
| **pPD1401** | *Control: only mTagBFP2:* mTagBFP2/AkaLuc/BlastR; BB: PuroR | pPD630 | L3465 |
| **pPD1402** | *Control: only dox-inducible mNeonGreen:* Dox-Inducible mNeonGreen; BlastR; BB: PuroR, DsRE2 | pPD630 | L3466 |
| **pPD1403** | *Control: only dox-inducible LumiScarlet:* Dox-Inducible LumiScarlet; BlastR; BB: PuroR | pPD630 | L3467 |
| **pPD1404** | *Control: only dox-inducible miRFP720:* Dox-Inducible miRFP720; BlastR; BB: PuroR | pPD630 | L3468 |
| **pPD1411** | *Basic HBS, SV40_min:* HBS(SV40_min) LumiScarlet; mTagBPF2/AkaLuc/BlastR; BB: miRFP720/PuroR | pPD1178 | L3469 |
| **pPD1412** | *Basic HBS, CMV_min* HBS(CMV_min) LumiScarlet; mTagBPF2/AkaLuc/BlastR; BB: miRFP720/PuroR | pPD1178 | L3470 |
| **pPD1413** | *Basic HBS, YB_TATA:* HBS(YB_TATA) LumiScarlet; mTagBPF2/AkaLuc/BlastR; BB: miRFP720/PuroR | pPD1178 | L3471 |
| **pPD1417** | *HBS – 3 copies:* [HBS(YB_TATA) LumiScarlet]x3; mTagBPF2/AkaLuc/BlastR; BB: miRFP720/PuroR | pPD1178 | L3475 |
| **pPD1418** | *HBS – 4 copies:* [HBS(YB_TATA) LumiScarlet]x4; mTagBPF2/AkaLuc/BlastR; BB: miRFP720/PuroR | pPD1178 | L3476 |
| **pPD1431** | *HBS – with mHIF1α feedback:* HBS(YB_TATA) LumiScarlet; HBS(YB_TATA) mHIF1α*; mTagBPF2/AkaLuc/BlastR; BB: miRFP720/PuroR | pPD1178 | L3481 |
| **pPD1432** | *HBS – with mHIF2α feedback:* HBS(YB_TATA) LumiScarlet; HBS(YB_TATA) mHIF2α; mTagBPF2/AkaLuc/BlastR; BB: miRFP720/PuroR | pPD1178 | L3482 |

| | | | |
|---|---|---|---|
| **pPD1433** | *HBS – with mHIF1α & HIF1β feedback:*<br>HBS(YB_TATA) LumiScarlet;<br>HBS(YB_TATA) mHIF1α*; HBS(YB_TATA) mHIF1β; mTagBPF2/AkaLuc/BlastR; BB: miRFP720/PuroR | pPD1178 | L3483 |
| **pPD1434** | *HBS – with mHIF2α & HIF1β feedback:*<br>HBS(YB_TATA) LumiScarlet<br>HBS(YB_TATA) mHIF2α; HBS(YB_TATA) mHIF1β; mTagBPF2/AkaLuc/BlastR; BB: miRFP720/PuroR | pPD1178 | L3484 |
| **Components for Integration Vectors (Figures 5-6)** | | | |
| **pPD1043** | TRE3GV mTagBFP2 | TUPV1 | L1584 |
| **pPD1031** | EF1alpha miRFP670-P2A-BlastR | TUPV2 | L1596 |
| **pPD1354** | Tre3GV LumiScarlet | TUPV1 | L3801 |
| **pPD1351** | HBS(CMV_min) LumiScarlet | TUPV1 | L3456 |
| **pPD1352** | HBS(SV40_min) LumiScarlet | TUPV1 | L3457 |
| **pPD1350** | HBS(YB_TATA) LumiScarlet | TUPV1 | L3455 |
| **pPD1357** | HBS(YB_TATA) LumiScarlet | TUPV2 | L3460 |
| **pPD1358** | HBS(YB_TATA) LumiScarlet | TUPV3 | L3461 |
| **pPD1359** | HBS(YB_TATA) LumiScarlet | TUPV4 | L3462 |
| **pPD1271** | HBS(YB_TATA) mHIF1α | TUPV2 | L3420 |
| **pPD1272** | HBS(YB_TATA) mHIF2α | TUPV2 | L3421 |
| **pPD1284** | HBS(YB_TATA) mHIF1β | TUPV3 | L3433 |
| **pPD1360** | CAG TetON3G | TUPV2 | L3463 |
| **pPD770** | CAG TetON3G | TUPV3 | L3101 |
| **pPD1286** | CAG TetON3G | TUPV4 | L3435 |
| **pPD1361** | CAG BlastR | TUPV3 | L3464 |
| **pPD1291** | CAG mTagBFP2-P2A-AkaLuc-2A-BlastR | TUPV1 | L3440 |
| **pPD1292** | CAG mTagBFP2-P2A-AkaLuc-2A-BlastR | TUPV2 | L3441 |
| **pPD1293** | CAG mTagBFP2-P2A-AkaLuc-2A-BlastR | TUPV3 | L3442 |
| **pPD1294** | CAG mTagBFP2-P2A-AkaLuc-2A-BlastR | TUPV4 | L3443 |
| **pPD1295** | CAG mTagBFP2-P2A-AkaLuc-2A-BlastR | TUPV5 | L3444 |

[1]Addgene numbers for new plasmids will be added prior to publication

**Supplementary Table 14. Cell lines obtained from external sources**

| Cell line | Source | Product ID/Reference |
|---|---|---|
| B16F10 | ATCC, VA, USA | CRL-6475, RRID: CVCL_0159 |
| HEK293FT | ThermoFisher/Life Technologies, MA, USA | R70007, RRID: CVCL_6911 |
| HEK293FT-LP | Ron Weiss, MIT | NAR (2) |
| TOP10 E. coli | ThermoFisher, MA, USA | C404010 |

**Supplementary Table 15. Cell lines generated in this study**

| Line# | Cell line name and description | Parental line(s) | Integrated Plasmid | Purity |
|---|---|---|---|---|
| **Figure 2B,C** | | | | |
| **PSD7** | *HBS, YB_TATA, unsorted:*<br>LP: HBS(YB_TATA) DsRE2; EBFP2/BlastR; PuroR | HEK293FT-LP | pPD788 | Selected[1] |
| **Figure 2D** | | | | |
| **PSD8** | *HBS, YB_TATA, Decile 1:*<br>LP: HBS(YB_TATA) DsRE2; EBFP2/BlastR; PuroR, Decile 1/10 | HEK293FT-LP, PSD7 | pPD788 | Sorted[2] |
| **PSD9** | *HBS, YB_TATA, Decile 2:*<br>LP: HBS(YB_TATA) DsRE2; EBFP2/BlastR; PuroR, Decile 2/10 | HEK293FT-LP, PSD7 | pPD788 | Sorted[2] |
| **PSD10** | *HBS, YB_TATA, Decile 3:*<br>LP: HBS(YB_TATA) DsRE2; EBFP2/BlastR; PuroR, Decile 3/10 | HEK293FT-LP, PSD7 | pPD788 | Sorted[2] |
| **PSD11** | *HBS, YB_TATA, Decile 4:*<br>LP: HBS(YB_TATA) DsRE2; EBFP2/BlastR; PuroR, Decile 4/10 | HEK293FT-LP, PSD7 | pPD788 | Sorted[2] |
| **PSD12** | *HBS, YB_TATA, Decile 5:*<br>LP: HBS(YB_TATA) DsRE2; EBFP2/BlastR; PuroR, Decile 5/10 | HEK293FT-LP, PSD7 | pPD788 | Sorted[2] |
| **PSD13** | *HBS, YB_TATA, Decile 6:*<br>LP: HBS(YB_TATA) DsRE2; EBFP2/BlastR; PuroR, Decile 6/10 | HEK293FT-LP, PSD7 | pPD788 | Sorted[2] |
| **PSD14** | *HBS, YB_TATA, Decile 7:*<br>LP: HBS(YB_TATA) DsRE2; EBFP2/BlastR; PuroR, Decile 7/10 | HEK293FT-LP, PSD7 | pPD788 | Sorted[2] |
| **PSD15** | *HBS, YB_TATA, Decile 8:*<br>LP: HBS(YB_TATA) DsRE2; EBFP2/BlastR; PuroR, Decile 8/10 | HEK293FT-LP, PSD7 | pPD788 | Sorted[2] |
| **PSD16** | *HBS, YB_TATA, Decile 9:*<br>LP: HBS(YB_TATA) DsRE2; EBFP2/BlastR; PuroR, Decile 9/10 | HEK293FT-LP, PSD7 | pPD788 | Sorted[2] |
| **PSD17** | *HBS, YB_TATA, Decile 10:*<br>LP: HBS(YB_TATA) DsRE2; EBFP2/BlastR; PuroR, Decile 10/10 | HEK293FT-LP, PSD7 | pPD788 | Sorted[2] |
| **Figures 3-4** | | | | |
| **PSD63** | *Basic HBS, CMV_min:*<br>HBS(CMV_min) DsRE2; EBFP2/BlastR | HEK293FT-LP, PSD7 | pPD787 | Sorted[3] |
| **PSD64** | *Basic HBS, YB_TATA:*<br>HBS(YB_TATA) DsRE2; EBFP2/BlastR | HEK293FT-LP, PSD7 | pPD788 | Sorted[3] |
| **PSD65** | *HBS, CMV_min, & dox-inducible HIF1α*:*<br>HBS(CMV_min) DsRE2; Dox-inducible HIF1α*; EBFP2/BlastR | HEK293FT-LP, PSD7 | pPD790 | Sorted[3] |
| **PSD66** | *Control: dox-inducible DsRE2:*<br>Dox-inducible DsRE2; EBP2/BlastR | HEK293FT-LP, PSD7 | pPD793 | Sorted[3] |
| **Supplementary Figures 9-10** | | | | |
| **PSD150** | B16F10-LP | B16F10 | pPD864 | Selected[4] |

| | | | | |
|---|---|---|---|---|
| **Figure 5B** | | | | |
| **JB009** | *Control: only mTagBFP2, unsorted:*<br>mTagBFP2/AkaLuc/BlastR; PuroR | B16F10-LP | pD1401 | Selected[5] |
| **JB010** | *Control: only dox-inducible mNeonGreen, unsorted:*<br>Dox-Inducible mNeonGreen; BlastR; PuroR | B16F10-LP | pD1402 | Selected[5] |
| **JB011** | *Control: only dox-inducible LumiScarlet, unsorted:*<br>Dox-Inducible LumiScarlet; BlastR; PuroR | B16F10-LP | pD1403 | Selected[5] |
| **JB012** | *Control: only dox-inducible miRFP720, unsorted:*<br>Dox-Inducible miRFP720; BlastR; PuroR | B16F10-LP | pD1404 | Selected[5] |
| **JB013** | *Basic HBS, SV40_min, unsorted:*<br>HBS(SV40_min) LumiScarlet; mTagBPF2/AkaLuc/BlastR; miRFP720/PuroR | B16F10-LP | pD1411 | Selected[5] |
| **JB014** | *Basic HBS, CMV_min, unsorted:*<br>HBS(CMV_min) LumiScarlet; mTagBPF2/AkaLuc/BlastR; miRFP720/PuroR | B16F10-LP | pD1412 | Selected[5] |
| **JB015** | *Basic HBS, YB_TATA, unsorted:*<br>HBS(YB_TATA) LumiScarlet; mTagBPF2/AkaLuc/BlastR; miRFP720/PuroR | B16F10-LP | pD1413 | Selected[5] |
| **JB019** | *HBS – 3 copies, unsorted:*<br>[HBS(YB_TATA) LumiScarlet]x3; mTagBPF2/AkaLuc/BlastR; miRFP720/PuroR | B16F10-LP | pD1417 | Selected[5] |
| **JB020** | *HBS – 4 copies, unsorted:*<br>[HBS(YB_TATA) LumiScarlet]x4; mTagBPF2/AkaLuc/BlastR; miRFP720/PuroR | B16F10-LP | pD1418 | Selected[5] |
| **JB025** | *HBS – with mHIF1α feedback, unsorted:*<br>HBS(YB_TATA) LumiScarlet; HBS(YB_TATA) mHIF1α*; mTagBPF2/AkaLuc/BlastR; miRFP720/PuroR | B16F10-LP | pPD1431 | Selected[5] |
| **JB026** | *HBS – with mHIF2α feedback, unsorted:*<br>HBS(YB_TATA) LumiScarlet; HBS(YB_TATA) mHIF2α; mTagBPF2/AkaLuc/BlastR; miRFP720/PuroR | B16F10-LP | pPD1432 | Selected[5] |
| **JB027** | *HBS – with mHIF1α & HIF1β feedback, unsorted:*<br>HBS(YB_TATA) LumiScarlet; HBS(YB_TATA) mHIF1α*; HBS(YB_TATA) mHIF1β; mTagBPF2/AkaLuc/BlastR; BB: miRFP720/PuroR | B16F10-LP | pPD1433 | Selected[5] |
| **JB028** | *HBS – with mHIF2α & HIF1β feedback, unsorted:*<br>HBS(YB_TATA) LumiScarlet HBS(YB_TATA) mHIF2α; HBS(YB_TATA) mHIF1β; mTagBPF2/AkaLuc/BlastR; BB: miRFP720/PuroR | B16F10-LP | pPD1434 | Selected[5] |
| **Figures 5C,D** | | | | |
| **JB051** | *Control: only mTagBFP2:*<br>mTagBFP2/AkaLuc/BlastR; PuroR | B16F10-LP, JB009 | pD1401 | Sorted[6] |
| **JB052** | *Control: only dox-inducible mNeonGreen:*<br>Dox-Inducible mNeonGreen; BlastR; PuroR | B16F10-LP, JB010 | pD1402 | Sorted[6] |
| **JB053** | *Control: only dox-inducible LumiScarlet:*<br>Dox-Inducible LumiScarlet; BlastR; PuroR | B16F10-LP, JB011 | pD1403 | Sorted[6] |
| **JB054** | *Control: only dox-inducible miRFP720:*<br>Dox-Inducible miRFP720; BlastR; PuroR | B16F10-LP, JB012 | pD1404 | [6] |
| **JB058** | *Basic HBS, YB_TATA:*<br>HBS(YB_TATA) LumiScarlet; mTagBPF2/AkaLuc/BlastR; miRFP720/PuroR | B16F10-LP, JB016 | pD1414 | Sorted[6] |

| | | | | |
|---|---|---|---|---|
| JB067 | *HBS – with mHIF1α feedback:*<br>HBS(YB_TATA) LumiScarlet; HBS(YB_TATA) mHIF1α*; mTagBPF2/AkaLuc/BlastR; miRFP720/PuroR | B16F10-LP, JB025 | pPD1431 | Sorted[6] |
| **JB068** | *HBS – with mHIF2α feedback:*<br>HBS(YB_TATA) LumiScarlet; HBS(YB_TATA) mHIF2α*; mTagBPF2/AkaLuc/BlastR; miRFP720/PuroR | B16F10-LP, JB026 | pPD1432 | Sorted[6] |

[1]With puromycin and then puromycin and blasticidin.

[2]From PDS7 into subpopulations based on the decile of EBFP2 fluorescence, as described for each line.

[3]From the parental line, which had been selected with puromycin and then puromycin and blasticidin, for the top 20% of EBFP2 expressing cells.

[4]With hygromycin.

[5]With puromycin and blasticidin.

[6]From the parental line, which had been selected with puromycin and blasticidin, for the top 20% of mTagBFP2 expressing cells.

**Supplementary Table 16. Model assumptions and rationale.**

| Species | Assumption | Justification |
|---|---|---|
| All | Transcription and translation can be represented as 1-step reactions in which components are not consumed | Simplest reasonable mechanism |
| HIF-1α and HIF-2α* (HIF-2α–$HAF_S$ complex) | HIF-1α and HIF-2α* can be used as proxies for HIF complexes; HIF-1β is in excess and binding is instantaneous and irreversible | Constitutive HIF-1β (3-9); α subunits will control dynamics |
| Reporter mRNA and HIF-1/2α mRNA (feedback topologies) | Transcription is linearly dependent on the sum of HIF-1α and HIF-2α*: $k_{txnBH}([HIF1\alpha_P] + [HIF2\alpha_P *])$ | Simplest reasonable mechanism |
| HIF-1α and HIF-2α | Multi-step degradations can be represented as 1-step reactions | Simplest reasonable mechanism |
| HIF-1α and HIF-2α | All $O_2$-dependent degradation can be represented as a single, 2nd order degradation: $-k_{dHP}pO_2[HIF1\alpha_P]$ | Not describing output dynamics in 21% $O_2$ |
| HIF-1α mRNA | Destabilization can be represented as a 2nd order degradation: $-k_{dH1R}[HIF1\alpha_R][aHIF1\alpha_R]$ | Destabilization decreases half-life of HIF-1α mRNA (10) |
| HIF-1α | HAF-dependent degradation can be represented as a single, 2nd order degradation dependent on the sum of the 2 HAF states: $-k_{dH1P}[HIF1\alpha_P]([HAF_P] + [HAF_S])$ | Simplest reasonable mechanism; HAF SUMOylation does not affect its degradation ability (11) |
| HAF | Experimentally observed short-term hypoxia degradation (6) can be represented as a piecewise, first-order protein degradation in 1% $O_2$: $k_{dHAF}(t) = \{k_{dHAF},\ t < t_{HAF}\ 0, \quad t \geq t_{HAF}$ | Unknown mode of regulation but mRNA levels not affected (12); simplest reasonable mechanism |
| HAF | SUMOylation can be represented as an irreversible, one step reaction | Simplest reasonable mechanism |
| HAF | SUMOylation rate is inversely proportional to oxygen pressure | Experimentally observed increase in SUMOylation at 1% $O_2$ vs. 20% $O_2$ (11) |

**Supplementary Table 17. ODEs and descriptions for Models D and D2 for the simple HBS.**

| State | Equation | Description | # |
|---|---|---|---|
| HAF mRNA | $\frac{d[HAF_R]}{dt} = k_{txn2} - k_{dR}[HAF_R]$ | transcription<br>basal degradation | 0 |
| HAF protein | $\frac{d[HAF_P]}{dt} = k_{tln}[HAF_R] - k_{dP}[HAF_P] - k_{dHAF}[HAF_P] - \frac{k_{bHS}}{pO_2}[HAF_P][SUMO_P]$ | translation<br>basal degradation<br>short-term hypoxic degradation<br>SUMOylation | 1 |
| SUMO mRNA | $\frac{d[SUMO_R]}{dt} = k_{txn} - k_{dR}[SUMO_R]$ | transcription<br>basal degradation | 2 |
| SUMO protein | $\frac{d[SUMO_P]}{dt} = k_{tln}[SUMO_R] - k_{dP}[SUMO_P] - \frac{k_{bHS}}{pO_2}[HAF_P][SUMO_P]$ | translation<br>basal degradation<br>SUMOylation | 3 |
| SUMO-ylated HAF | $\frac{d[HAF_S]}{dt} = \frac{k_{bHS}}{pO_2}[HAF_P][SUMO_P] - k_{dP}[HAF_S] - k_{bHH}[HIF2\alpha_P][HAF_S]$ | SUMOylation<br>basal degradation<br>HIF-2α– HAFs binding | 4 |
| Antisense HIF-1α RNA | $\frac{d[aHIF_R]}{dt} = k_{txnH}([HIF1\alpha_P] + [HIF2\alpha_P *]) - k_{dR}[aHIF_R]$ | transcription<br>basal degradation | 5 |
| HIF-1α mRNA | $\frac{d[HIF1\alpha_R]}{dt} = k_{txn} - k_{dR}[HIF1\alpha_R] - k_{dH1R}[aHIF1\alpha_R][HIF1\alpha_R]$ | transcription<br>basal degradation<br>antisense HIF-1α RNA destabilization | 6 |
| HIF-1α protein | $\frac{d[HIF1\alpha_P]}{dt} = k_{tln}[HIF1\alpha_R] - k_{dP}[HIF1\alpha_P] - k_{dHP}pO_2[HIF1\alpha_P] - k_{dH1P}[HIF1\alpha_P]([HAF_P] + [HAF_S])$ | translation<br>basal degradation<br>$O_2$-dependent degradation<br>HAF-dependent degradation | 7 |
| HIF-2α RNA | $\frac{d[HIF2\alpha_R]}{dt} = k_{txn} - k_{dR}[HIF2\alpha_R]$ | transcription<br>basal degradation | 8 |
| HIF-2α protein | $\frac{d[HIF2\alpha_P]}{dt} = k_{tln}[HIF2\alpha_R] - k_{dP}[HIF2\alpha_P] - k_{dHP}pO_2[HIF2\alpha_P] - k_{bHH}[HIF2\alpha_P][HAF_S]$ | translation<br>basal degradation<br>$O_2$-dependent degradation<br>HIF-2α– HAFs binding | 9 |
| HIF-2α–HAFs protein complex | $\frac{d[HIF2\alpha_P *]}{dt} = k_{bHH}[HIF2\alpha_P][HAF_S] - k_{dP}[HIF2\alpha_P *]$ | HIF-2α– HAFs binding<br>basal degradation | 10 |
| Reporter mRNA | $\frac{d[reporter_R]}{dt} = k_{txnBH}([HIF1\alpha_P] + [HIF2\alpha_P *]) - k_{dR}[reporter_R]$ | transcription<br>basal degradation | 11 |
| Reporter protein | $\frac{d[reporter_P]}{dt} = k_{tln}[reporter_R] - k_{dRep}[reporter_P]$ | translation<br>basal degradation | 12 |

[1]Models D and D2 have the same ODEs, which are shown above. The difference between the two models is whether $k_{txn2}$ if fixed (Model D) or free (Model D2).

[2]$pO_2$ = Oxygen pressure.

[3]Numbers correspond to the number used for each state in the model code.

**Supplementary Table 18. Parameter labels, descriptions, and fixed or calibrated values.**[1]

| Parameter | Free/fixed | Value and units | Description | Reference |
|---|---|---|---|---|
| $k_{txn}$ | Fixed | 1.0 U | Basal transcription rate for HIF and SUMO RNA species | (13) |
| $k_{dR}$ | Fixed | 2.7 $h^{-1}$ | Basal degradation rate for all RNA species | (13) |
| $k_{tln}$ | Fixed | 1.0 U | Basal translation rate for all protein species | (13) |
| $k_{dP}$ | Fixed | 0.35 $h^{-1}$ | Basal protein degradation rate (for all species except reporter) | (13) |
| $k_{dRep}$ | Fixed | 0.029 $h^{-1}$ | Basal reporter protein degradation rate | (13) |
| $t_{HAF}$ | Free | 43.43 h | Time at which HAF hypoxic degradation turns off | N/A |
| $k_{txn2}$ | Free | 24.17 U | HAF basal transcription rate | N/A |
| $k_{dHAF}$ | Free | 1.301 $h^{-1}$ | HAF hypoxic degradation rate | N/A |
| $k_{bHS}$ | Free | 7.394 mmHg*a.u.$^{-1}$*$h^{-1}$ | HAF SUMOylation rate | N/A |
| $k_{bHH}$ | Free | 0.3334 a.u.$^{-1}$*$h^{-1}$ | HIF-2α–SUMOylated HAF binding rate | N/A |
| $k_{txnH}$ | Free | 0.3012 U | HIF-dependent antisense HIF-1α RNA transcription rate | N/A |
| $k_{dH1R}$ | Free | 0.01279 a.u.$^{-1}$*$h^{-1}$ | antisense HIF-1α RNA-dependent HIF-1α mRNA degradation rate | N/A |
| $k_{dH1P}$ | Free | 0.1498 a.u.$^{-1}$*$h^{-1}$ | HAF-dependent HIF-1α protein degradation rate | N/A |
| $k_{dHP}$ | Free | 0.4490 mmHg$^{-1}$ $h^{-1}$ | $O_2$-dependent HIF-1/2α degradation rate | N/A |
| $k_{txnBH}$ | Free | 5.4055 U | HIF-dependent transcription at pHBS | N/A |

[1]U refers to an arbitrary transcription or translation unit as described previously (13,14). A.u. is arbitrary concentration unit. N/A means not applicable.

**Supplementary Table 19. ODEs and descriptions for Model A for the simple HBS.**

| State | Equation | Description | # |
|---|---|---|---|
| HAF mRNA | $\frac{d[HAF_R]}{dt} = k_{txn2} - k_{dR}[HAF_R]$ | transcription<br>basal degradation | 0 |
| HAF protein | $\frac{d[HAF_P]}{dt} = k_{tln}[HAF_R] - k_{dP}[HAF_P] - k_{bHH}[HIF2\alpha_P][HAF_P]$ | translation<br>basal degradation<br>HIF-2α– $HAF_P$ binding | 1 |
| Antisense HIF-1α RNA | $\frac{d[aHIF_R]}{dt} = k_{txnH}([HIF1\alpha_P] + [HIF2\alpha_P *]) - k_{dR}[aHIF_R]$ | transcription<br>basal degradation | 2 |
| HIF-1α mRNA | $\frac{d[HIF1\alpha_R]}{dt} = k_{txn} - k_{dR}[HIF1\alpha_R] - k_{dH1R}[aHIF1\alpha_R][HIF1\alpha_R]$ | transcription<br>basal degradation<br>antisense HIF-1α RNA destabilization | 3 |
| HIF-1α protein | $\frac{d[HIF1\alpha_P]}{dt} = k_{tln}[HIF1\alpha_R] - k_{dP}[HIF1\alpha_P] - k_{dHP}pO_2[HIF1\alpha_P] - k_{dH1P}[HIF1\alpha_P][HAF_P]$ | translation<br>basal degradation<br>$O_2$-dependent degradation<br>HAF-dependent degradation | 4 |
| HIF-2α RNA | $\frac{d[HIF2\alpha_R]}{dt} = k_{txn} - k_{dR}[HIF2\alpha_R]$ | transcription<br>basal degradation | 5 |
| HIF-2α protein | $\frac{d[HIF2\alpha_P]}{dt} = k_{tln}[HIF2\alpha_R] - k_{dP}[HIF2\alpha_P] - k_{dHP}pO_2[HIF2\alpha_P] - k_{bHH}[HIF2\alpha_P][HAF_P]$ | translation<br>basal degradation<br>$O_2$-dependent degradation<br>HIF-2α– $HAF_P$ binding | 6 |
| HIF-2α– $HAF_S$ protein complex | $\frac{d[HIF2\alpha_P *]}{dt} = k_{bHH}[HIF2\alpha_P][HAF_P] - k_{dP}[HIF2\alpha_P *]$ | HIF-2α– $HAF_P$ binding<br>basal degradation | 7 |
| Reporter mRNA | $\frac{d[reporter_R]}{dt} = k_{txnBH}([HIF1\alpha_P] + [HIF2\alpha_P *]) - k_{dR}[reporter_R]$ | transcription<br>basal degradation | 8 |
| Reporter protein | $\frac{d[DsRE2_P]}{dt} = k_{tln}[reporter_R] - k_{dRep}[reporter_P]$ | translation<br>basal degradation | 9 |

**Supplementary Table 20. ODEs and descriptions for Models B and B2 for the simple HBS.**[1]

| State | Equation | Description | # |
|---|---|---|---|
| HAF mRNA | $\frac{d[HAF_R]}{dt} = k_{txn2} - k_{dR}[HAF_R]$ | transcription<br>basal degradation | 0 |
| HAF protein | $\frac{d[HAF_P]}{dt} = k_{tln}[HAF_R] - k_{dP}[HAF_P] - k_{dHAF}[HAF_P] - k_{bHH}[HIF2\alpha_P][HAF_P]$ | translation<br>basal degradation<br>short-term hypoxic degradation<br>HIF-2α– $HAF_P$ binding | 1 |
| Antisense HIF-1α RNA | $\frac{d[aHIF_R]}{dt} = k_{txnH}([HIF1\alpha_P] + [HIF2\alpha_P *]) - k_{dR}[aHIF_R]$ | transcription<br>basal degradation | 2 |
| HIF-1α mRNA | $\frac{d[HIF1\alpha_R]}{dt} = k_{txn} - k_{dR}[HIF1\alpha_R] - k_{dH1R}[aHIF1\alpha_R][HIF1\alpha_R]$ | transcription<br>basal degradation<br>antisense HIF-1α RNA destabilization | 3 |
| HIF-1α protein | $\frac{d[HIF1\alpha_P]}{dt} = k_{tln}[HIF1\alpha_R] - k_{dP}[HIF1\alpha_P] - k_{dHP}pO_2[HIF1\alpha_P] - k_{dH1P}[HIF1\alpha_P][HAF_P]$ | translation<br>basal degradation<br>$O_2$-dependent degradation<br>HAF-dependent degradation | 4 |
| HIF-2α RNA | $\frac{d[HIF2\alpha_R]}{dt} = k_{txn} - k_{dR}[HIF2\alpha_R]$ | transcription<br>basal degradation | 5 |
| HIF-2α protein | $\frac{d[HIF2\alpha_P]}{dt} = k_{tln}[HIF2\alpha_R] - k_{dP}[HIF2\alpha_P] - k_{dHP}pO_2[HIF2\alpha_P] - k_{bHH}[HIF2\alpha_P][HAF_P]$ | translation<br>basal degradation<br>$O_2$-dependent degradation<br>HIF-2α– $HAF_P$ binding | 6 |
| HIF-2α– $HAF_S$ protein complex | $\frac{d[HIF2\alpha_P *]}{dt} = k_{bHH}[HIF2\alpha_P][HAF_P] - k_{dP}[HIF2\alpha_P *]$ | HIF-2α– $HAF_P$ binding<br>basal degradation | 7 |
| Reporter mRNA | $\frac{d[reporter_R]}{dt} = k_{txnBH}([HIF1\alpha_P] + [HIF2\alpha_P *]) - k_{dR}[reporter_R]$ | transcription<br>basal degradation | 8 |
| Reporter protein | $\frac{d[reporter_P]}{dt} = k_{tln}[reporter_R] - k_{dRep}[reporter_P]$ | translation<br>basal degradation | 9 |

[1]Models B and B2 have the same ODEs, which are shown above. The difference between the two models is whether $k_{txn2}$ if fixed (Model B) or free (Model B2).

**Supplementary Table 21. ODEs and descriptions for Model C for the simple HBS.**

| State | Equation | Description | # |
|---|---|---|---|
| HAF mRNA | $\frac{d[HAF_R]}{dt} = k_{txn2} - k_{dR}[HAF_R]$ | transcription<br>basal degradation | 0 |
| HAF protein | $\frac{d[HAF_P]}{dt} = k_{tln}[HAF_R] - k_{dP}[HAF_P] - \frac{k_{bHS}}{pO_2}[HAF_P][SUMO_P]$ | translation<br>basal degradation<br>SUMOylation | 1 |
| SUMO mRNA | $\frac{d[SUMO_R]}{dt} = k_{txn} - k_{dR}[SUMO_R]$ | transcription<br>basal degradation | 2 |
| SUMO protein | $\frac{d[SUMO_P]}{dt} = k_{tln}[SUMO_R] - k_{dP}[SUMO_P] - \frac{k_{bHS}}{pO_2}[HAF_P][SUMO_P]$ | translation<br>basal degradation<br>SUMOylation | 3 |
| SUMO-ylated HAF | $\frac{d[HAF_S]}{dt} = \frac{k_{bHS}}{pO_2}[HAF_P][SUMO_P] - k_{dP}[HAF_S] - k_{bHH}[HIF2\alpha_P][HAF_S]$ | SUMOylation<br>basal degradation<br>HIF-2α– HAF$_S$ binding | 4 |
| Antisense HIF-1α RNA | $\frac{d[aHIF_R]}{dt} = k_{txnH}([HIF1\alpha_P] + [HIF2\alpha_P *]) - k_{dR}[aHIF_R]$ | transcription<br>basal degradation | 5 |
| HIF-1α mRNA | $\frac{d[HIF1\alpha_R]}{dt} = k_{txn} - k_{dR}[HIF1\alpha_R] - k_{dH1R}[aHIF1\alpha_R][HIF1\alpha_R]$ | transcription<br>basal degradation<br>antisense HIF-1α RNA destabilization | 6 |
| HIF-1α protein | $\frac{d[HIF1\alpha_P]}{dt} = k_{tln}[HIF1\alpha_R] - k_{dP}[HIF1\alpha_P] - k_{dHP}pO_2[HIF1\alpha_P] - k_{dH1P}[HIF1\alpha_P]([HAF_P] + [HAF_S])$ | translation<br>basal degradation<br>O$_2$-dependent degradation<br>HAF-dependent degradation | 7 |
| HIF-2α RNA | $\frac{d[HIF2\alpha_R]}{dt} = k_{txn} - k_{dR}[HIF2\alpha_R]$ | transcription<br>basal degradation | 8 |
| HIF-2α protein | $\frac{d[HIF2\alpha_P]}{dt} = k_{tln}[HIF2\alpha_R] - k_{dP}[HIF2\alpha_P] - k_{dHP}pO_2[HIF2\alpha_P] - k_{bHH}[HIF2\alpha_P][HAF_S]$ | translation<br>basal degradation<br>O$_2$-dependent degradation<br>HIF-2α– HAF$_S$ binding | 9 |
| HIF-2α– HAF$_S$ protein complex | $\frac{d[HIF2\alpha_P *]}{dt} = k_{bHH}[HIF2\alpha_P][HAF_S] - k_{dP}[HIF2\alpha_P *]$ | HIF-2α– HAF$_S$ binding<br>basal degradation | 10 |
| Reporter mRNA | $\frac{d[reporter_R]}{dt} = k_{txnBH}([HIF1\alpha_P] + [HIF2\alpha_P *]) - k_{dR}[reporter_R]$ | transcription<br>basal degradation | 11 |
| Reporter protein | $\frac{d[reporter_P]}{dt} = k_{tln}[reporter_R] - k_{dRep}[reporter_P]$ | translation<br>basal degradation | 12 |

### SUPPLEMENTARY NOTES

#### Supplementary Note 1: Iterative model development and analysis

##### Model training data

The training data for each of the 3 HBS topologies consists of the baseline LumiScarlet reporter expression in MEPTRs at 21% $O_2$ and the 5-day expression time-course at 1% $O_2$ (days 0-4) (**Figure 5C**). Training data for the 3 HBS topologies were normalized to enable comparison to model simulations. Once converted from MFI to MEPTRs, the fluorescence value at each time point was divided by the mean fluorescence across days 0-4 of the simple HBS time series. The model simulations were normalized analogously; the simulated reporter protein concentration at each time point was divided by the mean simulated reporter protein concentration across days 0-4 of the simple HBS time series.

##### Simulation of hypoxia dynamics

Model simulations were run in a manner that recapitulated experimental conditions. The model equations for each HBS topology were solved in two sequential simulations, the first in 21% $O_2$ (normoxia) and the second in 1% $O_2$ (hypoxia). See **Supplementary Note 2** for normoxic and hypoxic oxygen pressure calculations. First, normoxia simulations were run with an initial condition of 0 arbitrary concentration units for each state, for 500 h to ensure that all model states reached a baseline steady state; the final time point concentration for each state from this simulation was used to initialize hypoxic simulations. Thus, the steady state reached under normoxic concentrations represents the baseline reporter expression of the cells at 21% $O_2$, before they are placed in the hypoxic incubator. The hypoxia simulations were run for either 96 h, and the reporter expression was defined as the simulated reporter concentration at each 24 h interval (corresponding to days 0-4 in the training data). Custom Python 3.8 scripts based on GAMES v2.0 (1) and the SciPy Python package (15) odeint ODE solver were used to solve model equations, with default error tolerance of $1.49012*10^{-8}$ a.u. (arbitrary concentration unit) for each model state. Visualizations of reporter dynamics only show the second simulation in 1% $O_2$, where the 0 h simulated reporter expression is the same as the baseline 21% $O_2$ reporter expression (**Figure 6C**).

##### Parameter estimation

*Cost function*

To quantify the agreement between the normalized simulation values and the normalized training data values for a particular parameter set, $\theta$, we used a $\chi^2$ cost function. Calibrated parameter sets are defined by having the lowest cost function value.

$$\chi^2(\theta) = \sum_{i=1}^{n} \left(y_i{}^{exp} - y_i^{sim}(\theta)\right)^2 \qquad \text{(S1)}$$

Here, $n$ is the total number of datapoints in the training data set, $y_i{}^{exp}$ is the i$^{th}$ datapoint in the training data, and $y_i^{sim}(\theta)$ is the simulated value of the i$^{th}$ datapoint using the parameter set $\theta$.

*Coefficient of determination*

We also used the coefficient of determination ($R^2$) to quantify the correlation between the normalized simulation values and the normalized training data values for a particular parameter set, $\theta$. $R^2$ was not directly utilized in the optimization algorithm, but it served as an additional, more interpretable, quantitative metric compared to $\chi^2$ for evaluation of optimized parameter sets (1). $R^2$ values fall between 0 and 1, with an $R^2 = 1$ indicating perfect correlation between the training data and simulation values.

$$R^2(\theta) = 1 - \frac{\sum_{i=1}^{n}(y_i{}^{exp} - y_i^{sim}(\theta))^2}{\sum_{i=1}^{n}(\bar{y}^{exp} - y_i^{sim}(\theta))^2} \qquad \text{(S2)}$$

Here, $n$ is the total number of datapoints in the training data set, $y_i{}^{exp}$ is the i$^{th}$ datapoint in the training data, $y_i^{sim}(\theta)$ is the simulated value of the i$^{th}$ datapoint using the parameter set $\theta$, and $\bar{y}^{exp}$ is the mean of the training data set.

*Parameter estimation method and analysis of calibrated parameter sets*
Parameters were estimated using a multi-start Levenberg-Marquardt optimization algorithm for least squares curve fitting, implemented with the LMFIT Python package (16) and described by Dray et al (1). We generated initial guesses using a Latin-Hypercube (17) global search across reasonable parameter bounds (implemented with the SALib Python package (18)), with $n_{search}$= 1000 parameter sets. We selected $n_{init}$= 100 parameter sets with the lowest $\chi^2$ to serve as initial guesses for optimization.

Calibrated parameter sets were analyzed based on qualitative and quantitative criteria. First, we visually inspected the model fit to the training data and time series of internal model states to ensure that trajectories were physically plausible. Next, we evaluated whether the calibrated parameter set yielded a model fit that satisfied our modeling objectives. Finally, we used the $\chi^2$ and $R^2$ values of the model fit to evaluate the quantitative agreement between the simulated values and training data values.

*Parallelization of computational tasks*
PEM evaluation and PEM runs were parallelized using 8 cores (chosen based on the number of cores available in the hardware used to run the simulations) to improve computational efficiency. Parallelization was implemented using scripts from GAMES v2.0 (1) and Python's multiprocessing package.

**Supplementary Note 2: Oxygen pressure calculations for model simulations**
We calculated the oxygen pressure in normoxia and hypoxia for model simulations using partial pressure calculations described in (19). First, we calculated the atmospheric pressure at Evanston's altitude relative to atmospheric pressure at sea level using an equation defined in (19):

$$P_a = P_0 * e^{-0.127a} \qquad \text{(S3)}$$

Here, $P_a$ is the atmospheric pressure at altitude a (in km), and $P_0$ is atmospheric pressure at sea level (101325 Pa). Evanston's altitude is 0.185 km, resulting in an atmospheric pressure of 98972 Pa or 742.4 mmHg. Next, we calculated the partial pressure of oxygen in the total atmospheric pressure. We subtracted the water pressure at 37°C and 100% humidity (47 mmHg), and the pressure of 5% $CO_2$ (37.1 mmHg)—all typical incubator conditions of cultured cells (19)—from the total atmospheric pressure. We then calculated the partial pressure of $O_2$ in normoxia (20.9% $O_2$) and hypoxia (1% $O_2$) from the remaining air pressure (658.2 mmHg). In normoxia, $pO_2$ = 138 mmHg, and in hypoxia, $pO_2$ = 6.6 mmHg. These oxygen pressures were substituted for $pO_2$ in model equations in normoxia and hypoxia simulations, respectively.

**Supplementary Note 3: Model ODEs for HBS topologies with feedback**
For the HIF feedback topologies, the only change to the model ODEs is in the mRNA equation for the HIF with feedback. An additional transcription term is added to describe the HIF-dependent transcription of the respective HIF, the same term used for transcription of reporter mRNA. For example, the HIF-1 mRNA equation in the HIF-1 Feedback HBS:

$$\begin{aligned}\frac{d[HIF1\alpha_R]}{dt} &= k_{txn} + k_{txnBH}([HIF1\alpha_P] + [HIF2\alpha_P *]) \\ &\quad -k_{dR}[HIF1\alpha_R] - k_{dH1R}[aHIF1\alpha_R][HIF1\alpha_R]\end{aligned} \qquad \text{(S4)}$$

**Supplementary Note 4: Parameter identifiability**
An important consideration and limitation of the HBS model is that there is more than one set of parameters that result in similar agreement to the experimental data, which means some parameters are not identifiable, i.e., able to be uniquely estimated given the model equations and the training data (1). Although the choice of parameter values among these parameter sets does not affect the fit to the training data, it is possible that this choice will affect

predictions made using the model. We chose not to perform parameter identifiability analysis and refinement like that in the GAMES paper (1) because making predictions is beyond the scope of this work; however, we used an alternative method to assess parameter identifiability to gain intuition that will guide future modeling work. We performed a parameter sensitivity analysis determine which parameters had the greatest impact on the overall fit to the training data. We independently varied each parameter by $\pm$ 10% of the calibrated value and calculated the $\chi^2$ between the simulation values and the training data for each parameter variation. The sensitivity of $\chi^2$ to each parameter was quantified via the percent change in $\chi^2$ relative to the $\chi^2$ for the calibrated parameter set. The results revealed that $\chi^2$ was insensitive to some parameters, particularly, $k_{txnH}$ and $k_{dH1R}$, indicating that these parameters are likely not well-constrained (**Supplementary Figure 16**). Future work employing the HBS model to make predictions will require a more rigorous parameter identifiability analysis (e.g., with the parameter profile likelihood approach (20)) and subsequent model reduction or experimental design, depending on the nature of the unidentifiable parameters and the modeling objectives.

**Supplementary Note 5: One-way ANOVAs**

Below are the outcomes from one-way ANOVAs and Tukey's HSD tests. Null hypotheses were that there existed no effects of sample preparation on the measured reporter expression, evaluated separately for each fluorescent protein.

Comparison of methodologies for fluorophore oxidation in **Figure 1C** and **Supplementary Figure 3** (non-fixed samples)

DsRed-Express2:

- Sample preparation $p = 5.87\times10^{-12}$
    - All sample preparation conditions resulted in significantly different measured reporter expression compared to the samples cultured under normoxia, and all had $p << 0.001$ except for the following preparation conditions:
        - Cultured in hypoxia and kept on ice with a pre-harvest oxidation step ($p = 0.0064$)
        - Cultured in hypoxia and kept at 37ºC with a post-harvest oxidation step ($p = 0.0025$)
    - For the samples that were oxidized post-harvest at various temperatures, the following differences were significant (all $p < 0.05$):
        - Ice vs. room temperature
        - Ice vs. 37ºC
        - 4ºC vs. room temperature
        - 4ºC vs. 37ºC

mTagBFP2:

- Sample preparation $p = 8.63\times10^{-6}$
    - No sample preparation conditions resulted in significantly different measured reporter expression compared to the samples cultured under normoxia (all $p > 0.05$) except for those cultured in hypoxia and kept on ice without an oxidation step ($p = 0.0002$) or at room temperature without an oxidation step ($p = 0.0049$)
    - For the samples that were oxidized post-harvest at various temperatures, no differences were significantly different (all $p > 0.05$)

mNeonGreen:

- Sample preparation $p = 6.98\times10^{-7}$
    - The following sample preparation conditions resulted in significantly different measured reporter expression compared to the samples cultured under normoxia:
        - Cultured in hypoxia and kept on ice) or at room temperature without an oxidation step (both $p << 0.001$)
        - Cultured in hypoxia and kept on ice with a post-harvest oxidation step ($p = 0.046$)
    - For the samples that were oxidized post-harvest at various temperatures, no differences were significantly different (all $p > 0.05$)

Comparison of methodologies for fluorophore oxidation in **Figure 1C** and **Supplementary Figure 3** (fixed samples)
DsRed-Express2:

- Sample preparation $p = 9.92 \times 10^{-7}$
  - All sample preparation conditions resulted in significantly different measured reporter expression compared to the samples cultured under normoxia (all $p < 0.01$)
  - For the samples that were oxidized post-harvest at various temperatures, no sample preparation conditions were significantly different except for the Ice vs. 37ºC conditions ($p = 0.038$)

mTagBFP2:

- Sample preparation $p > 0.05$

mNeonGreen:

- Sample preparation $p = 0.025$
  - No sample preparation conditions resulted in significantly different measured reporter expression compared to the samples cultured under normoxia (all $p > 0.05$) except for those cultured in hypoxia and kept at room temperature ($p = 0.037$)
  - For the samples that were oxidized post-harvest at various temperatures, no differences were significantly different (all $p > 0.05$)

Below are the outcomes from one-way ANOVA and Tukey's HSD tests. Null hypotheses were that there existed no effects of level of constitutive EBFP2 expression (expression bin) on measured reporter expression fold induction for cobalt treated vs. untreated samples.

Comparison of fold inductions in each constitutive EBFP2 expression bin in **Figure 2G**

- Constitutive EBFP2 expression bin $p = 0.043$
  - No bins had significantly different reporter expression fold induction (all $p > 0.05$) except for bin 1 vs. bin 10 ($p = 0.035$)

**Supplementary Note 6: Two-way ANOVAs**
Below are the outcomes from two-way ANOVAs and Tukey's HSD tests. Null hypotheses were that there existed no effects of level of constitutive EBFP2 expression (expression bin), cobalt treatment, or their interaction on the measured reporter expression.

Comparison of untreated vs. cobalt treated samples in each constitutive EBFP2 expression bin in **Figure 2C**

- Constitutive EBFP2 expression bin $p = 3.09 \times 10^{-18}$
  - Bins 1 vs. 10 and 2 vs. 10 were significantly different (both $p < 0.05$)
- Cobalt treatment $p = 8.14 \times 10^{-27}$
- Interaction between constitutive EBFP2 expression bin and cobalt treatment $p = 6.62 \times 10^{-16}$
  - All bins had significantly different reporter expression upon cobalt treatment (all $p < 0.05$) except for bin 1 ($p = 1.0$), bin 2 ($p = 0.98$), and bin 3 ($p = 0.23$)
  - For cobalt treated samples in each bin, all differences were statistically significant (all $p < 0.05$) except for:
    - Bin 1 vs. bin 2
    - Bin 1 vs. bin 3
    - Bin 2 vs. bin 3
    - Bin 2 vs. bin 4
    - Bin 3 vs. bin 4
    - Bin 3 vs. bin 5
    - Bin 3 vs. bin 6
    - Bin 4 vs. bin 5
    - Bin 4 vs. bin 6
    - Bin 4 vs. unsorted

      - Bin 5 vs. bin 6
      - Bin 5 vs. bin 7
      - Bin 5 vs. unsorted
      - Bin 6 vs. bin 7
      - Bin 6 vs. bin 8
      - Bin 6 vs. unsorted
      - Bin 7 vs. bin 8
      - Bin 7 vs. unsorted
      - Bin 8 vs. bin 9
      - Bin 8 vs. unsorted
    - For untreated samples in each bin, no differences were statistically different (all $p > 0.05$)

Below are the outcomes from two-way ANOVAs and Tukey's HSD tests. Null hypotheses were that there existed no effects of level of constitutive EBFP2 expression at the time of sorting (sorted or unsorted population), cobalt treatment, or their interaction on the measured reporter expression.

Comparison of untreated vs. cobalt treated samples in each constitutive EBFP2 expression sorted population in **Figure 2D**

- Constitutive EBFP2 expression sorted population $p = 1.07\times10^{-29}$
    - No sorted populations are significantly different (all $p > 0.05$)
- Cobalt treatment $p = 1.32\times10^{-56}$
- Interaction between constitutive EBFP2 expression sorted population and cobalt treatment $p = 1.85\times10^{-29}$
    - All sorted populations had significantly different reporter expression upon cobalt treatment (all $p << 0.01$)
    - For cobalt treated samples in each sorted population, all differences were statistically significant except for:
      - Unsorted population vs. population 4
      - Unsorted population vs. population 5
      - Unsorted population vs. population 6
      - Population 2 vs. population 3
      - Population 4 vs. population 5
      - Population 4 vs. population 6
      - Population 5 vs. population 6
      - Population 8 vs. population 9
      - Population 9 vs. population 10

Below are the outcomes from two-way ANOVAs and Tukey's HSD tests. Null hypotheses were that there existed no effects of minimal promoter, cobalt treatment, or their interaction on the measured reporter expression.

Comparison of untreated vs. cobalt treated samples for each minimal promoter in **Figure 3B**

HEK293FT (**Figure 3B left):**

- Minimal promoter $p = 1.37\times10^{-6}$
    - No minimal promoters were significantly different (all $p > 0.05$)
- Cobalt treatment $p = 2.64\times10^{-10}$
- Interaction between minimal promoter and cobalt treatment $p = 1.59\times10^{-7}$
    - The CMV and YB_TATA minimal promoters each had significantly different reporter expression upon cobalt treatment (both $p << 0.001$)
    - For cobalt treated samples for each minimal promoter, all differences were statistically significant (all $p < 0.001$)

B16F10 (**Figure 3B right):**

- Minimal promoter $p = 3.85\times10^{-7}$
  - No minimal promoters were significantly different (all $p > 0.05$)
- Cobalt treatment $p = 4.58\times10^{-11}$
- Interaction between minimal promoter and cobalt treatment $p = 3.39\times10^{-6}$
  - Each minimal promoter had significantly different reporter expression upon cobalt treatment (all $p < 0.01$)
  - For cobalt treated samples for each minimal promoter, the following differences were statistically significant (both $p << 0.001$):
    - CMV vs. YB_TATA
    - SV40 vs. YB_TATA

Below are the outcomes from two-way ANOVAs and Tukey's HSD tests. Null hypotheses were that there existed no effects of minimal promoter, sample treatment, or their interaction on the measured reporter expression.

Comparison of normoxia vs. hypoxia treated samples for each minimal promoter in **Figure 5B** (SV40, CMV, and YB_TATA)

- Minimal promoter $p = 2.93\times10^{-7}$
  - No minimal promoters were significantly different (all $p > 0.05$)
- Treatment $p = 9.03\times10^{-10}$
- Interaction between minimal promoter and treatment $p = 0.0251$
  - Each minimal promoter had significantly different reporter expression for normoxia vs. hypoxia treatment (all $p << 0.001$)
  - For hypoxia treated samples for each minimal promoter, the following differences were statistically significant (both $p << 0.001$):
    - CMV vs. SV40
    - SV40 vs. YB_TATA

Below are the outcomes from two-way ANOVAs and Tukey's HSD tests. Null hypotheses were that there existed no effects of HBS copy number, sample treatment, or their interaction on the measured reporter expression.

Comparison of normoxia vs. hypoxia treated samples for each HBS copy number in **Figure 5B** (YB_TATA, YB_TATA + 2 productive HBS copies, and YB_TATA + 3 productive HBS copies)

- HBS copy number $p = 7.68\times10^{-8}$
  - No copy numbers were significantly different (all $p > 0.05$)
- Treatment $p = 6.70\times10^{-11}$
- Interaction between copy number and treatment $p = 7.26\times10^{-8}$
  - Each copy number had significantly different reporter expression for normoxia vs. hypoxia treatment (all $p < 0.001$)
  - For hypoxia treated samples for each copy number, the following differences were statistically significant (both $p << 0.001$):
    - YB_TATA vs. YB_TATA + 2 productive HBS copies
    - YB_TATA vs. YB_TATA + 3 productive HBS copies

Below are the outcomes from two-way ANOVAs and Tukey's HSD tests. Null hypotheses were that there existed no effects of HBS topology, sample treatment, or their interaction on the measured reporter expression.

Comparison of normoxia vs. hypoxia treated samples for each HBS topology in **Figure 5B** (no feedback, HIF1α feedback, HIF2α feedback, HIF1α and HIF1β feedback, and HIF2α and HIF1β feedback)

- HBS topology $p = 6.02\times10^{-16}$
  - No topologies were significantly different (all $p > 0.05$)
- Treatment $p = 5.63\times10^{-27}$
- Interaction between topology and treatment $p = 1.52\times10^{-15}$

- Each topology had significantly different reporter expression for normoxia vs. hypoxia treatment (all $p << 0.001$)
- For hypoxia treated samples for each topology, the following differences were statistically significant (all $p << 0.001$):
  - No feedback topology vs. any other topology
  - HIF1α feedback topology vs. any other feedback topology

**Supplementary Note 7: Two-way repeated measures ANOVAs**
Below are the outcomes from two-way repeated measures ANOVAs and Tukey's HSD tests. Null hypotheses were that there existed no effects of number of days of treatment, sample treatment, or their interaction on the measured reporter expression.

Comparison of normoxia, normoxia and doxycycline, hypoxia, and hypoxia and doxycycline treated samples in over 4 days in **Figure 1F**

- Day $p = 4.67\times10^{-34}$
- Sample treatment $p = 1.27\times10^{-11}$
  - All sample treatments were statistically different ($p << 0.001$) except for
    - Hypoxia vs. normoxia ($p = 1$)
    - Hypoxia and doxycycline vs. normoxia and doxycycline ($p = 0.99$)
- Interaction between day sample treatment $p = 4.18\times10^{-31}$
  - For each day of treatment, no days were statistically different for the normoxia and doxycycline vs. hypoxia and doxycycline conditions (all $p > 0.05$)
  - For each day of treatment, no days were statistically different for the normoxia vs. hypoxia conditions (all $p > 0.05$)
  - For each day of treatment, the normoxia vs. normoxia and doxycycline conditions were statistically different (all $p << 0.001$) except for day 0 ($p = 1$)
  - For each day of treatment, the hypoxia vs. hypoxia and doxycycline conditions were statistically different (all $p << 0.001$) except for day 0 ($p = 1$)

Comparison of normoxia, normoxia and cobalt, and hypoxia treated samples in over 4 days in **Figure 3C**

YB_TATA (**Figure 3C left)**:

- Day $p = 7.79\times10^{-28}$
- Sample treatment $p = 7.00\times10^{-9}$
  - Hypoxia vs. normoxia and hypoxia vs. normoxia and cobalt were statistically different (both $p << 0.001$)
- Interaction between day sample treatment $p = 1.13\times10^{-28}$
  - For each day of treatment, the hypoxia vs. normoxia conditions were statistically different ($p << 0.001$) except for day 0 ($p = 1$)
  - For each day of treatment, the hypoxia vs. normoxia and cobalt conditions were statistically different ($p << 0.001$) except for day 0 ($p = 1$)
  - For hypoxia treated conditions on each day, all differences were statistically different (all $p << 0.001$)

CMV (**Figure 3C right**):

- Day $p = 6.37\times10^{-25}$
- Sample treatment $p = 5.04\times10^{-9}$
  - Hypoxia vs. normoxia and hypoxia vs. normoxia and cobalt were statistically different (both $p << 0.001$)
- Interaction between day sample treatment $p = 5.94\times10^{-24}$

    - For each day of treatment, the hypoxia vs. normoxia conditions were statistically different ($p$ << 0.001) except for day 0 ($p = 1$)
    - For each day of treatment, the hypoxia vs. normoxia and cobalt conditions were statistically different ($p$ << 0.001) except for day 0 ($p = 1$)
    - For hypoxia treated conditions on each day, all differences were statistically different ($p$ << 0.001)

Comparison of normoxia, normoxia and cobalt, and physoxia treated samples in over 4 days in **Figure 3D**

YB_TATA (**Figure 3D left)**:

- Day $p = 2.50\times10^{-10}$
- Sample treatment $p = 4.35\times10^{-4}$
    - Physoxia vs. normoxia were statistically different ($p = 2.0\times10^{-4}$)
    - Normoxia vs. normoxia and cobalt were statistically different ($p = 0.043$)
- Interaction between day sample treatment $p = 8.73\times10^{-9}$
    - For each day of treatment, the following days were statistically different for the normoxia vs. normoxia and cobalt conditions (all other $p > 0.05$):
        - Day 2 ($p = 5.0\times10^{-4}$)
    - For each day of treatment, the following days were statistically different for the normoxia vs. physoxia conditions (all other $p > 0.05$):
        - Day 2 ($p$ << 0.001)
        - Day 3 ($p = 2.3\times10^{-3}$)
        - Day 4 ($p$ << 0.001)
    - For each day of treatment, the following days were statistically different for the normoxia and cobalt vs. physoxia conditions (all other $p > 0.05$):
        - Day 2 ($p = 1.4\times10^{-3}$)
        - Day 4 ($p$ << 0.001)
    - For physoxia treated conditions on each day, all differences were statistically different (all $p < 0.05$) except for day 0 vs. day 1 and day 2 vs. day 4 (both $p = 1$)

CMV (**Figure 3D right**):

- Day $p = 2.07\times10^{-14}$
- Sample treatment $p = 1.86\times10^{-5}$
    - Physoxia vs. normoxia were statistically different ($p$ << 0.001)
    - Normoxia vs. normoxia and cobalt were statistically different ($p = 9.0\times10^{-4}$)
- Interaction between day sample treatment $p = 1.27\times10^{-11}$
    - For each day of treatment, the following days were statistically different for the normoxia vs. normoxia and cobalt conditions (all other $p > 0.05$):
        - Day 1 ($p$ << 0.001)
        - Day 2 ($p$ << 0.001)
        - Day 3 ($p$ << 0.001)
        - Day 4 ($p = 7.0\times10^{-4}$)
    - For each day of treatment, the following days were statistically different for the normoxia vs. physoxia conditions (all other $p > 0.05$):
        - Day 2 ($p$ << 0.001)
        - Day 3 ($p$ << 0.001)
        - Day 4 ($p$ << 0.001)
    - For each day of treatment, the following days were statistically different for the normoxia and cobalt vs. physoxia conditions (all other $p > 0.05$):
        - Day 1 ($p = 0.017$)
        - Day 3 ($p$ << 0.001)
        - Day 4 ($p$ << 0.001)

- For physoxia treated conditions on each day, all differences were statistically different (all $p < 0.05$) except for:
  - Day 0 vs. day 1 ($p = 0.43$)
  - Day 2 vs. day 3 ($p = 0.75$)
  - Day 2 vs. day 4 ($p = 1$)
  - Day 3 vs. day 4 ($p = 0.43$)

Comparison of normoxia, normoxia and cobalt, normoxia and doxycycline, and hypoxia treated samples in over 4 days in **Figure 4B**

- Day $p = 6.99\times10^{-45}$
- Sample treatment $p = 8.45\times10^{-16}$
  - Hypoxia vs. normoxia and doxycycline, normoxia vs. normoxia and doxycycline, and normoxia and cobalt vs. normoxia and doxycycline were statistically different (all $p << 0.001$)
- Interaction between day sample treatment $p = 2.24\times10^{-44}$
  - For each day of treatment, the normoxia and doxycycline vs. hypoxia conditions were statistically different ($p << 0.001$) except for day 0 ($p = 1$)
  - For each day of treatment, the normoxia and doxycycline vs. normoxia and cobalt conditions were statistically different ($p << 0.001$) except for day 0 ($p = 1$)
  - For each day of treatment, the normoxia and doxycycline vs. normoxia conditions were statistically different ($p << 0.001$) except for day 0 ($p = 1$)
  - For each day of treatment, the hypoxia vs. normoxia conditions were statistically different ($p << 0.001$) except for day 0 ($p = 1$)
  - For each day of treatment, the hypoxia vs. normoxia and cobalt conditions were statistically different ($p = 0.012$ for day 1; all other $p << 0.001$) except for day 0 ($p = 1$)
  - For normoxia and doxycycline treated conditions on each day, all differences were statistically different ($p < 0.001$)
  - For hypoxia treated conditions on each day, all differences were statistically different ($p < 0.001$)

Below are the outcomes from two-way repeated measures ANOVAs and Tukey's HSD tests. Null hypotheses were that there existed no effects of number of days of treatment, HBS topology, or their interaction on the measured reporter expression, evaluated for hypoxia treatment only.

Comparison of hypoxia (1% $O_2$) treated samples in over 4 days in **Figure 5C**

- Day $p = 7.121.00\times10^{-17}$
- HBS topology $p = 6.56\times10^{-5}$
  - No topologies were statistically different (all $p > 0.05$)
- Interaction between day and HBS topology $p = 5.82\times10^{-5}$
  - For each day of treatment, the following days were statistically different for the no feedback HBS vs. HIF1α feedback HBS (all other $p > 0.05$)
    - Day 1 ($p = 0.004$)
    - Day 2 ($p << 0.001$)
    - Day 3 ($p = 0.011$)
  - For each day of treatment, the following days were statistically different for the no feedback HBS vs. HIF2α feedback HBS (all other $p > 0.05$)
    - Day 2 ($p << 0.001$)
    - Day 3 ($p = 0.0006$)
  - For each day of treatment, the following days were statistically different for the HIF1α feedback HBS vs. HIF2α feedback HBS (all other $p > 0.05$):
    - Day 1 ($p = 0.023$)
  - For the HIF1α feedback HBS on each day, no differences were statistically significant (all $p > 0.05$) except for

        - Comparisons between day 0 and any other day (all $p << 0.001$)
        - Day 1 vs. day 2 ($p = 0.001$)
        - Day 1 vs. day 3 ($p = 0.041$)
    - For the HIF2α feedback HBS on each day, the following differences were statistically significant (all other $p > 0.05$)
        - )
        - Comparisons between day 0 and any other day (all $p << 0.001$)
        - Comparisons between day 1 and day 2, 3, or 4 (all $p << 0.001$)
    - For the no feedback HBS on each day, the following differences were statistically significant (all other $p > 0.05$)
        - Day 0 vs. day 1 ($p = 0.027$)
        - Comparisons between day 0 and day 2, 3, or 4 (all $p << 0.001$)
        - Comparisons between day 1 and day 3 or4 (both $p < 0.05$)

Comparison of physoxia (5% $O_2$) treated samples in over 4 days in **Figure 5D**

- Day $p = 1.51\times10^{-13}$
- HBS topology $p = 2.50\times10^{-5}$
    - The HIF1α feedback HBS vs. no feedback HBS were statistically different ($p = 0.003$)
    - The HIF1α feedback HBS vs. HIF2α feedback HBS were statistically different ($p < 0.001$)
- Interaction between day and HBS topology $p = 2.95\times10^{-6}$
    - For each day of treatment, all days were statistically different for the no feedback HBS vs. HIF1α feedback HBS (all $p < 0.01$) except for day 0 ($p = 0.96$)
    - For each day of treatment, no days were statistically different for the no feedback HBS vs. HIF2α feedback HBS (all $p > 0.05$) except for day 4 ($p << 0.001$)
    - For each day of treatment, all days were statistically different for the HIF1α feedback HBS vs. HIF2α feedback HBS (all $p < 0.001$) except for day 0 ($p = 0.95$) and day 2 ($p = 0.088$)
    - For the HIF1α feedback HBS on each day, all differences were statistically significant (all $p < 0.05$) except for
        - Day 1 vs. day 3 ($p = 0.55$)
        - Day 2 vs. day 3 ($p = 0.87$)
    - For the HIF2α feedback HBS on each day, the following differences were statistically significant (all other $p > 0.05$)
        - Comparisons between day 0 and either day 1 or day 4 (all $p < 0.05$)
    - For the no feedback HBS on each day, the following differences were statistically significant (all other $p > 0.05$)
        - Comparisons between day 4 and any other day (all $p << 0.001$)

**Supplementary Note 8: two-tailed Welch's *t*-tests**

Below are the outcomes two-tailed Welch's *t*-tests followed by BH procedure. Null hypotheses were that there existed no effect of cobalt dose on measured reporter expression.

Cobalt dose response in **Figure 2B**

- 0 μM $CoCl_2$ vs. 3 μM $CoCl_2$ $p = 0.35$ (not significant, n.s.)
- 0 μM $CoCl_2$ vs. 15 μM $CoCl_2$ $p = 0.40$ (n.s.)
- 0 μM $CoCl_2$ vs. 30 μM $CoCl_2$ $p = 0.33$ (n.s.)
- 0 μM $CoCl_2$ vs. 150 μM $CoCl_2$ $p = 0.025$
- 0 μM $CoCl_2$ vs. 300 μM $CoCl_2$ $p = 0.0077$
- 0 μM $CoCl_2$ vs. 600 μM $CoCl_2$ $p = 0.0077$

Below are the outcomes two-tailed Welch's *t*-tests followed by BH procedure. Null hypotheses were that there existed no effect of number of days of treatment in hypoxia on measured reporter expression vs. normoxia baseline, evaluated for each HBS topology simultaneously.

Comparison of hypoxia (1% $O_2$) treated samples over 4 days vs. normoxia baseline in **Figure 5C**
No feedback HBS:

- Normoxia baseline vs. hypoxia day 0 $p = 0.016$
- Normoxia baseline vs. hypoxia day 1 $p = 0.0046$
- Normoxia baseline vs. hypoxia day 2 $p = 0.0052$
- Normoxia baseline vs. hypoxia day 3 $p = 0.0022$
- Normoxia baseline vs. hypoxia day 4 $p = 0.00010$

HIF1α feedback HBS:

- Normoxia baseline vs. hypoxia day 0 $p = 0.064$ (not significant, n.s.)
- Normoxia baseline vs. hypoxia day 1 $p = 0.011$
- Normoxia baseline vs. hypoxia day 2 $p = 0.0051$
- Normoxia baseline vs. hypoxia day 3 $p = 0.023$
- Normoxia baseline vs. hypoxia day 4 $p = 0.0052$

HIF2α feedback HBS:

- Normoxia baseline vs. hypoxia day 0 $p = 0.027$
- Normoxia baseline vs. hypoxia day 1 $p = 0.00037$
- Normoxia baseline vs. hypoxia day 2 $p = 0.011$
- Normoxia baseline vs. hypoxia day 3 $p = 0.0031$
- Normoxia baseline vs. hypoxia day 4 $p = 0.0051$

Below are the outcomes two-tailed Welch's *t*-tests followed by BH procedure. Null hypotheses were that there existed no effect of number of days of treatment in physoxia on measured reporter expression vs. normoxia baseline, evaluated for each HBS topology simultaneously.

Comparison of physoxia (5% $O_2$) treated samples over 4 days vs. normoxia baseline in **Figure 5D**
No feedback HBS:

- Normoxia baseline vs. hypoxia day 0 $p = 0.97$ (not significant, n.s.)
- Normoxia baseline vs. hypoxia day 1 $p = 2.20\times10^{-4}$
- Normoxia baseline vs. hypoxia day 2 $p = 2.68\times10^{-3}$
- Normoxia baseline vs. hypoxia day 3 $p = 2.20\times10^{-4}$
- Normoxia baseline vs. hypoxia day 4 $p = 0.034$

HIF1α feedback HBS:

- Normoxia baseline vs. hypoxia day 0 $p = 0.19$ (n.s.)
- Normoxia baseline vs. hypoxia day 1 $p = 2.20\times10^{-4}$
- Normoxia baseline vs. hypoxia day 2 $p = 1.37\times10^{-3}$
- Normoxia baseline vs. hypoxia day 3 $p = 4.01\times10^{-4}$
- Normoxia baseline vs. hypoxia day 4 $p = 0.017$

HIF2α feedback HBS:

- Normoxia baseline vs. hypoxia day 0 $p = 0.19$ (n.s.)
- Normoxia baseline vs. hypoxia day 1 $p = 5.94\times10^{-3}$
- Normoxia baseline vs. hypoxia day 2 $p = 1.31\times10^{-3}$
- Normoxia baseline vs. hypoxia day 3 $p = 4.01\times10^{-4}$
- Normoxia baseline vs. hypoxia day 4 $p = 9.04\times10^{-3}$
